# Long-range regulatory effects of Neandertal DNA in modern humans

**DOI:** 10.1101/2021.10.20.464628

**Authors:** Danat Yermakovich, Vasili Pankratov, Urmo Võsa, Bayazit Yunusbayev, Estonian Biobank Research Team, Michael Dannemann

## Abstract

The admixture between modern humans and Neandertals has resulted in ∼2% of the genomes of present-day non-Africans being composed of Neandertal DNA. Association studies have shown that introgressed DNA significantly influences phenotypic variation in people today and that several of the phenotype-associated archaic variants had links to expression regulation as well. In general, introgressed DNA has been demonstrated to significantly affect the transcriptomic landscape in people today. However, little is known about how much of that impact is mediated through long-range regulatory effects that have been shown to explain ∼20% of expression variation.

Here we identified 60 transcription factors (TFs) with their top cis-eQTL SNP being of Neandertal ancestry in GTEx and predicted long-range Neandertal DNA-induced regulatory effects by screening for the predicted target genes of those TFs. We show that genes in regions devoid of Neandertal DNA are enriched among the target genes of some of these TFs. Furthermore, archaic cis-eQTLs for these TFs included multiple candidates for local adaptation and have associations with various immune traits, schizophrenia, blood cell type composition and anthropometric measures. Finally, we show that our results can be replicated in empirical trans-eQTLs with Neandertal variants.

Our results suggest that the regulatory reach of Neandertal DNA goes beyond the 40% of genomic sequence that it still covers in present-day non-Africans and that via this mechanism Neandertal DNA additionally influences the phenotypic variation in people today.

## Introduction

The genome sequences of archaic humans such as Neandertals and Denisovans have shown that modern humans admixed with both archaic groups ∼55,000 years ago (Meyer et al., 2012; Prüfer et al., 2017; Vernot et al., 2016). As a result of these admixtures ∼2% of the genomes of present-day non-Africans are composed of Neandertal DNA and an additional 2-5% of the genomes of Oceanians are derived from Denisovans (Sankararaman et al., 2016). Several studies have used information from genome-wide association studies to link archaic DNA to their potential phenotypic effects. These studies have shown that Neandertal DNA significantly influences skin and hair traits, immunity and behavioral phenotypes (Dannemann and Kelso, 2017; McArthur et al., 2021; Quach et al., 2016; Simonti et al., 2016). Many of the phenotype-associated archaic variants have been associated with transcriptional regulatory effects as well, including several instances of Neandertal DNA with additional evidence for local adaptation (Gittelman et al., 2016). In general it has been demonstrated that both Neandertal and Denisovan DNA have significantly shaped the transcriptomic landscape in people today (Dannemann et al., 2017; Gokhman et al., 2020; McCoy et al., 2017; Quach et al., 2016; Silvert et al., 2019; Vespasiani et al., n.d.). The analyses of the regulatory impact of introgressed archaic DNA have been based on cis-eQTLs, allele-specific expression, methylation data and presence of archaic variants in regulatory sequence motifs. At the same time it has been demonstrated that regulatory sequences in modern human genomes also have been subject to negative selection against archaic DNA (Petr et al., 2019; Telis et al., 2020). Both observations indicate that one major mechanism through which archaic DNA is influencing modern human phenotype variation is by its impact on gene expression regulation. These observations are perhaps not surprising given that the heritability of many phenotypes including disease is often mediated by gene expression (Yao et al., 2020).

However, previous studies have also shown that among eQTLs, cis-eQTLs only explain ∼6% of gene expression variation (Ouwens et al., 2020). In contrast, trans-eQTLs, i.e. long-distance regulatory effects, have been shown to explain with ∼20% a substantially larger proportion of gene expression differences. The underlying mechanisms of the long-range impact of trans-eQTLs include transcription factor (TF) activation and chromatin-chromatin interactions that can modulate the expression of large sets of genes across the genome (Marbach et al., 2016; Rao et al., 2014). Given the polygenic nature of many complex traits and the growing evidence that many of those traits are regulated by complex gene expression networks, trans-eQTLs and their ability to impact many genes simultaneously might provide an important mechanism in understanding the molecular bases of polygenic phenotypes (Boyle et al., 2017).

Large consortia like GTEx have revolutionized our understanding of the genetic basis of gene expression regulation (Consortium and The GTEx Consortium, 2020). Thousands of cis-eQTLs have been annotated in a diverse set of 49 tissues. However, while the current sample sizes of up to 700 donors per tissue provide sufficient power to robustly annotate cis-eQTLs only 163 trans-eQTLs were detected in GTEx across all tissues combined. This discrepancy between the number of cis and trans-eQTLs can be explained by both the smaller effect sizes of trans-eQTLs compared to cis-eQTLs and the substantially larger number of variant-gene pairs tested in a genome-wide trans-eQTL screen, that also require higher statistical significance thresholds to account for multiple testing (Ouwens et al., 2020). Recently, the eQTLGen consortium has published blood expression data from 31,684 samples, a dataset with a substantially increased association power compared to other existing datasets like GTEx (Võsa et al., 2021). The authors provide a genome-wide cis-eQTL maps and a trans-eQTL screen for 10,317 trait-associated variants. The study also has provided new insight into the regulatory mechanism of trans-eQTLs and has shown that ∼47% of significant trans-eQTLs could be linked to direct and indirect transcription factor (TF) activity.

Here we used this information provided by the eQTLGen study to scan the GTEx cohort for Neandertal-DNA-associated cis-eQTLs that are linked to expression levels of TF genes and used this information to predict potential trans-eQTL effects. Using TF target gene information we then explored the potential genomic range of the predicted trans-eQTLs that are associated with Neandertal DNA, with a particular focus on regions that are devoid of Neandertal DNA. In order to understand which groups of phenotypes are potentially affected by predicted Neandertal trans-eQTL variants we explored available GWAS results for association between the underlying variants of those predicted trans-eQTLs and disease and non-disease traits. In addition, we tested if our candidate trans-eQTLs showed signals of positive selection in present-day human populations. Finally, in order to explore the robustness of these phenotypic and evolutionary inferences for our predicted trans-eQTLs we compared our results to empirical Neandertal trans-eQTLs in eQTLGen.

## Results

### Trans-eQTLs in GTEx and eQTLGen

We first explored the trans-eQTL-associated variants in GTEx and eQTLGen for SNPs that are of likely Neandertal ancestry. We will refer to these SNPs as aSNPs (archaic SNPs) throughout the manuscript. Such aSNPs have previously been annotated in 1,000 Genomes Project individuals based on (i) their allele-sharing patterns of variants that are absent in the African Yoruba population - a population with the lowest levels of inferred Neandertal ancestry in the 1,000 Genomes cohort - and found in the high coverage genomes of three archaic humans (two Neandertals and one Denisovan) as well as non-Africans and (ii) their link to a haplotype that is not compatible with incomplete lineage sorting, another genomic phenomenon that can lead to a similar sharing pattern (Materials and methods).

Among the 163 trans-eQTLs in GTEx we didn’t find any aSNPs. However, the limited number of annotated trans-eQTLs in GTEx makes it difficult to evaluate the significance of this observation. Among the 3,853 significant trans-eQTLs (out of 10,317 tested SNPs) reported by the eQTLGen study (Võsa et al., 2021)] with a cohort size which surpasses GTEx in its statistical power, we found 18 to be aSNPs. Among these 18 Neandertal-linked trans-eQTLs were 3 pairs of SNPs, showing high levels of linkage disequilibrium among them (r^2^>0.5 between the two aSNPs for all three pairs, Materials and methods). We have collapsed these three pairs, reducing the numbers of independent Neandertal trans-eQTL associations to 15 (Table S1). Ten of these trans-eQTLs were associated with a significantly altered expression of one target gene, while the five other trans-eQTLs were linked to expression changes of multiple genes with 2, 3, 13, 27 and 34 target genes respectively. Notably, the archaic alleles for the two trans-eQTLs with 27 and 34 target genes were associated with directional impacts on expression (Table S1). In the presence of the archaic allele at the given loci, 67% (18/27, P=0.12, Binomial Test) of genes associated with *rs72973711* subjected to an expression-increasing effect, while 78% (28/34, P=2.0×10^−4^, Binomial Test) of genes related to *rs13098911/rs13063635* region show lower expression levels.

We compared the proportion of significant Neandertal trans-eQTLs to the fraction in 100 same size random sets of frequency-matched non-archaic variants in the pool of 10,317 eQTLGen trans-eQTL SNPs. The number of significant trans-eQTLs in the non-archaic sets were between 12 and 24 (95% confidence interval), which was comparable to the number of significant aSNPs (P=0.58) (Materials and methods). However, it’s not possible to assess how both the absence of aSNPs in GTEx trans-eQTLs and the comparable numbers of aSNP and frequency-matched non-archaic variants in eQTLGen trans-eQTLs translate to the genome-wide long-range regulatory impact of Neandertal DNA.

### Prediction of Neandertal-linked trans-eQTL effects

The limited number of SNPs tested for trans-eQTL effects in eQTLGen and the reduced power to test for trans-eQTLs in smaller datasets like GTEx prevents us from directly associating Neandertal variants with long-range regulatory effects on a genome-wide scale. However, the observation of the involvement of TFs to initiate and facilitate long-range regulatory effects allows us to use this information to predict potential genomic regulatory reach of Neandertal DNA. In this study, we focussed on one particular mechanism: the effect of Neandertal variants on nearby TFs expression and on the predicted targets of the affected TF. Overall, up to 9% of the trans-eQTL effects observed by Võsa & Claringbould et al. can be linked to this particular mechanism, with another 38% involving TFs at some other stage in the regulatory chain. In order to identify TFs that show evidence for a regulatory link to nearby Neandertal DNA we scanned the GTEx dataset for SNPs that are the top cis-eQTL for TFs in a given tissue (such genes are from here on referred to as “cTF”). Across 49 diverse tissues we found a total of 60 TFs for which the top cis-eQTL SNP was an aSNP (90 unique aSNPs; and 60 cTFs we refer to as “a-cTF”) in at least one tissue (FDR<0.05, Table S2, Methods). Most of these a-cTFs were found to be significantly regulated in only one (42 a-cTFs) or two (10 a-cTFs) tissues. Conversely, five a-cTFs were found in more than 6 tissues. *ZNF143* and *ZNF189* showed significant aSNP cis-eQTLs in 17 and 16 tissues respectively, including diverse sets of tissues such as the heart, arteries, blood, skin, adipose tissues, intestine and the brain (Table S2). For both genes we observed a consistently lower expression in the presence of the archaic allele. Among the other three a-cTFs with cis-eQTL in multiple tissues was *STAT2* with a total of seven tissues with aSNP cis-eQTLs. The affected tissues with such regulatory effects included two adipose tissues, liver, skin, nerve, cerebellar and artery tissues, with five of these tissues showing a lower expression of *STAT2* in the presence of the archaic alleles. The cis-eQTL aSNP assosiations with *STAT2* were linked to an archaic haplotype that also showed trans-eQTL associations in eQTLGen (*rs2066807/rs2066819*). One of the two trans-eQTL genes associated with the archaic variants was *IFI16* - a predicted target of *STAT2* (ENCODE). In eQTLGen, *STAT2* showed significantly lower expression in the presence of the archaic alleles (P=3.3×10^−122^, Z-Score=-23.5, Table S1). *STAT2* has been shown to function as a transcription activator. The lower expression of *IFI16* in the presence of the archaic alleles therefore suggests that a direct regulation by *STAT2* is consistent with these observations. Furthermore, among the 18 significant aSNP-associated trans-eQTLs in eQTLGen the aSNPs *rs2066807/rs2066819* were the only case where the reconstructed regulatory mechanism of the trans-eQTL was via a direct TF regulation (Võsa et al., 2021). The aSNP for a second trans-eQTL (*rs12908161*) was in high LD (r^2^>0.5) with the a-cTF *ZNF592* and therefore a second candidate for a direct transcription factor regulatory mechanism. However, none of the TF target gene databases provided target gene information for *ZNF592*, preventing us from testing whether trans-eQTL genes for *rs12908161* are predicted targets of this a-cTF.

In general, at least one significantly a-cTF was found in 40 of the 49 tissues. The tissue with the most active TFs was skeletal muscle, lung, adipose (subcutaneous), skin, thyroid (7 a-cTFs in each tissue). Across all tissues, 65.4% of a-cTFs showed a higher expression in the presence of the archaic allele (P=4.8×10^−4^, Binomial test). This expression bias was tentatively more pronounced in brain tissues (85.0% vs. 61.9% in non-brain tissues, P=0.07, Fisher’s exact test).

Twenty-nine of the 60 GTEx a-cTFs also showed significant cis-eQTL associations with the same aSNPs in eQTLGen (FDR<0.05). The correlation of effect sizes between GTEx and eQTLGen for the 29 overlapping GTEx a-cTFs aSNPs was high (Spearman’s ρ=0.66, P=2.1×10^−13^) and highly consistent (79% showed the same expression direction). The eQTLGen data also included data from GTEx blood expression data. When removing GTEx blood a-cTFs from the analysis the correlation remained comparable (Spearman’s ρ=0.67, P=2.9×10^−13^, 79% shared expression direction), suggesting that the correlation is not primarily driven by the partially shared data in eQTLGen and GTEx.

These results suggest that a-cTFs in GTEx are reproducible in other cohorts and provide support for the robustness of the cTFs for our analyses. In addition, finding almost half of the a-cTFs in the blood tissue from eQTLGen suggests that the higher statistical power in eQTLGen allows it to detect more subtle regulatory effects that are more pronounced in non-blood tissues in GTEx and hence are not detected in GTEx blood samples due to the smaller cohort.

### Interaction between Neandertal-linked TFs

Next, we sought to investigate whether the 60 a-cTFs potentially affect shared biological pathways and therefore tested the level of interaction between them and whether they are enriched in particular functional pathways (Materials and methods). We first generated a protein-protein interaction network between all 60 a-cTFs using the STRING interaction database (Szklarczyk et al., 2019). STRING provides the protein-protein interaction information based on different resources, such as experimentally validated interaction or curated databases, predictions algorithms, protein homology, co-expression information and text mining. All but five a-cTFs were predicted to interact with one or more other a-cTF via at least one of these types of interactions (Figure 1A). The five a-cTFs with the largest numbers of interactions showed between 15 and 21 links to other cTFs - higher levels than all other a-cTFs (13 or less). The gene with the most interactions was *JUN* with 21 links to other proteins (no other a-cTFs showed more than 16). In order to put the level of connectivity between Neandertal-linked a-cTFs in perspective we compared it to the connectivity levels of 100 randomly sampled sets of 60 non-archaic cTFs. We additionally sampled these background cTF sets in a way that they match the tissue prevalence of the a-cTFs. A protein-protein interaction can be supported by up to eight different types of resources. We considered both: (i) an interaction supported by at least one resource and (ii) the total number of resources that support an interaction. We found that for both analyses the connectivity for a-cTFs (for i and ii) was larger compared to the background sets. The average number of interactions between a-cTFs was 1.8-fold (for (i), P<0.01) and 1.9-fold (for (ii), P=0.02) higher compared to the background cTF sets (Figure 1B). However, the numbers of interactions for the top a-cTFs (for (i) *JUN* with 21 and for (ii) *KAT2B* with 42) were not statistically different from the interactions of the top cTFs in the background sets (for (i) P=0.54; for (ii) P=0.74), suggesting that the elevated connectivity levels for a-cTFs was not primarily driven by the a-cTFs with the most interactions.

**Figure 1:**
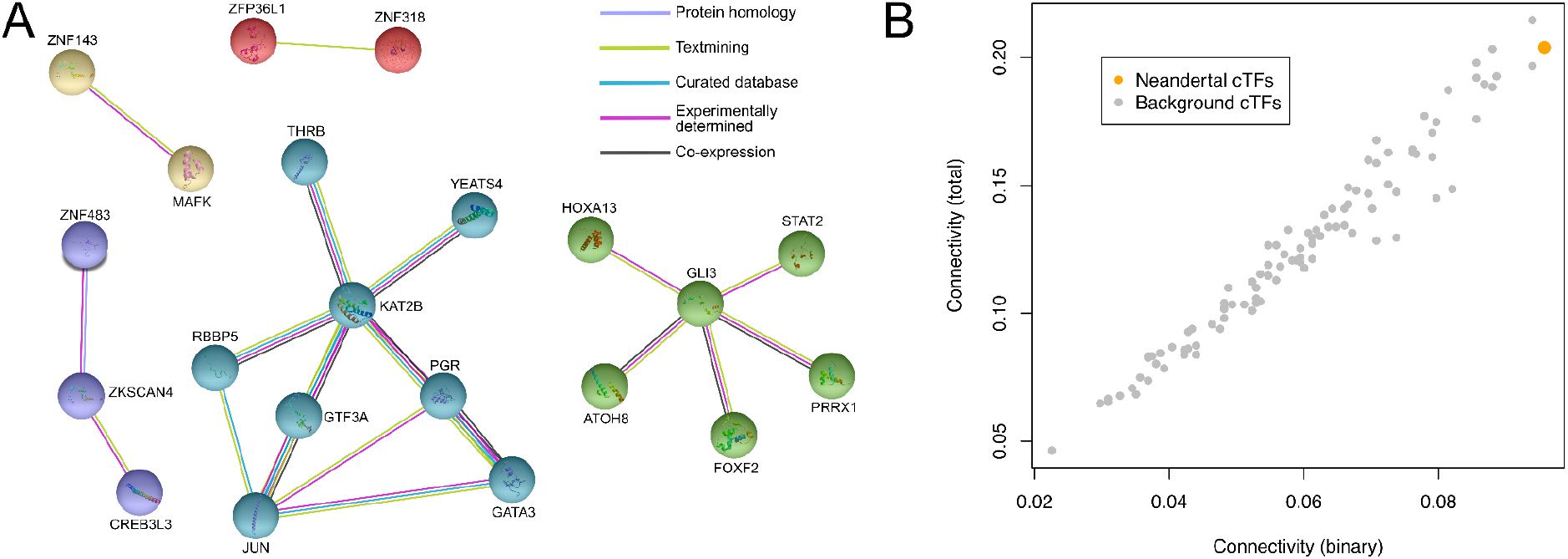
Protein-protein interaction network of Neandertal a-cTFs. **(A)** Protein-protein interaction network for Neandertal a-cTFs generated with STRING. Only a-cTFs that show at least one connection to another a-cTF with a medium connection confidence of 0.4 are displayed. Interaction types are colored according to the legend that is illustrated on the top right hand side of the panel. **(B)** The mean connectivity of the Neandertal network (orange) and 100 randomly sampled and tissue-prevalence-matched non-Neandertal cTF networks (gray) are displayed. The x-axis measures connectivity based on the presence of any type of interaction between cTFs while the y-axis considers the total number of types of interactions between cTFs.

Next, we tested whether the set of 60 a-cTFs shows any functional enrichment in the Gene Ontology (GO), KEGG or Reactome (Material and methods). We first tested the 60 a-cTFs compared against a background set of all cTFs in GTEx. We didn’t find any significantly enriched GO category or pathway at FDR<0.05. The absence of an enrichment result might be explained by the reduced power of this analysis that is based on 60 genes and a background of ∼1,000 genes. However, when we tested for enrichment compared to all protein-coding genes we found 18 GO categories and one Reactome pathway enriched (Table S3). Not unexpectedly, ten of the GO categories and the one enriched REACTOME pathway were related to transcription factor related processes. However, we also found other enriched categories that were related to circadian rhythm, hippocampus development, steroid receptor activity, and several additional developmental processes (artery development, cell fate determination, gland morphogenesis and leucocyte differentiation).

Taken together, our results suggest that we found elevated levels of interactivity between a-cTFs and that they could be involved in regulating several developmental processes.

#### The regulatory reach of predicted trans-eQTLs

In order to understand the potential reach of a-cTFs we next analyzed their predicted target genes. Target gene prediction information across seven prediction databases was available for 27 of the 60 GTEx a-cTFs (Table S4). However, prediction information varied widely between databases (Table S5). For example, while the 27 GTEx a-cTFs were present in at least one of the seven prediction databases, a-cTFs in individual databases ranged from just one (MotifMap) to 18 (Human TF-TG). Consequently, also the number of predicted target genes for these sets of a-cTFs showed substantial differences between databases and ranged between 189 (MotifMap) and 17,086 (ENCODE) for all a-cTFs combined.

Interestingly, some of the predicted target genes were located in genomic regions that have previously been reported to be devoid of Neandertal and Denisovan ancestry (Sankararaman et al., 2016; Vernot et al., 2016). Among the 257 protein-coding genes located in the five autosomal deserts reported by both studies 248 were within the predicted target gene sets for all TFs combined. Almost all of those genes were predicted to be regulated by at least one of the 27 GTEx a-cTFs (244/248 desert genes) (Table S4). We also found one empirical archaic trans-eQTL in eQTLGen (trans-eQTL aSNP *rs4805834*) for which the affected gene *TCEA1* is located within one of the autosomal deserts on chromosome 8 (54.5-65.4MB). In addition we found that *UTP14A*, which was linked to trans-eQTL aSNP *rs12603526*, was also located on one of the deserts that were reported on chromosome X (Figure 2).

**Figure 2:**
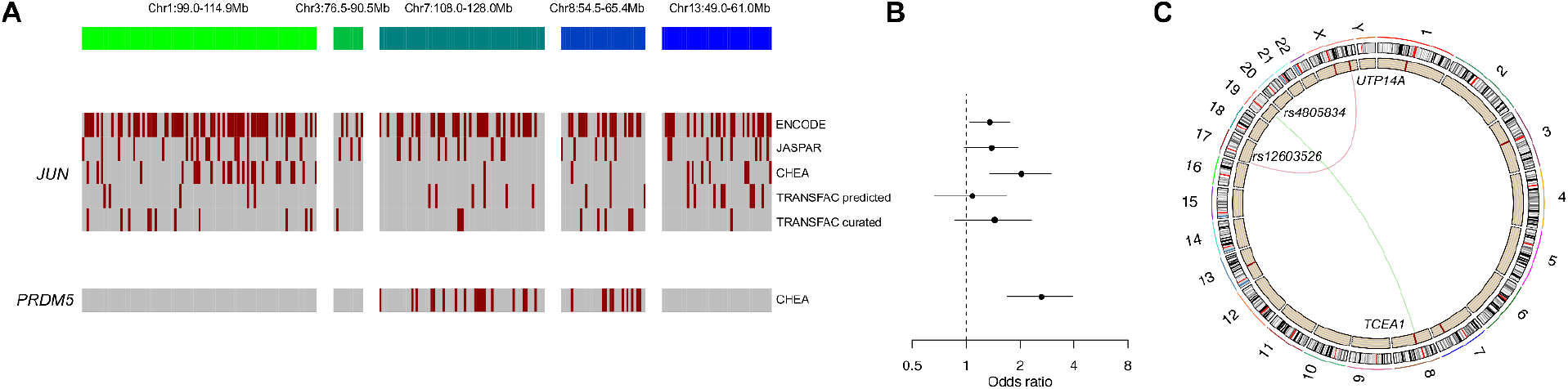
The regulatory Impact of Neandertal DNA on archaic deserts. **(A)** Predicted TF target genes for two a-cTFs (*JUN, and PRDM5*) with an over-proportional overlap with genes within autosomal archaic deserts. Each tile represents a desert gene (aligned in chromosomal order) and predicted targets for a given database are colored in red. **(B)** Odds ratios with 95% confidence intervals representing the proportional overlap between genes in archaic deserts and target genes for a given a-cTF and prediction database, as displayed in panel A are shown. **(C)** Circo plot showing the connection between genomic coordinates for two Neandertal trans-eQTL aSNPs and their predicted desert target genes. Archaic deserts are highlighted in the inner band in red.

Our results suggest that genes in deserts are not out of regulatory reach of Neandertal DNA. However, we were seeking to further assess the significance of the role of Neandertal DNA on desert genes. For this purpose we tested for TFs with predicted target genes that were over-represented in archaic deserts. We again leveraged information from the seven TF target prediction database to screen for such instances. If prediction information for a TF was available for more than one database we required all odds ratios to be larger than 1 and at least two enrichment P values (Fisher’s exact test) to be smaller than 0.05, at least one of which remaining at that significance after multiple testing correction. TFs with prediction information in only one database were required to have a more restricted FDR threshold of 10^−4^. We found 17 of the 1,693 GTEx cTFs passed that criterion. Interestingly, this list included *JUN and PRDM5*, two a-cTFs (*JUN*: Brain cortex; *PRDM5*: Frontal cortex BA9 and spleen). And while predicted target genes for *JUN* were evenly distributed across all five deserts, targets for *PRDM5* showed substantial differences in their prevalence between individual deserts (Figure 2A-B). All 28 *PRDM5* desert target genes were found on chromosomes 7 (18 of 68) and 8 (10 of 35). The desert region on chromosome 7 is the largest of all autosomal archaic deserts. Two genes within the desert on chromosome 7 that have been associated with modern human-specific biology are *FOXP2* and *ROBO2*. Both genes are among the targets of a-cTFs (*FOXP2* predicted target of *FOXC1, FOXF2, GATA3, HOXA13, JUN, MAFK, RBBP5, THRB, ZNF143*; *ROBO2* predicted target of *GATA3, JUN, LMX1A, MAFK, RBBP5, THRB, ZNF143*) (Maricic et al., 2013; Peyrégne et al., 2017). The desert on chromosome 1 also encompasses five amylase genes, a gene group that has previously been reported to be shaped by human-specific evolutionary processes (Hardy et al., 2015; Inchley et al., 2016; Perry et al., 2007). Four of those gene are also predicted to be regulated by a-cTFs (*AMY1B* and *AMY1C* [both predicted targets of *FOXC1, JUN*]; *AMY2A* [*JUN, PGR*]; *AMY2B* [*FOXC1, HOXA13, JUN, PGR*]).

Hence, our results suggest that genes in archaic deserts, including several candidates for modern human specific biology, might nevertheless be influenced by introgressed archaic DNA. However it remains unclear how many of the predicted target genes are in fact regulated by these TFs and in which tissue or developmental stage these effects are relevant.

### The impact of Neandertal-linked trans-eQTLs on modern human phenotype variation

Furthermore, we sought to investigate links between a-cTF aSNPs and their potential phenotypic effects. We therefore explored whether aSNPs of the 60 GTEx a-cTFs showed phenotype associations (P<5×10^−8^) in GWAS available in the EBI GWAS catalog (MacArthur et al., 2017), Immunobase and Biobank Japan (Materials and methods). We found that aSNPs of 13 a-cTF loci have significant associations, including a region on chromosome 12 (cis-eQTL TF: *STAT2*) which was also detected as a trans-eQTL in eQTLGen (Table S1,S2). The aSNPs at this locus (*rs2066807/rs2066819*) were associated with an decreased risk in autoimmune disease and less prevalent in individuals with psoriasis. This locus is linked to multiple additional phenotypes, including multiple anthropometric measures related to increased height and body mass and several pulmonary functional measurements. The aSNPs for seven additional a-cTFs (*ATOH7, CCDC88A, FOXC1, HOXA13, ZNF592* and *ZKSCAN4*) showed associations with height measures, making this by far the most frequently associated phenotype among a-cTFs (Table S6). We then compared the height associations (“Standing height” from UK Biobank P<5×10^−8^) each of the 133 a-cTF aSNPs across all tissues (with 90 unique aSNPs and 60 unique a-cTFs) to height associations in 100 sets of 133 random non-archaic SNPs with cTF cis-eQTLs and each set sampled to match the tissue prevalence of the 133 a-cTF aSNPs. We found that the height associations in the a-cTF aSNP set were on average 1.6 times higher compared to the background sets, a result that was borderline significant (P=0.06). Other non-height phenotype associations included breast cancer, grip strength, body fat measures, varicose veins, and abnormal red blood cell volumes.

In comparison, the 10,317 SNPs tested for trans-eQTL effects by Võsa & Claringbould et al. were originally selected so as to have significant associations in GWAS (P<5×10^−8^) in the EBI GWAS catalog (MacArthur et al., 2017), Immunobase and a blood trait GWAS. By this selection a link between these trans-eQTLs and phenotypic effects had already been established. We explored these phenotype databases and additional association data from Biobank Japan to annotate significant phenotype associations for the trans-eQTL aSNPs in eQTLGen (Table S7). We found that the majority of those candidates (8/15) were showing associations with various blood cell composition measures. These results are consistent with the fact that most of the tested and trans-eQTL-associated variants by Võsa & Claringbould et al. were linked to this group of phenotypes. However, one trans-eQTLs that also showed associations not related to blood cells was the one for aSNPs *rs13098911/rs13063635*. These aSNP showed associations with mouth ulcers and other dental issues. In addition, the archaic alleles for this risk locus have previously also been connected with increased risks for a severe Covid-19 phenotype (Zeberg and Pääbo, 2020) and Celiac disease, an autoimmune disorder, where dietary gluten intake causes inflammation in the small intestine at gluten intake (Caio et al., 2019). Two additional trans-eQTL loci (aSNPs *rs12908161* and *rs7811653*) showed associations with anthropometric and pulmonary measures. The directional effects for the phenotypes included increasing and decreasing height, weight and pulmonary measures. The remaining phenotype associations for aSNPs of other Neandertal trans-eQTLs were related to increased risk for colorectal cancer (*rs12603526*), electrocardiogram measurements (*rs10919070/rs10919071*), hypospadias (*rs7811653*), lipid measures *(rs6784615*), bone density (*rs72647484*), schizophrenia (*rs12908161*) and brain connectivity measurements (*rs16997087*).

In general our results suggest that aSNPs linked to both a-cTFs and trans-eQTL effects often show phenotype associations as well. The associated phenotypic effects range across multiple biological systems, including anthropomorphic measures, immunity, brain-related traits and cell type composition.

#### Evidence for local adaptation

One of the two a-cTFs with significant trans-eQTL effects in eQTLGen was associated with aSNPs on chromosome 12. The aSNPs fall in a genomic region that has previously been linked to a Neandertal DNA with signals of positive selection in Papuans (Mendez et al., 2012). We therefore re-evaluated the frequency distribution of all GTEx a-cTF and eQTLGen trans-eQTL aSNPs in present-day non-African populations to explore whether we can find evidence for selection for any of our candidate aSNPs. We quantified the frequencies of these aSNPs in 15 Eurasian populations from the 1,000 Genomes cohort and 16 SGDP Papuans and compared it to the frequency distribution of all detected aSNPs (1000 Genomes Project Consortium et al., 2015; Mallick et al., 2016). As previously reported by Mendez et al. we found that while the region on chromosome 12 showed frequencies of <10% in all 1,000 Genomes Eurasians, the frequency was, with 57%, substantially higher in Papuans (Figure 3A-B). This frequency in Papuans puts these aSNPs within the top 5% of all aSNPs in that population. This observation is even more remarkable for a Neandertal haplotype, given that in Papuans many aSNPs are derived from Denisovans and expected to have an on average higher frequency compared to their Neandertal counterparts. In total we found that aSNPs of nine empirical trans-eQTL candidates and 21 a-cTFs reached the top 5% allele frequencies distribution among detected aSNPs in a given population. Particular outliers were aSNPs associated with the a-cTFs *ATOH7, KAT2B, LMX1A, PLEK, PRDM5, SOX12, ZNF10* and *ZNF410* (Table S8) and three archaic trans-eQTLs: *rs13063635/rs13098911, rs13043612* and *rs11650665* (Table S9), which were found at archaic allele frequencies in the top 1% among aSNPs. Most of the aSNPs that reached frequencies within the top 1% aSNP distribution in a given population, showed the most extreme frequencies in Europeans and are found at elevated frequencies in multiple European and Asian populations (Figure 3B, Tables S8,S9). However, we also observed instances where such aSNPs are entirely absent or only found at extremely low frequencies in one geographic area (a-cTFs *ZNF410, KAT2B*) or show the highest frequencies in non-European populations (a-cTFs *LMX1A, SOX12* and trans-eQTL *rs13063635/rs13098911*). These results are consistent with a detection bias of association-study-based functional annotation of aSNPs that are more prevalent in Europeans, due to the general over-representation of expression and phenotype data from cohorts of predominantly European ancestry (Dannemann et al., 2020), including GTEx and eQTLGen. The results also illustrate that some of the high-frequency aSNPs show population-specific frequencies, consistent with potential local selection pressures.

**Figure 3:**
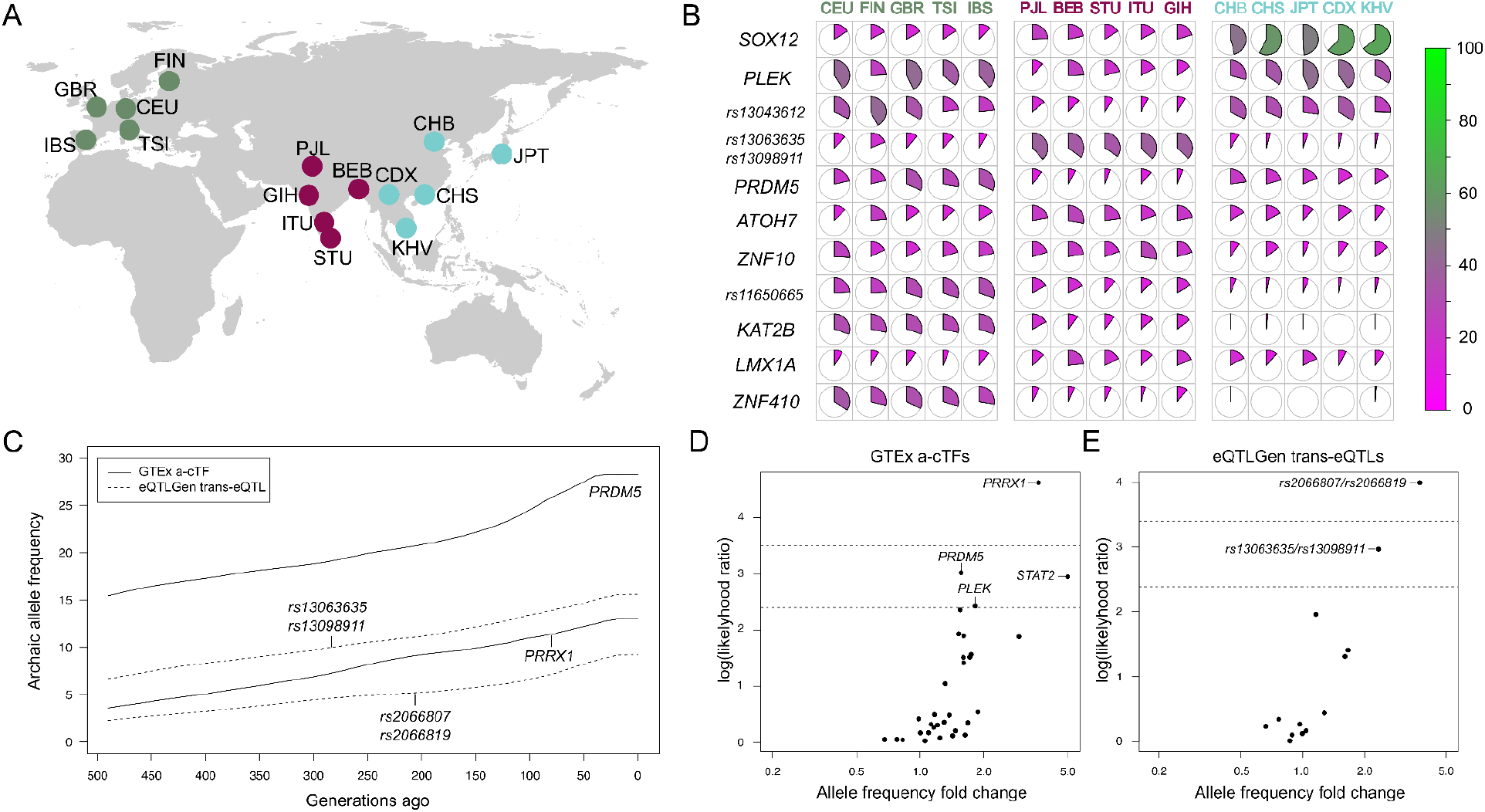
The frequency of trans-eQTL aSNPs in Eurasian populations. **(A)** Geographic distribution of 15 Eurasian 1,000 Genomes populations, colors match the populations’ names in panel B. **(B)** Pie charts illustrating the frequencies of a-cTF aSNPs (annotated with the corresponding a-cTF) and trans-eQTL (annotated with the rs ID) aSNPs (y-axis) that fall into the top 1% aSNP distribution in at least one of 15 Eurasian 1,000 Genomes populations (x axis). **(C)** The estimated posterior probability for the Neandertal allele frequency for two a-cTF and two trans-eQTL aSNP candidates with logLRs above the 90^th^ percentile of the background distributions over the last 500 generations in the Estonian population are shown. For each significant a-cTF **(D)** and trans-eQTL **(E)** aSNP locus the average frequency change in the Estonian population over the last 500 generations (x-axis) and the corresponding average log likelihood ratio (y axis) are displayed. The 90^th^ and 95^th^ percentiles for a given logLR distribution are shown as dotted lines.

Additionally, we sought to explore whether any of the a-cTF and trans-eQTLs aSNPs show evidence for recent directional allele frequency changes, possibly as a result of selective pressures. For that we used a dataset of around 1,800 whole genome sequences from the Estonian Biobank as by leveraging such a big sample from a rather homogeneous population we can gain power in robustly reconstructing frequency changes in the recent past. We first generated local genealogical trees for this dataset using Relate (Speidel et al., 2019) and then applied CLUES (Stern et al., 2019), an approximate full-likelihood method for inferring allele frequency trajectories to our trans-eQTL candidate aSNPs (Materials and methods). This analysis revealed aSNPs that experienced either an increase or a decrease in the frequency of the Neandertal allele over the last 500 generations. Most of those changes were characterized by low log likelihood ratios (logLR) suggesting little evidence for rejecting neutrality (Figure 3D-F, Table S10,S11). In order to assess whether any of the reconstructed frequency changes are unexpectedly high, we compared the logLRs to two background distributions, one per a database. The logLRs of trans-eQTL-associated aSNPs were compared to logLRs of the set of ∼2,900 significant trans-eQTL SNPs from eQTLGen that we were able to test in the Estonian dataset. For a-cTF-associated aSNPs we generated a total of 370 non-archaic SNPs of cTFs in GTEx that show the same allele frequency distribution and tissue prevalence as aSNPs of the 60 Neandertal a-cTFs (Materials and methods).

We tested 40 a-cTFs aSNPs (associated with 36 a-cTFs) passing our minimum minor allele frequency (5%) and quality filters (Materials and methods). We found that aSNPs associated with four a-cTFs (*PLEK, PRDM5, PRRX1* and *STAT2*) showed logLRs that fell within the top 10% distribution of logLRs for background SNPs. Notably, all four a-cTFs aSNP were cis-eQTLs in brain tissues, including *PRRX1* (Caudate basal ganglia), which showed the largest logLRs and the only instance that fell within the top 5% logLR background distribution (97^th^ percentile). Unlike the other three a-cTFs, the corresponding aSNP for *PRRX1* (*rs77401317*) does not show allele frequencies that are within the upper extreme (>95^th^ percentile) of the aSNP distribution in present-day populations (Table S10). In fact this SNP is absent in East Asians, and found at frequencies between 2-4% in South Asians and 9-13% in Europeans.

Furthermore, we also found that the archaic allele frequencies for two trans-eQTL candidates increased by more than two-fold over the last 500 generations in the population ancestral to Estonians (*rs13063635*/*rs13098911* and *rs2066807*/*rs2066819*). Compatible with this observation, the corresponding logLRs for each of the two trans-eQTL candidates aSNP pairs were high compared to ∼2,900 significant trans-eQTL SNPs from eQTLGen, reaching the 93^rd^ (*rs13063635*/*rs13098911* and 97^th^ percentile for only *rs13063635*) and 97^th^ (*rs2066807*/*rs2066819*) percentile of the trans-eQTL SNP logLR distribution (Table S11, Materials and methods).

Both analysis on the frequencies of a-cTF and trans-eQTL-associated aSNPs in populations and recent frequency changes in Estonian genomes suggest that some of these archaic variants show exceptionally high frequencies in some present-day populations and frequency changes that are suggestive of non-neutral processes like positive selection. However, the example of *PRRX1* together with the aSNPs at the *STAT2* locus also demonstrate that some archaic variants that are not necessarily frequency outliers (frequencies below 10%) in a given population nevertheless show signals or recent selective pressures.

## Discussion

In this study we explored the GTEx tissue expression dataset for Neandertal DNA associations with expression changes in TFs. We have then used this information together with TF target gene data to predict potential genome-wide regulatory effects that are initiated via the expression modulation of TFs. We detected 60 TFs that had in at least one of the 49 GTEx tissues, their top cis-eQTL SNP being of likely Neandertal ancestry (a-cTFs). Using protein-protein interaction information we show that the connectivity between these a-cTFs is larger than interaction between other TFs with GTEx cis-eQTL with non-archaic SNPs. The a-cTF with the largest number of interactions with other a-cTFs was *JUN*. Notably, *JUN* was one of two a-cTFs (*PRDM5*) with predicted target genes being enriched in genomic regions devoid of Neandertal and Denisovan DNA. These regions have previously been suggested to be a result of negative selection against introgressed DNA and have been shown to be enriched for brain-related genes and depleted of testes expressed genes (Buisan et al., 2021; Sankararaman et al., 2016; Vernot et al., 2016). Our results suggest that these regions are not out of regulatory reach for Neandertal DNA and that in fact some TFs that show cis-eQTLs with Neandertal DNA particulary target genes in these regions. Interestingly, both *JUN* and *PRDM5* were aSNP-associated cis-eQTLs in brain tissues (*JUN*: Cortex; *PRDM5*: Frontal cortex, spleen), an observation providing a potential link to the phenotypic consequences of the predicted regulatory effects. This observation is also consistent with our functional enrichment results of the 60 a-cTFs, that included with circadian rhythm and hippocampus development two GO categories that were over-represented in a-cTFs - a result partially driven by *JUN* as well (Table S3).

Both *JUN* and *PRDM5* were also among the 14 a-cTFs with archaic allele frequencies that reached the top 2% of aSNPs segregating in at least one 1,000 Genomes population (highest values: *JUN* 98^th^ percentile in Toscani / *PRDM5* 99^th^ percentile in Iberians). In addition, a-cTF aSNPs of *PRDM5* also showed substantial frequency increases of more than 10 percentage points in the last 500 generations in our Estonian test dataset, suggesting that positive selection was possibly acting on these archaic variants in the last ∼10,000 years. Strikingly, each of the four a-cTFs associated with aSNPs that showed substantial recent frequency changes (top 10% logLR background distribution for *PLEK, PRDM5, PRRX1* and *STAT2*) showed among their cis-eQTL associations one related to a brain tissue (Table S2). This result is in contrast with the affected tissues of a-cTFs that show associated aSNPs with frequencies in the top 1% aSNP distribution in 1,000 Genomes populations. Here, only two of the eight such a-cTF show a brain tissue cis-eQTL (*PLEK, PRDM5*). These results are compatible with recent selection particularly targeting brain-related a-cTFs.

Another a-cTF with aSNPs showing evidence for recent selection was *STAT2*. The cis-eQTL aSNPs for *STAT2* were consistently found at frequencies below 10% in 1,000 Genomes populations, frequencies that were lower than for many other a-cTF aSNPs in our study (maximum allele frequency for *STAT2* aSNPs: 9%; 24% of a-cTFs have lower maximum aSNP frequency, Table S8). Only in Papuans we found that *STAT2* aSNPs showed frequencies of 57%, a value that is equivalent with the 97^th^ percentile of frequencies of aSNPs found in that population. These results are also consistent with previous reports on positive selection on Neandertal DNA in the regions of *STAT2* in Papuans (Mendez et al., 2012). However, our analysis of predicted frequency changes over the last 500 generations suggests that the *STAT2* aSNPs might have been a target of positive selection outside of Oceania as well. *STAT2* showed significant aSNP cis-eQTLs in seven GTEx tissues and also eQTLGen. Interestingly, the directional effects differed between tissues. In two GTEx tissues (Cerebellar hemisphere and liver) the expression of *STAT2* was higher in archaic allele carriers. Conversely, the expression of *STAT2* in the blood expression data of eQTLGen and in GTEx suprapubic skin, two adipose tissue, tibial nerve and artery was lower in the presence of the archaic allele. These results suggest tissue-specific function of the archaic alleles in different tissues.

However, our results are primarily based on information from association analyses and computational prediction algorithms. It is therefore conceivable that some of our candidate aSNP are in high linkage disequilibrium with non-archaic variants that are truly causal. This point is even more relevant considering that the variant with the strongest effect size doesn’t necessarily translate to actual molecular consequences (Battle and Montgomery, 2014). In addition we note that the TF target gene prediction algorithms we used in our study are based on different prediction strategies, including computational frameworks and inferences based on experimental data. Consequently the accuracy of these algorithms varies (Jayaram et al., 2016) and is hard to exactly quantify as it often remains unclear in which tissue or developmental stage the predicted effects are functionally relevant. Nevertheless, particularly the example of *STAT2* has shown that the continuously growing functional information can help in providing robust evidence for functional effects of the archaic variants. First, the fact that aSNPs were associated with significant cis-eQTLs in multiple tissues in two databases - GTEx and eQTLGen - provides a robust support for a functional link between Neandertal DNA and the regulation of this TF across tissues. Second, the eQTLGen trans-eQTL screen includes a link between two aSNPs in the *STAT2* region and IFI16, a predicted target of *STAT2*, adding an empirical example that is consistent with the TF target gene prediction for this TF. Third, a recent study that experimentally tested aSNPs for their regulatory potential to modulate gene expression identified aSNPs with such a potential and using CRISPR technologies demonstrated that three of the detected variants modified the expression of *STAT2* in an immune cell line (Table S12, (Jagoda et al., 2021)).

Specifically technologies that enable the experimental testing of variants in a high-throughput manner are highly valuable for Neandertal DNA, that often segregates on long haplotypes carrying tens or even more than 100 aSNPs in high LD. The high LD between these SNPs makes it practically impossible to disentangle functional information from association-based analyses. A similar study by the same authors has run a second experiment to investigate the regulatory potential of a Covid-associated Neandertal haplotype (Jagoda et al., n.d.). The tested aSNPs in this study overlap with a second significant trans-eQTL locus that we identified in eQTLGen and is linked to the aSNPs *rs13063635* and *rs13098911* (Table S12). These two aSNPs were in LD with three expression modulating aSNPs that were identified by this study, providing candidates for the underlying regulatory effects. This second locus was together with the *STAT2* region not only the two examples of experimental testing of putative causal variants but also the only two trans-eQTL aSNP loci in eQTLGen that showed evidence of positive selection. In addition, both loci also showed common phenotype associations including autoimmune disorders (psoriasis and Celiac disease, Tables S2,S3), suggesting that Neandertal DNA influences such phenotypes via the modulation of gene expression. In general, a total of 12 of the 60 GTEx a-cTF loci showed significant phenotype associations, too (Table S6). Notably, seven of these loci had associations with height and other body measures. Another four loci showed links to blood cell type composition. No other phenotype had associations with more than one of the 12 a-cTF loci. An additional eight of the 15 eQTLGen trans-eQTL loci showed associations with blood cell type measures. The only other group of phenotypes beside autoimmune diseases and blood cell types with more than one locus linked to it was height, with three loci linked to it (Table S7). And while these observations clearly indicate that many of the candidate aSNPs for putative long-range regulatory effects could also result in phenotypic effects on various disease and non-disease phenotypes, it remains hard to quantify how these results translate to the entire set of Neandertal DNA in people today. First, in this study we have investigated only one potential regulatory mechanism, i.e. the expression modulation of a TF and implications for its predicted targets. In order to complete this picture, future genome-wide trans-eQTL screens can further test the robustness of our inferences. Second, given that most of our inferences are based on association analyses, we are more likely to pick up variants with an on average higher minor allele frequency, likely biasing our current view on aSNPs with elevated frequencies.

Nevertheless it is intriguing to observe that highly polygenic phenotypes such as height (Yang et al., 2010) were often linked to regulatory activity of a-cTFs who themselves have the ability to influence expression levels of large sets of genes and therefore represent a polygenic equivalent on the transcriptomic level. This observation, together with the elevated connectivity of aSNP a-cTFs is compatible with a complex regulatory machinery that Neandertal DNA in modern humans is still significantly influencing. It has previously been proposed that complex regulatory networks are underlying the genetic basis of complex traits (Boyle et al., 2017). Our results on the expression modulation of multiple highly interactive TFs by introgressed archaic DNA, that each of them control hundreds or even thousands of genes, would be very much compatible with this model.

Our study also provides a new perspective on the role of Neandertal DNA on archaic desert in the genomes of present-day people. Previous work has often considered these deserts as genomic regions that are unlinked from introgressed archaic DNA. Our results suggest that these regions are not out of reach for the regulatory and potentially phenotypic impact of Neandertal DNA. Particularly TFs that are active in the brain and involved in brain-related functions, like circadian rhythm, showed various predicted target genes in these regions. These observations might be helpful in the ongoing interpretation of both previous links between introgressed Neandertal DNA and behavioral phenotypes (Dannemann and Kelso, 2017; Simonti et al., 2016) and the extent of potential phenotypic differences between modern and archaic humans.

In our study we explored the potential genome-wide impact of Neandertal DNA that is associated with expression changes in TFs. Võsa & Claringbould et al. have demonstrated that at least 9% of trans-eQTLs can be explained by this mechanism (Võsa et al., 2021). Prior estimates of ∼20% of the general expression variation that can be explained by trans-eQTLs (Ouwens et al., 2020) suggest that the detected TFs in our study could be responsible for ∼1.8% of the expression variation initiated by Neandertal DNA. This proportion is substantial, considering that all cis-eQTLs are estiamted to be responsible for only ∼6% of expression variation. Consistent with other previous studies that have investigated the regulatory importance of Neandertal DNA and its potential phenotypic consequences for present-day people (Barker et al., 2020; Colbran et al., 2019; Dannemann et al., 2017; Gittelman et al., 2016; Gokhman et al., 2020; McCoy et al., 2017; Petr et al., 2019; Silvert et al., 2019; Telis et al., 2020) we show that the inferred genome-wide long-range regulatory implication of archaic DNA that we investigated in our study is associated with various phenotypic effects and some of the underlying archaic variants potential targets of positive selection.

## Materials and methods

### eQTL datasets

We downloaded cis and trans-eQTL summary statistics from eQTLGen (Võsa et al., 2021) and GTEx (Consortium and The GTEx Consortium, 2020) [v8]. We annotated significant cis and trans-eQTLs based on an FDR cutoff of 0.05. TFs with significant cis-eQTLs in GTEx were defined as cTF and cTFs that were associated with an archaic SNP (see definition in next section) were called a-cTFs.

### Annotation of introgressed Neandertal variants

We identified Neandertal variants in the eQTLGen, GTEx and the phenotype cohorts described below based on a list of previously annotated introgressed Neandertal marker variants, referred to as aSNPs (Dannemann, 2021). These aSNPs have been annotated in the 1,000 genomes project dataset (v5a) (1000 Genomes Project Consortium et al., 2015; Dannemann, 2021) and were defined based on the following conditions: (i) one allele is fixed in the African Yoruba population (ii) the second allele is present in a homozygous state at least in one of the genomes of the Altai Neandertal, Vindjia Neandertal or Denisovan (Meyer et al., 2012; Prüfer et al., 2017, 2014) and (iii) the allele in (ii) being present in at least one non-African 1,000 Genomes individual. We (iv) required these inferred aSNPs to overlap with previously annotated haplotypes that have been shown to exceed a length that is consistent with incomplete lineage sorting (ILS), a genomic phenomenon that can also result in allele sharing properties described in (i-iii). We detected 384,896 autosomal aSNPs in the 1,000 Genomes that met these criteria. In addition we detected 151,388 aSNPs in 16 Papuans from the Simons Genome Diversity Project (SGDP) cohort (Mallick et al., 2016) applying the same criterias (i-ii) and the modified condition (iii) that the archaic allele being required to be detected across Papuans in the SGDP cohort. Finally, we required each candidate aSNP in GTEx and eQTLGen to show r^2^>0.5 with at least three additional aSNPs that passed (i-iv).

### Enrichment analysis of Neandertal trans-eQTLs in eQTLGen

We detected 77 aSNPs among the 10,317 SNPs that have been included in the eQTLGen trans-eQTL screen. Among those aSNPs were 91 pairs (31 aSNPs) that showed elevated levels of linkage disequilibrium (LD) (r^2^>0.5). We collapsed these aSNPs into nine loci and choose one random aSNP to represent a given locus, which resulted in 55 independent aSNPs. Out of these variants a total of 15 aSNPs showed at least one significant (FDR<0.05) trans-eQTL association. We then compared the proportion of significant Neandertal trans-eQTLs to the fraction in 100 same size random sets of frequency-matched non-archaic variants. To this end, we LD pruned all tested SNPs (10,317) based on r^2^ > 0.5 in European populations from 1,000 Genomes (1000 Genomes Project Consortium et al., 2015). We counted significant trans-eQTLs in non-archaic sets, compared these numbers to the value of significant aSNPs, and calculated an empirical two-sided p-value.

### Connectivity and functional enrichment of Neandertal a-cTFs

We constructed protein-protein interaction networks using information about human protein interactions from the STRING database (v11, (Szklarczyk et al., 2019)). The network displayed in Figure 2A is based on the default minimum median confidence of 0.4 and a k-means clustering (k=5). The average connectivity between a-cTFs and background sets of non-archaic cTFs were calculated based on all interactions without any confidence cutoff applied.

In addition we tested for functional enrichment of a-cTFs in the Gene Ontology (GO, (Harris et al., 2004)), KEGG (Kanehisa and Goto, 2000) and REACTOME (Gillespie et al., 2022) using the enrichment software implemented in STRING. Gene pathway and GO categories with an odds ratio > 1 and false discovery rate < 0.05 were considered to be enriched for a-cTFs.

### TF target gene prediction databases

We leveraged TF target gene prediction information from seven databases. The target gene information in these databases is based on computational prediction algorithms (MotifMap (Xie et al., 2009), Human TFTG (“Causal Mechanistic Regulatory Network for Glioblastoma Deciphered Using Systems Genetics Network Analysis,” 2016), JASPAR (Mathelier et al., 2014), TRANSFAC predicted (Matys et al., 2006)) and experimental information (ENCODE (“A user’s guide to the encyclopedia of DNA elements (ENCODE),” 2011), CHEA (Lachmann et al., 2010), TRANSFAC curated (Matys et al., 2006)). The numbers of evaluated TFs and predicted targets varies substantially between these databases (Table S4,S5). In order to ensure comparability between information from these databases we equalized gene names from all databases to common GeneSymbols, collapsed duplexes and limited the analysis to protein coding genes. Human TFs without target gene information in any of these databases were annotated based on AnimalTFDB3.0 (Hu et al., 2018).

### Phenotype databases

We extracted phenotypic information for both a-cTFs and trans-eQTL associated aSNPs from EBI GWAS catalog (https://www.ebi.ac.uk/gwas/) (MacArthur et al., 2017), Immunobase, a blood cell traits study (Astle et al., 2016) and the Biobank Japan for associations with candidate aSNPs (Table S6, S7). We considered all phenotype associations with P<5×10^−8^.

### Definition of archaic deserts

We used information on shared archaic deserts of Neandertal and Denisovan introgression from the studies of Sankararaman et al. and Vernot et al. (Sankararaman et al., 2016; Vernot et al., 2016). We considered all autosomal deserts reported by both studies and collapsed overlapping deserts into one by using the range of the combined regions. That resulted in five deserts that were defined by the following hg19 coordinates: chr1:99Mb-114.9Mb, chr3:76.5Mb-90.5Mb, chr7:108Mb-128Mb, chr8:54.5Mb-65.4Mb and chr13:49Mb-61Mb. We defined desert genes as those entirely located within a desert’s borders.

### Evidence for natural selection acting on trans-eQTL aSNPs

We used CLUES (Stern et al., 2019) to estimate changes in the allele frequency of the aSNPs over the last 500 generations and to test whether these changes might have happened under natural selection. CLUES was run as in Marnetto et al. (Marnetto et al., n.d.). Briefly, we started with building local trees by applying Relate (version 1.1.4) (Speidel et al., 2019) to whole genome sequences of 2,420 Estonian Biobank participants (dataset is described in (Kals et al., 2019; Pankratov et al., 2020)). When running Relate for tree building we used the strict callability mask, recombination map and the reconstructed human ancestral genome all generated based on the 1,000 Genomes Project (GRCh37), a mutation rate of 1.25×10^−8^ and an effective population size of 30,000. Next, we extracted subtrees corresponding to a subset of 1,800 individuals by removing individuals that were related (relatives up to 3rd degree), PCA or singleton count outliers and individuals with excessive IBD sharing (≥ 166.2 cM) with 2 or more other individuals in the dataset. This step was implemented to run CLUES on a more homogeneous dataset and to reduce run time. Coalescent rate over time for the Estonian population needed for CLUES was estimated based on 100 randomly sampled individuals. When running CLUES we extracted the local tree with the SNP of interest and re-sampled its branch length 200 times. We then removed te first 100 samples and took every 5^th^ tree (20 in total) for importance sampling.

We applied this analysis to two sets of aSNPs and two corresponding background sets. The first set of aSNPs (predicted archaic trans-eQTLs, a-cTF) altogether included 40 of the 90 a-cTF aSNPs after quality filters. Our filtering criteria removed 23 aSNPs with a minor allele frequency (MAF) below 5% in 1,800 individuals of the Estonian whole genome sequence data, 17 aSNPs not passing the applied genomic mask and two not mapping to Relate trees. From the remaining 48 aSNPs we kept one aSNP per each group of aSNPs in LD of r^2^ >= 0.6, resulting in a set of 35 unlinked aSNPs. For a-cTF aSNPs not passing above-mentioned filters we searched for aSNPs that are in LD (r^2^ >= 0.6) outside of the original set of 90 GTEx aSNPs. We found 5 additional aSNPs that we used as proxies for those a-cTF aSNPs in the CLUES analysis.

In addition, we generated a background set for the a-cTF aSNPs based on 2,404 cTF SNPs with no evidence of Neandertal ancestry. We removed 188 SNPs without genotype information and 377 SNPs with MAF below 5% in 1,800 Estonians as well as 560 SNPs not passing the genomic mask, resulting in a total of 1,279 SNPs. Next, for each of the 40 a-cTF described above we sampled 10 SNPs out of this pool of cTF SNPs without replacement and matching for MAF and tissue prevalence of a-cTF aSNPs. This resulted in a set of 400 unique cTF SNPs which we LD-pruned at r^2^ >= 0.6 ending up with a background set of 370 cTF SNPs.

To obtain the second set of aSNPs we started with the 18 trans-eQTL aSNPs and removed two that had a MAF<5% in the Estonian cohort. One aSNP that was not mapped to a local tree was replaced by a proxy aSNP with r^2^ of 0.98. Thus, 16 trans-eQTL aSNPs were analyzed. We also generated a background dataset for trans-eQTL aSNP by running CLUES as described above for 2,826 (out of 3,835 significant) trans-eQTL non-archaic SNPs being present on our local trees and having MAF >= 5% in 1,800 Estonians.

To account for the variation in the results coming from the uncertainty in branch length estimation we ran the analysis (starting from sampling branch length) twice for each a-cTF and trans-eQTL aSNP and took the mean of the frequencies and log likelihood ratios. For the three pairs of trans-eQTL aSNPs in LD (r^2^>0.5) we ran CLUES for each SNPs and then averaged the estimates within each pair. In the case of both background sets CLUES was run once for each SNP.

## Acknowledgments

We would like to thank Mayukh Mondal for his comments on the manuscript. Some of the analyses were carried out with the facilities of the High-Performance Computing Center of the University of Tartu.

## Consortia

Estonian Biobank Research Team:

Andres Metspalu, Mari Nelis, Lili Milani, Reedik Mägi & Tõnu Esko

## Data, script and code availability

### Script and code availability

Code related to the analyses presented in this manuscript is available at:

- https://github.com/SillySabertooth/Neandertal_trans-eQTLs
- https://myersgroup.github.io/relate/
- https://github.com/35ajstern/clues

### Data availability

*Summary statistics for eQTLs:*

- eQTLGen: https://www.eqtlgen.org/
- GTEx: https://gtexportal.org/home/datasets

*Transcription factor target prediction information:*

- https://maayanlab.cloud/Harmonizome/dataset/
- http://tfbsdb.systemsbiology.net/download

*GWAS summary statistics:*

- Immunobase: https://genetics.opentargets.org/immunobase
- GWAS catalog: https://www.ebi.ac.uk/gwas/

*Genotype data*

- Neandertal and Denisovan genomes: http://ftp.eva.mpg.de/neandertal/ and http://ftp.eva.mpg.de/denisova
- 1,000 Genomes: http://ftp.1000genomes.ebi.ac.uk/vol1/ftp/
- Simons Genome Diversity Project: https://www.simonsfoundation.org/simons-genome-diversity-project/, respectively
- Estonia Biobank WGS: https://genomics.ut.ee/en/access-biobank (Estonian Biobank ethics approval number nr 1.1-12/2859)

## Supplementary information

**Table S1: Summary statistics for significant trans-eQTLs associated with aSNPs in eQTLGen**.

**Table S2: Summary statistics for aSNPs of a-cTFs in GTEx**.

**Table S3: Functional enrichments results**

**Table S4: Transcription factor target information**.

**Table S5: Overlap in available transcription factors with prediction information between databases**.

**Table S6: Significant phenotype associations for GTEx a-cTF aSNPs**.

**Table S7: Significant phenotype associations for trans-eQTL associated aSNPs**.

**Table S8: Population allele frequency information for a-cTF aSNPs**

**Table S9: Population allele frequency information for trans-eQTL aSNPs**

**Table S10: CLUES allele fFrequency trajectory estimates for a-cTF aSNPs**.

**Table S11: CLUES allele frequency trajectory estimates for trans-eQTL aSNPs**.

**Table S12: MPRA regulatory information for trans-eQTL and a-cTF aSNPs**.

## References

1000 Genomes Project Consortium, Auton A, Brooks LD, Durbin RM, Garrison EP, Kang HM, Korbel JO, Marchini JL, McCarthy S, McVean GA, Abecasis GR. 2015. A global reference for human genetic variation. Nature 526:68–74.

Astle WJ, Elding H, Jiang T, Allen D, Ruklisa D, Mann AL, Mead D, Bouman H, Riveros-Mckay F, Kostadima MA, Lambourne JJ, Sivapalaratnam S, Downes K, Kundu K, Bomba L, Berentsen K, Bradley JR, Daugherty LC, Delaneau O, Freson K, Garner SF, Grassi L, Guerrero J, Haimel M, Janssen-Megens EM, Kaan A, Kamat M, Kim B, Mandoli A, Marchini J, Martens JHA, Meacham S, Megy K, O’Connell J, Petersen R, Sharifi N, Sheard SM, Staley JR, Tuna S, van der Ent M, Walter K, Wang S-Y, Wheeler E, Wilder SP, Iotchkova V, Moore C, Sambrook J, Stunnenberg HG, Di Angelantonio E, Kaptoge S, Kuijpers TW, Carrillo-de-Santa-Pau E, Juan D, Rico D, Valencia A, Chen L, Ge B, Vasquez L, Kwan T, Garrido-Martín D, Watt S, Yang Y, Guigo R, Beck S, Paul DS, Pastinen T, Bujold D, Bourque G, Frontini M, Danesh J, Roberts DJ, Ouwehand WH, Butterworth AS, Soranzo N. 2016. The Allelic Landscape of Human Blood Cell Trait Variation and Links to Common Complex Disease. Cell 167:1415–1429.e19.

A user’s guide to the encyclopedia of DNA elements (ENCODE). 2011.. PLoS Biol 9. doi:10.1371/journal.pbio.1001046

Barker HR, Parkkila S, Tolvanen MEE. 2020. Evolution is in the details: Regulatory differences in modern human and Neanderthal. bioRxiv. doi:10.1101/2020.09.04.282749

Battle A, Montgomery SB. 2014. Determining causality and consequence of expression quantitative trait loci. Hum Genet 133:727–735.

Boyle EA, Li YI, Pritchard JK. 2017. An Expanded View of Complex Traits: From Polygenic to Omnigenic. Cell 169:1177–1186.

Buisan R, Moriano J, Andirkó A, Boeckx C. 2021. A distinct brain expression profile for genes within large introgression deserts and under positive selection in Homo sapiens. bioRxiv. doi:10.1101/2021.03.26.437167

Caio G, Volta U, Sapone A, Leffler DA, De Giorgio R, Catassi C, Fasano A. 2019. Celiac disease: a comprehensive current review. BMC Med 17:142.

Causal Mechanistic Regulatory Network for Glioblastoma Deciphered Using Systems Genetics Network Analysis. 2016.. Cell Systems 3:172–186.

Colbran LL, Gamazon ER, Zhou D, Evans P, Cox NJ, Capra JA. 2019. Inferred divergent gene regulation in archaic hominins reveals potential phenotypic differences. Nat Ecol Evol 3:1598–1606.

Consortium TG, The GTEx Consortium. 2020. The GTEx Consortium atlas of genetic regulatory effects across human tissues. Science. doi:10.1126/science.aaz1776

Dannemann M. 2021. The Population-Specific Impact of Neandertal Introgression on Human Disease. Genome Biol Evol 13. doi:10.1093/gbe/evaa250

Dannemann M, He Z, Heide C, Vernot B, Sidow L, Kanton S, Weigert A, Treutlein B, Pääbo S, Kelso J, Camp JG. 2020. Human Stem Cell Resources Are an Inroad to Neandertal DNA Functions. Stem Cell Reports 15:214–225.

Dannemann M, Kelso J. 2017. The Contribution of Neanderthals to Phenotypic Variation in Modern Humans. Am J Hum Genet 101:578–589.

Dannemann M, Prüfer K, Kelso J. 2017. Functional implications of Neandertal introgression in modern humans. Genome Biol 18:61.

Gillespie M, Jassal B, Stephan R, Milacic M, Rothfels K, Senff-Ribeiro A, Griss J, Sevilla C, Matthews L, Gong C, Deng C, Varusai T, Ragueneau E, Haider Y, May B, Shamovsky V, Weiser J, Brunson T, Sanati N, Beckman L, Shao X, Fabregat A, Sidiropoulos K, Murillo J, Viteri G, Cook J, Shorser S, Bader G, Demir E, Sander C, Haw R, Wu G, Stein L, Hermjakob H, D’Eustachio P. 2022. The reactome pathway knowledgebase 2022. Nucleic Acids Res 50:D687–D692.

Gittelman RM, Schraiber JG, Vernot B, Mikacenic C, Wurfel MM, Akey JM. 2016. Archaic Hominin Admixture Facilitated Adaptation to Out-of-Africa Environments. Curr Biol 26:3375–3382.

Gokhman D, Nissim-Rafinia M, Agranat-Tamir L, Housman G, García-Pérez R, Lizano E, Cheronet O, Mallick S, Nieves-Colón MA, Li H, Alpaslan-Roodenberg S, Novak M, Gu H, Osinski JM, Ferrando-Bernal M, Gelabert P, Lipende I, Mjungu D, Kondova I, Bontrop R, Kullmer O, Weber G, Shahar T, Dvir-Ginzberg M, Faerman M, Quillen EE, Meissner A, Lahav Y, Kandel L, Liebergall M, Prada ME, Vidal JM, Gronostajski RM, Stone AC, Yakir B, Lalueza-Fox C, Pinhasi R, Reich D, Marques-Bonet T, Meshorer E, Carmel L. 2020. Differential DNA methylation of vocal and facial anatomy genes in modern humans. Nat Commun 11:1189.

Hardy K, Brand-Miller J, Brown KD, Thomas MG, Copeland L. 2015. THE IMPORTANCE OF DIETARY CARBOHYDRATE IN HUMAN EVOLUTION. Q Rev Biol 90:251–268.

Harris MA, Clark J, Ireland A, Lomax J, Ashburner M, Foulger R, Eilbeck K, Lewis S, Marshall B, Mungall C, Richter J, Rubin GM, Blake JA, Bult C, Dolan M, Drabkin H, Eppig JT, Hill DP, Ni L, Ringwald M, Balakrishnan R, Cherry JM, Christie KR, Costanzo MC, Dwight SS, Engel S, Fisk DG, Hirschman JE, Hong EL, Nash RS, Sethuraman A, Theesfeld CL, Botstein D, Dolinski K, Feierbach B, Berardini T, Mundodi S, Rhee SY, Apweiler R, Barrell D, Camon E, Dimmer E, Lee V, Chisholm R, Gaudet P, Kibbe W, Kishore R, Schwarz EM, Sternberg P, Gwinn M, Hannick L, Wortman J, Berriman M, Wood V, de la Cruz N, Tonellato P, Jaiswal P, Seigfried T, White R, Gene Ontology Consortium. 2004. The Gene Ontology (GO) database and informatics resource. Nucleic Acids Res 32:D258–61.

Hu H, Miao Y-R, Jia L-H, Yu Q-Y, Zhang Q, Guo A-Y. 2018. AnimalTFDB 3.0: a comprehensive resource for annotation and prediction of animal transcription factors. Nucleic Acids Res 47:D33–D38.

Inchley CE, Larbey CDA, Shwan NAA, Pagani L, Saag L, Antão T, Jacobs G, Hudjashov G, Metspalu E, Mitt M, Eichstaedt CA, Malyarchuk B, Derenko M, Wee J, Abdullah S, Ricaut F-X, Mormina M, Mägi R, Villems R, Metspalu M, Jones MK, Armour JAL, Kivisild T. 2016. Selective sweep on human amylase genes postdates the split with Neanderthals. Sci Rep 6:37198.

Jagoda E, Marnetto D, Montinaro F, Richard D, Pagani L, Capellini TD. n.d. Regulatory dissection of the severe COVID-19 risk locus introgressed by Neanderthals. doi:10.1101/2021.06.12.448149

Jagoda E, Xue JR, Reilly SK, Dannemann M, Racimo F, Huerta-Sanchez E, Sankararaman S, Kelso J, Pagani L, Sabeti PC, Capellini TD. 2021. Detection of Neanderthal Adaptively Introgressed Genetic Variants that Modulate Reporter Gene Expression in Human Immune Cells. Mol Biol Evol. doi:10.1093/molbev/msab304

Jayaram N, Usvyat D, R Martin AC. 2016. Evaluating tools for transcription factor binding site prediction. BMC Bioinformatics 17:547.

Kals M, Nikopensius T, Läll K, Pärn K, Sikka TT, Suvisaari J, Salomaa V, Ripatti S, Palotie A, Metspalu A, Esko T, Palta P, Mägi R. 2019. Advantages of genotype imputation with ethnically matched reference panel for rare variant association analyses. bioRxiv. doi:10.1101/579201

Kanehisa M, Goto S. 2000. KEGG: Kyoto Encyclopedia of Genes and Genomes. Nucleic Acids Res 28:27–30.

Lachmann A, Xu H, Krishnan J, Berger SI, Mazloom AR, Ma’ayan A. 2010. ChEA: transcription factor regulation inferred from integrating genome-wide ChIP-X experiments. Bioinformatics 26. doi:10.1093/bioinformatics/btq466

MacArthur J, Bowler E, Cerezo M, Gil L, Hall P, Hastings E, Junkins H, McMahon A, Milano A, Morales J, Pendlington ZM, Welter D, Burdett T, Hindorff L, Flicek P, Cunningham F, Parkinson H. 2017. The new NHGRI-EBI Catalog of published genome-wide association studies (GWAS Catalog). Nucleic Acids Res 45:D896–D901.

Mallick S, Li H, Lipson M, Mathieson I, Gymrek M, Racimo F, Zhao M, Chennagiri N, Nordenfelt S, Tandon A, Skoglund P, Lazaridis I, Sankararaman S, Fu Q, Rohland N, Renaud G, Erlich Y, Willems T, Gallo C, Spence JP, Song YS, Poletti G, Balloux F, van Driem G, de Knijff P, Romero IG, Jha AR, Behar DM, Bravi CM, Capelli C, Hervig T, Moreno-Estrada A, Posukh OL, Balanovska E, Balanovsky O, Karachanak-Yankova S, Sahakyan H, Toncheva D, Yepiskoposyan L, Tyler-Smith C, Xue Y, Abdullah MS, Ruiz-Linares A, Beall CM, Di Rienzo A, Jeong C, Starikovskaya EB, Metspalu E, Parik J, Villems R, Henn BM, Hodoglugil U, Mahley R, Sajantila A, Stamatoyannopoulos G, Wee JTS, Khusainova R, Khusnutdinova E, Litvinov S, Ayodo G, Comas D, Hammer MF, Kivisild T, Klitz W, Winkler CA, Labuda D, Bamshad M, Jorde LB, Tishkoff SA, Watkins WS, Metspalu M, Dryomov S, Sukernik R, Singh L, Thangaraj K, Pääbo S, Kelso J, Patterson N, Reich D. 2016. The Simons Genome Diversity Project: 300 genomes from 142 diverse populations. Nature 538:201–206.

Marbach D, Lamparter D, Quon G, Kellis M, Kutalik Z, Bergmann S. 2016. Tissue-specific regulatory circuits reveal variable modular perturbations across complex diseases. Nat Methods 13:366–370.

Maricic T, Günther V, Georgiev O, Gehre S, Curlin M, Schreiweis C, Naumann R, Burbano HA, Meyer M, Lalueza-Fox C, de la Rasilla M, Rosas A, Gajovic S, Kelso J, Enard W, Schaffner W, Pääbo S. 2013. A recent evolutionary change affects a regulatory element in the human FOXP2 gene. Mol Biol Evol 30:844–852.

Marnetto D, Pankratov V, Mondal M, Montinaro F, Pärna K, Vallini L, Molinaro L, Saag L, Loog L, Montagnese S, Costa R, Metspalu M, Eriksson A, Pagani L. n.d. Ancestral contributions to contemporary European complex traits. doi:10.1101/2021.08.03.454888

Mathelier A, Zhao X, Zhang AW, Parcy F, Worsley-Hunt R, Arenillas DJ, Buchman S, Chen CY, Chou A, Ienasescu H, Lim J, Shyr C, Tan G, Zhou M, Lenhard B, Sandelin A, Wasserman WW. 2014. JASPAR 2014: an extensively expanded and updated open-access database of transcription factor binding profiles. Nucleic Acids Res 42. doi:10.1093/nar/gkt997

Matys V, Kel-Margoulis OV, Fricke E, Liebich I, Land S, Barre-Dirrie A, Reuter I, Chekmenev D, Krull M, Hornischer K, Voss N, Stegmaier P, Lewicki-Potapov B, Saxel H, Kel AE, Wingender E. 2006. TRANSFAC and its module TRANSCompel: transcriptional gene regulation in eukaryotes. Nucleic Acids Res 34. doi:10.1093/nar/gkj143

McArthur E, Rinker DC, Capra JA. 2021. Quantifying the contribution of Neanderthal introgression to the heritability of complex traits. Nat Commun 12:4481.

McCoy RC, Wakefield J, Akey JM. 2017. Impacts of Neanderthal-Introgressed Sequences on the Landscape of Human Gene Expression. Cell 168:916–927.e12.

Mendez FL, Watkins JC, Hammer MF. 2012. A haplotype at STAT2 Introgressed from neanderthals and serves as a candidate of positive selection in Papua New Guinea. Am J Hum Genet 91:265–274.

Meyer M, Kircher M, Gansauge M-T, Li H, Racimo F, Mallick S, Schraiber JG, Jay F, Prüfer K, de Filippo C, Sudmant PH, Alkan C, Fu Q, Do R, Rohland N, Tandon A, Siebauer M, Green RE, Bryc K, Briggs AW, Stenzel U, Dabney J, Shendure J, Kitzman J, Hammer MF, Shunkov MV, Derevianko AP, Patterson N, Andrés AM, Eichler EE, Slatkin M, Reich D, Kelso J, Pääbo S. 2012. A high-coverage genome sequence from an archaic Denisovan individual. Science 338:222–226.

Ouwens KG, Jansen R, Nivard MG, van Dongen J, Frieser MJ, Hottenga J-J, Arindrarto W, Claringbould A, van Iterson M, Mei H, Franke L, Heijmans BT, A C’t Hoen P, van Meurs J, Brooks AI, BIOS Consortium, Penninx BWJH, Boomsma DI. 2020. A characterization of cis- and trans-heritability of RNA-Seq-based gene expression. Eur J Hum Genet 28:253–263.

Pankratov V, Montinaro F, Kushniarevich A, Hudjashov G, Jay F, Saag L, Flores R, Marnetto D, Seppel M, Kals M, Võsa U, Taccioli C, Möls M, Milani L, Aasa A, Lawson DJ, Esko T, Mägi R, Pagani L, Metspalu A, Metspalu M. 2020. Differences in local population history at the finest level: the case of the Estonian population. Eur J Hum Genet 28:1580–1591.

Perry GH, Dominy NJ, Claw KG, Lee AS, Fiegler H, Redon R, Werner J, Villanea FA, Mountain JL, Misra R, Carter NP, Lee C, Stone AC. 2007. Diet and the evolution of human amylase gene copy number variation. Nat Genet 39:1256–1260.

Petr M, Pääbo S, Kelso J, Vernot B. 2019. Limits of long-term selection against Neandertal introgression. Proc Natl Acad Sci U S A 116:1639–1644.

Peyrégne S, Boyle MJ, Dannemann M, Prüfer K. 2017. Detecting ancient positive selection in humans using extended lineage sorting. Genome Res 27:1563–1572.

Prüfer K, de Filippo C, Grote S, Mafessoni F, Korlević P, Hajdinjak M, Vernot B, Skov L, Hsieh P, Peyrégne S, Reher D, Hopfe C, Nagel S, Maricic T, Fu Q, Theunert C, Rogers R, Skoglund P, Chintalapati M, Dannemann M, Nelson BJ, Key FM, Rudan P, Kućan Ž, Gušić I, Golovanova LV, Doronichev VB, Patterson N, Reich D, Eichler EE, Slatkin M, Schierup MH, Andrés AM, Kelso J, Meyer M, Pääbo S. 2017. A high-coverage Neandertal genome from Vindija Cave in Croatia. Science 358:655–658.

Prüfer K, Racimo F, Patterson N, Jay F, Sankararaman S, Sawyer S, Heinze A, Renaud G, Sudmant PH, de Filippo C, Li H, Mallick S, Dannemann M, Fu Q, Kircher M, Kuhlwilm M, Lachmann M, Meyer M, Ongyerth M, Siebauer M, Theunert C, Tandon A, Moorjani P, Pickrell J, Mullikin JC, Vohr SH, Green RE, Hellmann I, Johnson PLF, Blanche H, Cann H, Kitzman JO, Shendure J, Eichler EE, Lein ES, Bakken TE, Golovanova LV, Doronichev VB, Shunkov MV, Derevianko AP, Viola B, Slatkin M, Reich D, Kelso J, Pääbo S. 2014. The complete genome sequence of a Neanderthal from the Altai Mountains. Nature 505:43–49.

Quach H, Rotival M, Pothlichet J, Loh Y-HE, Dannemann M, Zidane N, Laval G, Patin E, Harmant C, Lopez M, Deschamps M, Naffakh N, Duffy D, Coen A, Leroux-Roels G, Clément F, Boland A, Deleuze J-F, Kelso J, Albert ML, Quintana-Murci L. 2016. Genetic Adaptation and Neandertal Admixture Shaped the Immune System of Human Populations. Cell 167:643–656.e17.

Rao SSP, Huntley MH, Durand NC, Stamenova EK, Bochkov ID, Robinson JT, Sanborn AL, Machol I, Omer AD, Lander ES, Aiden EL. 2014. A 3D map of the human genome at kilobase resolution reveals principles of chromatin looping. Cell 159:1665–1680.

Sankararaman S, Mallick S, Patterson N, Reich D. 2016. The Combined Landscape of Denisovan and Neanderthal Ancestry in Present-Day Humans. Curr Biol 26:1241–1247.

Silvert M, Quintana-Murci L, Rotival M. 2019. Impact and Evolutionary Determinants of Neanderthal Introgression on Transcriptional and Post-Transcriptional Regulation. Am J Hum Genet 104:1241–1250.

Simonti CN, Vernot B, Bastarache L, Bottinger E, Carrell DS, Chisholm RL, Crosslin DR, Hebbring SJ, Jarvik GP, Kullo IJ, Li R, Pathak J, Ritchie MD, Roden DM, Verma SS, Tromp G, Prato JD, Bush WS, Akey JM, Denny JC, Capra JA. 2016. The phenotypic legacy of admixture between modern humans and Neandertals. Science 351:737–741.

Speidel L, Forest M, Shi S, Myers SR. 2019. A method for genome-wide genealogy estimation for thousands of samples. Nat Genet 51:1321–1329.

Stern AJ, Wilton PR, Nielsen R. 2019. An approximate full-likelihood method for inferring selection and allele frequency trajectories from DNA sequence data. PLoS Genet 15:e1008384.

Szklarczyk D, Gable AL, Lyon D, Junge A, Wyder S, Huerta-Cepas J, Simonovic M, Doncheva NT, Morris JH, Bork P, Jensen LJ, Mering C von. 2019. STRING v11: protein-protein association networks with increased coverage, supporting functional discovery in genome-wide experimental datasets. Nucleic Acids Res 47:D607–D613.

Telis N, Aguilar R, Harris K. 2020. Selection against archaic hominin genetic variation in regulatory regions. Nat Ecol Evol 4:1558–1566.

Vernot B, Tucci S, Kelso J, Schraiber JG, Wolf AB, Gittelman RM, Dannemann M, Grote S, McCoy RC, Norton H, Scheinfeldt LB, Merriwether DA, Koki G, Friedlaender JS, Wakefield J, Pääbo S, Akey JM. 2016. Excavating Neandertal and Denisovan DNA from the genomes of Melanesian individuals. Science 352:235–239.

Vespasiani DM, Jacobs GS, Brucato N, Cox MP, Romero IG. n.d. Denisovan introgression has shaped the immune system of present-day Papuans. doi:10.1101/2020.07.09.196444

Võsa U, Claringbould A, Westra H-J, Bonder MJ, Deelen P, Zeng B, Kirsten H, Saha A, Kreuzhuber R, Yazar S, Brugge H, Oelen R, de Vries DH, van der Wijst MGP, Kasela S, Pervjakova N, Alves I, Favé M-J, Agbessi M, Christiansen MW, Jansen R, Seppälä I, Tong L, Teumer A, Schramm K, Hemani G, Verlouw J, Yaghootkar H, Sönmez Flitman R, Brown A, Kukushkina V, Kalnapenkis A, Rüeger S, Porcu E, Kronberg J, Kettunen J, Lee B, Zhang F, Qi T, Hernandez JA, Arindrarto W, Beutner F, BIOS Consortium, i2QTL Consortium, Dmitrieva J, Elansary M, Fairfax BP, Georges M, Heijmans BT, Hewitt AW, Kähönen M, Kim Y, Knight JC, Kovacs P, Krohn K, Li S, Loeffler M, Marigorta UM, Mei H, Momozawa Y, Müller-Nurasyid M, Nauck M, Nivard MG, Penninx BWJH, Pritchard JK, Raitakari OT, Rotzschke O, Slagboom EP, Stehouwer CDA, Stumvoll M, Sullivan P, ’t Hoen PAC, Thiery J, Tönjes A, van Dongen J, van Iterson M, Veldink JH, Völker U, Warmerdam R, Wijmenga C, Swertz M, Andiappan A, Montgomery GW, Ripatti S, Perola M, Kutalik Z, Dermitzakis E, Bergmann S, Frayling T, van Meurs J, Prokisch H, Ahsan H, Pierce BL, Lehtimäki T, Boomsma DI, Psaty BM, Gharib SA, Awadalla P, Milani L, Ouwehand WH, Downes K, Stegle O, Battle A, Visscher PM, Yang J, Scholz M, Powell J, Gibson G, Esko T, Franke L. 2021. Large-scale cis- and trans-eQTL analyses identify thousands of genetic loci and polygenic scores that regulate blood gene expression. Nat Genet 53:1300–1310.

Xie X, Rigor P, Baldi P. 2009. MotifMap: a human genome-wide map of candidate regulatory motif sites. Bioinformatics 25. doi:10.1093/bioinformatics/btn605

Yang J, Benyamin B, McEvoy BP, Gordon S, Henders AK, Nyholt DR, Madden PA, Heath AC, Martin NG, Montgomery GW, Goddard ME, Visscher PM. 2010. Common SNPs explain a large proportion of the heritability for human height. Nat Genet 42:565–569.

Yao DW, O’Connor LJ, Price AL, Gusev A. 2020. Quantifying genetic effects on disease mediated by assayed gene expression levels. Nat Genet 52:626–633.

Zeberg H, Pääbo S. 2020. The major genetic risk factor for severe COVID-19 is inherited from Neanderthals. Nature 587:610–612.

