## Supplementary Tables for "Long-range regulatory effects of Neandertal DNA in modern humans"

**Table S1: eQTLGen trans-eQTLs (FDR<0.05) with aSNPs**

| Pvalue | SNP | SNPChr | SNPPos | AssessedAllele | OtherAllele | Zscore | Gene | GeneSymbol | GeneChr | GenePos | NrCohorts | NrSamples | FDR | BonferroniP |
|---|---|---|---|---|---|---|---|---|---|---|---|---|---|---|
| 2.95E-06 | rs72647484 | 1 | 22587728 | C | T | -4.6746 | ENSG00000184785 | SMIM10 | X | 134125734 | 14 | 8269 | 1.81E-02 | 1.00E+00 |
| 5.67E-06 | rs10919070 | 1 | 169099037 | C | A | 4.5383 | ENSG00000160563 | MED27 | 9 | 134845394 | 26 | 26022 | 3.11E-02 | 1.00E+00 |
| 3.75E-06 | rs10919071 | 1 | 169099483 | G | A | 4.6246 | ENSG00000160563 | MED27 | 9 | 134845394 | 25 | 25638 | 2.22E-02 | 1.00E+00 |
| 1.18E-12 | rs13063635 | 3 | 46173072 | C | G | -7.1075 | ENSG00000143365 | RORC | 1 | 151791447 | 36 | 31569 | 0.00E+00 | 2.40E-04 |
| 7.45E-08 | rs13063635 | 3 | 46173072 | C | G | -5.3799 | ENSG00000256553 | TRAV1-2 | 14 | 22111453 | 30 | 22495 | 8.00E-04 | 1.00E+00 |
| 1.07E-07 | rs13063635 | 3 | 46173072 | C | G | -5.3148 | ENSG00000113088 | GZMK | 5 | 54325239 | 36 | 31569 | 1.02E-03 | 1.00E+00 |
| 1.51E-07 | rs13063635 | 3 | 46173072 | C | G | -5.2515 | ENSG00000085563 | ABCB1 | 7 | 87237893 | 35 | 31529 | 1.35E-03 | 1.00E+00 |
| 4.00E-07 | rs13063635 | 3 | 46173072 | C | G | -5.0689 | ENSG00000111796 | KLRB1 | 12 | 9753814 | 35 | 31355 | 2.97E-03 | 1.00E+00 |
| 4.77E-07 | rs13063635 | 3 | 46173072 | C | G | -5.0354 | ENSG00000008256 | CYTH3 | 7 | 6256841 | 35 | 31355 | 3.56E-03 | 1.00E+00 |
| 2.21E-06 | rs13063635 | 3 | 46173072 | C | G | 4.7335 | ENSG00000136111 | TBC1D4 | 13 | 75957529 | 36 | 31569 | 1.41E-02 | 1.00E+00 |
| 3.84E-06 | rs13063635 | 3 | 46173072 | C | G | 4.62 | ENSG00000172005 | MAL | 2 | 95705579 | 35 | 31355 | 2.26E-02 | 1.00E+00 |
| 4.49E-06 | rs13063635 | 3 | 46173072 | C | G | -4.5872 | ENSG00000206561 | COLQ | 3 | 15527449 | 35 | 31355 | 2.55E-02 | 1.00E+00 |
| 5.20E-06 | rs13063635 | 3 | 46173072 | C | G | -4.5564 | ENSG00000144290 | SLC4A10 | 2 | 162561317 | 36 | 31569 | 2.89E-02 | 1.00E+00 |
| 7.00E-06 | rs13063635 | 3 | 46173072 | C | G | 4.4939 | ENSG00000113263 | ITK | 5 | 156626072 | 36 | 31569 | 3.69E-02 | 1.00E+00 |
| 5.39E-27 | rs13098911 | 3 | 46235201 | T | C | -10.7586 | ENSG00000143365 | RORC | 1 | 151791447 | 35 | 31185 | 0.00E+00 | 1.10E-18 |
| 3.05E-15 | rs13098911 | 3 | 46235201 | T | C | -7.8887 | ENSG00000111796 | KLRB1 | 12 | 9753814 | 34 | 30971 | 0.00E+00 | 6.21E-07 |
| 6.02E-15 | rs13098911 | 3 | 46235201 | T | C | -7.8036 | ENSG00000113088 | GZMK | 5 | 54325239 | 35 | 31185 | 0.00E+00 | 1.22E-06 |
| 3.91E-14 | rs13098911 | 3 | 46235201 | T | C | -7.5637 | ENSG00000256553 | TRAV1-2 | 14 | 22111453 | 29 | 22111 | 0.00E+00 | 7.97E-06 |
| 5.97E-12 | rs13098911 | 3 | 46235201 | T | C | -6.8802 | ENSG00000085563 | ABCB1 | 7 | 87237893 | 34 | 31145 | 0.00E+00 | 1.21E-03 |
| 1.47E-11 | rs13098911 | 3 | 46235201 | T | C | -6.7503 | ENSG00000008256 | CYTH3 | 7 | 6256841 | 34 | 30971 | 0.00E+00 | 3.00E-03 |
| 6.71E-11 | rs13098911 | 3 | 46235201 | T | C | -6.5268 | ENSG00000206561 | COLQ | 3 | 15527449 | 34 | 30971 | 0.00E+00 | 1.37E-02 |
| 2.00E-10 | rs13098911 | 3 | 46235201 | T | C | -6.3608 | ENSG00000062524 | LTK | 15 | 41800960 | 35 | 31185 | 0.00E+00 | 4.08E-02 |
| 4.80E-10 | rs13098911 | 3 | 46235201 | T | C | -6.2252 | ENSG00000118402 | ELOVL4 | 6 | 80640913 | 35 | 31185 | 1.76E-05 | 9.78E-02 |
| 6.97E-10 | rs13098911 | 3 | 46235201 | T | C | -6.1668 | ENSG00000144290 | SLC4A10 | 2 | 162561317 | 35 | 31185 | 1.71E-05 | 1.42E-01 |
| 1.93E-09 | rs13098911 | 3 | 46235201 | T | C | -6.0037 | ENSG00000137501 | SYTL2 | 11 | 85463725 | 34 | 31145 | 4.72E-05 | 3.92E-01 |
| 7.84E-09 | rs13098911 | 3 | 46235201 | T | C | -5.7716 | ENSG00000171735 | CAMTA1 | 1 | 7337575 | 35 | 31185 | 1.26E-04 | 1.00E+00 |
| 1.69E-08 | rs13098911 | 3 | 46235201 | T | C | -5.6405 | ENSG00000119537 | KDSR | 18 | 61014851 | 34 | 30971 | 1.96E-04 | 1.00E+00 |
| 2.05E-08 | rs13098911 | 3 | 46235201 | T | C | -5.6078 | ENSG00000152268 | SPON1 | 11 | 14136780 | 35 | 31185 | 2.05E-04 | 1.00E+00 |
| 2.68E-08 | rs13098911 | 3 | 46235201 | T | C | -5.5613 | ENSG00000197635 | **DPP4** | 2 | 162889901 | 35 | 31185 | 3.01E-04 | 1.00E+00 |
| 3.54E-08 | rs13098911 | 3 | 46235201 | T | C | -5.5125 | ENSG00000112419 | PHACTR2 | 6 | 144005152 | 35 | 31185 | 3.88E-04 | 1.00E+00 |
| 5.66E-08 | rs13098911 | 3 | 46235201 | T | C | -5.4291 | ENSG00000146250 | PRSS35 | 6 | 84228808 | 35 | 31185 | 6.71E-04 | 1.00E+00 |
| 8.81E-08 | rs13098911 | 3 | 46235201 | T | C | 5.35 | ENSG00000138795 | LEF1 | 4 | 109029406 | 34 | 30971 | 8.74E-04 | 1.00E+00 |
| 9.74E-08 | rs13098911 | 3 | 46235201 | T | C | -5.3315 | ENSG00000157985 | AGAP1 | 2 | 236718965 | 34 | 30971 | 9.29E-04 | 1.00E+00 |
| 1.55E-07 | rs13098911 | 3 | 46235201 | T | C | -5.2461 | ENSG00000182472 | CAPN12 | 19 | 39240685 | 34 | 31145 | 1.35E-03 | 1.00E+00 |
| 1.59E-07 | rs13098911 | 3 | 46235201 | T | C | -5.2413 | ENSG00000157111 | TMEM171 | 5 | 72421881 | 35 | 31185 | 1.36E-03 | 1.00E+00 |
| 4.26E-07 | rs13098911 | 3 | 46235201 | T | C | -5.0566 | ENSG00000218357 | LL22NC03-75H12.2 | 22 | 47869954 | 20 | 10736 | 3.24E-03 | 1.00E+00 |

**Table S1: eQTLGen trans-eQTLs (FDR<0.05) with aSNPs**

| Pvalue | SNP | SNPChr | SNPPos | AssessedAllele | OtherAllele | Zscore | Gene | GeneSymbol | GeneChr | GenePos | NrCohorts | NrSamples | FDR | BonferroniP |
|---|---|---|---|---|---|---|---|---|---|---|---|---|---|---|
| 4.85E-07 | rs13098911 | 3 | 46235201 | T | C | 5.0321 | ENSG00000136111 | TBC1D4 | 13 | 75957529 | 35 | 31185 | 3.62E-03 | 1.00E+00 |
| 7.96E-07 | rs13098911 | 3 | 46235201 | T | C | -4.9364 | ENSG00000162630 | B3GALT2 | 1 | 193151979 | 35 | 31185 | 5.68E-03 | 1.00E+00 |
| 9.70E-07 | rs13098911 | 3 | 46235201 | T | C | -4.8974 | ENSG00000117090 | SLAMF1 | 1 | 160597487 | 35 | 31185 | 6.71E-03 | 1.00E+00 |
| 1.66E-06 | rs13098911 | 3 | 46235201 | T | C | 4.791 | ENSG00000179715 | PCED1B | 12 | 47551914 | 33 | 25191 | 1.09E-02 | 1.00E+00 |
| 2.34E-06 | rs13098911 | 3 | 46235201 | T | C | -4.7217 | ENSG00000156787 | WDR67 | 8 | 124109300 | 34 | 26110 | 1.48E-02 | 1.00E+00 |
| 2.38E-06 | rs13098911 | 3 | 46235201 | T | C | -4.7179 | ENSG00000130649 | CYP2E1 | 10 | 135354317 | 35 | 31185 | 1.50E-02 | 1.00E+00 |
| 2.51E-06 | rs13098911 | 3 | 46235201 | T | C | -4.7075 | ENSG00000158321 | AUTS2 | 7 | 69660979 | 35 | 31185 | 1.56E-02 | 1.00E+00 |
| 3.55E-06 | rs13098911 | 3 | 46235201 | T | C | 4.636 | ENSG00000120915 | EPHX2 | 8 | 27375688 | 35 | 31185 | 2.12E-02 | 1.00E+00 |
| 3.67E-06 | rs13098911 | 3 | 46235201 | T | C | 4.6294 | ENSG00000113263 | ITK | 5 | 156626072 | 35 | 31185 | 2.18E-02 | 1.00E+00 |
| 4.37E-06 | rs13098911 | 3 | 46235201 | T | C | -4.5927 | ENSG00000080546 | SESN1 | 6 | 109361831 | 35 | 31185 | 2.50E-02 | 1.00E+00 |
| 4.97E-06 | rs13098911 | 3 | 46235201 | T | C | -4.566 | ENSG00000008283 | CYB561 | 17 | 61516702 | 34 | 30971 | 2.78E-02 | 1.00E+00 |
| 4.83E-07 | rs6784615 | 3 | 52506426 | C | T | 5.033 | ENSG00000205269 | TMEM170B | 6 | 11560847 | 25 | 20622 | 3.60E-03 | 1.00E+00 |
| 8.12E-06 | rs56032325 | 6 | 52281072 | A | T | -4.4618 | ENSG00000250366 | LINC00617 | 14 | 96367318 | 13 | 5502 | 4.17E-02 | 1.00E+00 |
| 1.29E-06 | rs7811653 | 7 | 46402269 | A | C | -4.8404 | ENSG00000159873 | CCDC117 | 22 | 29176972 | 36 | 31644 | 8.69E-03 | 1.00E+00 |
| 4.40E-06 | rs72996113 | 11 | 100453046 | T | C | 4.5916 | ENSG00000211852 | TRAJ37 | 14 | 22972766 | 9 | 3831 | 2.51E-02 | 1.00E+00 |
| 1.41E-30 | rs2066807 | 12 | 56740682 | G | C | -11.4943 | ENSG00000265688 | MAFG-AS1 | 17 | 79887167 | 19 | 14631 | 0.00E+00 | 2.87E-22 |
| 5.28E-06 | rs2066807 | 12 | 56740682 | G | C | -4.5533 | ENSG00000163565 | IFI16 | 1 | 158997351 | 35 | 31086 | 2.93E-02 | 1.00E+00 |
| 4.35E-28 | rs2066819 | 12 | 56750204 | T | C | -10.9883 | ENSG00000265688 | MAFG-AS1 | 17 | 79887167 | 19 | 14629 | 0.00E+00 | 8.85E-20 |
| 1.19E-06 | rs12908161 | 15 | 85207825 | G | A | -4.8576 | ENSG00000183625 | CCR3 | 3 | 46256646 | 35 | 31355 | 7.97E-03 | 1.00E+00 |
| 2.46E-06 | rs12908161 | 15 | 85207825 | G | A | -4.7117 | ENSG00000165046 | LETM2 | 8 | 38255385 | 16 | 14263 | 1.53E-02 | 1.00E+00 |
| 5.71E-06 | rs12908161 | 15 | 85207825 | G | A | -4.5369 | ENSG00000101460 | MAP1LC3A | 20 | 33141403 | 35 | 31355 | 3.13E-02 | 1.00E+00 |
| 7.81E-06 | rs12603526 | 17 | 800593 | C | T | -4.47 | ENSG00000156697 | UTP14A | X | 129051917 | 14 | 8997 | 4.04E-02 | 1.00E+00 |
| 1.30E-07 | rs11650665 | 17 | 76259847 | G | T | 5.2783 | ENSG00000204435 | CSNK2B | 6 | 31635566 | 15 | 11496 | 1.17E-03 | 1.00E+00 |
| 3.37E-10 | rs72973711 | 18 | 74072245 | T | A | 6.2806 | ENSG00000167600 | CYP2S1 | 19 | 41706199 | 32 | 30031 | 1.81E-05 | 6.87E-02 |
| 3.52E-09 | rs72973711 | 18 | 74072245 | T | A | -5.905 | ENSG00000214688 | C10orf105 | 10 | 73484519 | 20 | 16642 | 8.98E-05 | 7.17E-01 |
| 2.01E-08 | rs72973711 | 18 | 74072245 | T | A | -5.6108 | ENSG00000136999 | NOV | 8 | 120432569 | 32 | 30031 | 2.06E-04 | 1.00E+00 |
| 2.01E-08 | rs72973711 | 18 | 74072245 | T | A | -5.6107 | ENSG00000138613 | APH1B | 15 | 63584771 | 32 | 30031 | 2.06E-04 | 1.00E+00 |
| 2.72E-08 | rs72973711 | 18 | 74072245 | T | A | 5.5586 | ENSG00000186529 | CYP4F3 | 19 | 15762670 | 30 | 29932 | 3.25E-04 | 1.00E+00 |
| 4.26E-08 | rs72973711 | 18 | 74072245 | T | A | -5.4795 | ENSG00000134755 | DSC2 | 18 | 28664159 | 31 | 29817 | 4.99E-04 | 1.00E+00 |
| 5.57E-08 | rs72973711 | 18 | 74072245 | T | A | 5.432 | ENSG00000158470 | B4GALT5 | 20 | 48289948 | 31 | 29817 | 6.61E-04 | 1.00E+00 |
| 9.53E-08 | rs72973711 | 18 | 74072245 | T | A | 5.3356 | ENSG00000164850 | GPER | 7 | 1127647 | 31 | 24956 | 9.09E-04 | 1.00E+00 |
| 9.61E-08 | rs72973711 | 18 | 74072245 | T | A | 5.334 | ENSG00000131467 | PSME3 | 17 | 40986088 | 32 | 30031 | 9.09E-04 | 1.00E+00 |
| 1.66E-07 | rs72973711 | 18 | 74072245 | T | A | -5.2344 | ENSG00000165092 | ALDH1A1 | 9 | 75605468 | 31 | 29991 | 1.38E-03 | 1.00E+00 |
| 2.40E-07 | rs72973711 | 18 | 74072245 | T | A | 5.1653 | ENSG00000088766 | CRLS1 | 20 | 6003717 | 32 | 30031 | 1.94E-03 | 1.00E+00 |
| 2.89E-07 | rs72973711 | 18 | 74072245 | T | A | -5.1304 | ENSG00000152784 | PRDM8 | 4 | 81115258 | 32 | 30031 | 2.30E-03 | 1.00E+00 |
| 7.57E-07 | rs72973711 | 18 | 74072245 | T | A | -4.946 | ENSG00000154330 | PGM5 | 9 | 71058896 | 32 | 30031 | 5.43E-03 | 1.00E+00 |

**Table S1: eQTLGen trans-eQTLs (FDR<0.05) with aSNPs**

| Pvalue | SNP | SNPChr | SNPPos | AssessedAllele | OtherAllele | Zscore | Gene | GeneSymbol | GeneChr | GenePos | NrCohorts | NrSamples | FDR | BonferroniP |
|---|---|---|---|---|---|---|---|---|---|---|---|---|---|---|
| 7.85E-07 | rs72973711 | 18 | 74072245 | T | A | 4.9391 | ENSG00000159423 | ALDH4A1 | 1 | 19213600 | 29 | 24702 | 5.63E-03 | 1.00E+00 |
| 1.23E-06 | rs72973711 | 18 | 74072245 | T | A | 4.8513 | ENSG00000125810 | CD93 | 20 | 23063481 | 32 | 30031 | 8.26E-03 | 1.00E+00 |
| 1.39E-06 | rs72973711 | 18 | 74072245 | T | A | 4.8262 | ENSG00000070404 | FSTL3 | 19 | 679888 | 32 | 30031 | 9.26E-03 | 1.00E+00 |
| 1.47E-06 | rs72973711 | 18 | 74072245 | T | A | 4.815 | ENSG00000113161 | HMGCR | 5 | 74645041 | 32 | 30031 | 9.74E-03 | 1.00E+00 |
| 1.92E-06 | rs72973711 | 18 | 74072245 | T | A | 4.7613 | ENSG00000121316 | PLBD1 | 12 | 14688939 | 30 | 24742 | 1.25E-02 | 1.00E+00 |
| 1.93E-06 | rs72973711 | 18 | 74072245 | T | A | 4.7603 | ENSG00000138834 | MAPK8IP3 | 16 | 1788251 | 32 | 30031 | 1.26E-02 | 1.00E+00 |
| 3.36E-06 | rs72973711 | 18 | 74072245 | T | A | 4.6475 | ENSG00000183307 | CECR6 | 22 | 17599723 | 32 | 30031 | 2.01E-02 | 1.00E+00 |
| 3.91E-06 | rs72973711 | 18 | 74072245 | T | A | -4.616 | ENSG00000115339 | GALNT3 | 2 | 166627646 | 14 | 13438 | 2.29E-02 | 1.00E+00 |
| 4.21E-06 | rs72973711 | 18 | 74072245 | T | A | 4.6009 | ENSG00000135604 | STX11 | 6 | 144490585 | 32 | 30031 | 2.43E-02 | 1.00E+00 |
| 4.43E-06 | rs72973711 | 18 | 74072245 | T | A | 4.5903 | ENSG00000196562 | SULF2 | 20 | 46350226 | 31 | 29817 | 2.53E-02 | 1.00E+00 |
| 5.83E-06 | rs72973711 | 18 | 74072245 | T | A | 4.5324 | ENSG00000184489 | PTP4A3 | 8 | 142421856 | 31 | 29817 | 3.19E-02 | 1.00E+00 |
| 6.94E-06 | rs72973711 | 18 | 74072245 | T | A | 4.4958 | ENSG00000011198 | ABHD5 | 3 | 43753734 | 32 | 30031 | 3.67E-02 | 1.00E+00 |
| 7.58E-06 | rs72973711 | 18 | 74072245 | T | A | 4.4767 | ENSG00000183019 | C19orf59 | 19 | 7743113 | 31 | 24956 | 3.94E-02 | 1.00E+00 |
| 8.18E-06 | rs72973711 | 18 | 74072245 | T | A | -4.4602 | ENSG00000071967 | CYBRD1 | 2 | 172396700 | 31 | 29817 | 4.19E-02 | 1.00E+00 |
| 1.62E-06 | rs4805834 | 19 | 33453659 | T | C | -4.7953 | ENSG00000187735 | TCEA1 | 8 | 54907100 | 14 | 6421 | 1.07E-02 | 1.00E+00 |
| 5.12E-07 | rs16997087 | 20 | 16054982 | C | T | 5.0217 | ENSG00000142856 | ITGB3BP | 1 | 63982916 | 16 | 14263 | 3.74E-03 | 1.00E+00 |
| 9.31E-15 | rs13043612 | 20 | 4132364 | G | C | -7.7482 | ENSG00000022840 | RNF10 | 12 | 120993340 | 37 | 31684 | 0.00E+00 | 1.89E-06 |
| 1.67E-10 | rs13043612 | 20 | 4132364 | G | C | -6.3891 | ENSG00000136732 | GYPC | 2 | 127433877 | 37 | 31684 | 0.00E+00 | 3.39E-02 |
| 4.11E-10 | rs13043612 | 20 | 4132364 | G | C | -6.2498 | ENSG00000167671 | UBXN6 | 19 | 4451407 | 37 | 31684 | 1.78E-05 | 8.37E-02 |
| 2.25E-09 | rs13043612 | 20 | 4132364 | G | C | 5.9784 | ENSG00000172331 | BPGM | 7 | 134348062 | 37 | 31684 | 6.22E-05 | 4.59E-01 |
| 2.82E-09 | rs13043612 | 20 | 4132364 | G | C | 5.9419 | ENSG00000143995 | MEIS1 | 2 | 66730792 | 36 | 31644 | 6.11E-05 | 5.74E-01 |
| 9.72E-09 | rs13043612 | 20 | 4132364 | G | C | -5.7355 | ENSG00000100325 | ASCC2 | 22 | 30209434 | 37 | 31684 | 1.37E-04 | 1.00E+00 |
| 9.87E-09 | rs13043612 | 20 | 4132364 | G | C | 5.7331 | ENSG00000196517 | SLC6A9 | 1 | 44477155 | 37 | 31684 | 1.37E-04 | 1.00E+00 |
| 1.01E-07 | rs13043612 | 20 | 4132364 | G | C | 5.3249 | ENSG00000169313 | P2RY12 | 3 | 151078884 | 35 | 31430 | 9.48E-04 | 1.00E+00 |
| 3.91E-07 | rs13043612 | 20 | 4132364 | G | C | -5.0733 | ENSG00000159346 | ADIPOR1 | 1 | 202918825 | 36 | 31644 | 2.91E-03 | 1.00E+00 |
| 1.10E-06 | rs13043612 | 20 | 4132364 | G | C | 4.8722 | ENSG00000146007 | ZMAT2 | 5 | 140082256 | 37 | 31684 | 7.49E-03 | 1.00E+00 |
| 5.28E-06 | rs13043612 | 20 | 4132364 | G | C | -4.5534 | ENSG00000163374 | YY1AP1 | 1 | 155644014 | 37 | 31684 | 2.93E-02 | 1.00E+00 |
| 7.24E-06 | rs13043612 | 20 | 4132364 | G | C | -4.4863 | ENSG00000161203 | AP2M1 | 3 | 183897178 | 36 | 31470 | 3.78E-02 | 1.00E+00 |
| 7.64E-06 | rs13043612 | 20 | 4132364 | G | C | -4.4748 | ENSG00000100104 | SRRD | 22 | 26883871 | 37 | 31684 | 3.97E-02 | 1.00E+00 |

**Table S2: GTEx cis-eQTLs (FDR<0.05) between aSNPs and TFs**

| phenotype_id | num_var | beta_shape1 | beta_shape2 | true_df | pval_true_df | variant_id | tss_distance | ma_samples | ma_count | maf | ref_factor | pval_nominal | slope | slope_se | pval_perm | pval_beta | qval | pval_nominal_threshold | tissue | gene | hg19 |
|---|---|---|---|---|---|---|---|---|---|---|---|---|---|---|---|---|---|---|---|---|---|
| ENSG00000171786.5 | 6200 | 1.03 | 784.827 | 107.587 | 1.79622e-05 | chr1_159918911_C_T_b38 | -448156 | 8 | 9 | 0.0288462 | 1 | 8.42E-06 | 0.823796 | 0.176694 | 1.05E-02 | 1.21E-02 | 3.23E-02 | 3.14E-05 | Brain_Hypothalamus | NHLH1 | 1_159888701 |
| ENSG00000162761.14 | 6290 | 1.04 | 808.636 | 247.267 | 9.98117e-06 | chr1_165347134_G_A_b38 | -9581 | 61 | 67 | 0.101824 | 1 | 3.31E-06 | 0.326767 | 0.0688148 | 5.70E-03 | 6.48E-03 | 9.93E-03 | 6.34E-05 | Breast_Mammary_Tissue | LMX1A | 1_165316371 |
| ENSG00000116132.11 | 6932 | 1.02 | 471 | 118.65 | 6.58725e-05 | chr1_170665754_T_A_b38 | 3026 | 36 | 40 | 0.116279 | 1 | 2.55E-05 | 0.249672 | 0.0572086 | 2.60E-02 | 2.80E-02 | 4.31E-02 | 7.94E-05 | Brain_Caudate_basal_ganglia | PRRX1 | 1_170634895 |
| ENSG00000117222.13 | 6023 | 1.04 | 812.096 | 106.553 | 3.67382e-06 | chr1_205487623_G_A_b38 | 365608 | 19 | 19 | 0.0608974 | 1 | 1.36E-06 | -0.55838 | 0.109569 | 1.40E-03 | 2.29E-03 | 8.19E-03 | 3.18E-05 | Brain_Hypothalamus | RBBP5 | 1_205456751 |
| ENSG00000117707.15 | 5856 | 1.04 | 678.208 | 139.708 | 3.59537e-05 | chr1_213297057_A_G_b38 | -686124 | 20 | 20 | 0.0515464 | 1 | 1.43E-05 | 0.62754 | 0.139992 | 1.77E-02 | 2.00E-02 | 3.54E-02 | 5.44E-05 | Adrenal_Gland | PROX1 | 1_213470400 |
| ENSG00000187801.14 | 6134 | 1.06 | 666.947 | 102.18 | 4.8666e-05 | chr1_40881914_T_C_b38 | 431807 | 21 | 21 | 0.0686275 | 1 | 1.93E-05 | -0.603085 | 0.135172 | 2.21E-02 | 2.56E-02 | 4.59E-02 | 5.41E-05 | Brain_Putamen_basal_ganglia | ZFP69B | 1_41347586 |
| ENSG00000177606.6 | 6082 | 1.06 | 680.64 | 130.112 | 1.92314e-05 | chr1_58434644_C_A_b38 | -349683 | 74 | 81 | 0.221311 | 1 | 7.43E-06 | 0.313442 | 0.0673826 | 9.80E-03 | 9.80E-03 | 1.64E-02 | 7.01E-05 | Brain_Cortex | JUN | 1_58900316 |
| ENSG00000179774.8 | 6064 | 1.06 | 340.29 | 135.106 | 2.21761e-16 | chr10_68268558_T_C_b38 | 36455 | 67 | 79 | 0.203608 | 1 | 1.84E-18 | -0.754614 | 0.0754373 | 1.00E-04 | 1.22E-14 | 1.52E-13 | 1.15E-04 | Adrenal_Gland | ATOH7 | 10_70028315 |
| ENSG00000107485.15 | 8090 | 1.05 | 1088.91 | 371.177 | 3.21359e-13 | chr10_8519244_C_T_b38 | 465640 | 136 | 152 | 0.157676 | 1 | 1.64E-14 | -0.484609 | 0.0608522 | 1.00E-04 | 1.05E-10 | 1.39E-10 | 1.22E-04 | Thyroid | GATA3 | 10_8561207 |
| ENSG00000082175.14 | 7313 | 1.05 | 551.049 | 184.962 | 1.04475e-05 | chr11_101262781_C_G_b38 | 132257 | 78 | 88 | 0.165414 | 1 | 2.50E-06 | 0.221429 | 0.0457411 | 3.40E-03 | 4.37E-03 | 7.04E-03 | 9.42E-05 | Colon_Sigmoid | PGR | 11_101133512 |
| ENSG00000166478.9 | 6319 | 1.03 | 403.288 | 276.261 | 1.36453e-06 | chr11_9388086_G_T_b38 | -72617 | 46 | 49 | 0.062982 | 1 | 1.76E-07 | 0.334113 | 0.0625682 | 6.00E-04 | 4.36E-04 | 6.18E-04 | 1.67E-04 | Adipose_Visceral_Omentum | ZNF143 | 11_9409633 |
| ENSG00000166478.9 | 6319 | 1.03 | 447.279 | 239.12 | 9.31049e-08 | chr11_9388086_G_T_b38 | -72617 | 40 | 40 | 0.0615385 | 1 | 1.23E-08 | 0.455444 | 0.0775158 | 1.00E-04 | 3.03E-05 | 6.47E-05 | 1.23E-04 | Heart_Left_Ventricle | ZNF143 | 11_9409633 |
| ENSG00000166478.9 | 6319 | 1.05 | 399.84 | 289.219 | 1.28445e-12 | chr11_9410543_C_T_b38 | -50160 | 50 | 53 | 0.0644769 | 1 | 1.30E-14 | 0.502761 | 0.0623744 | 1.00E-04 | 1.88E-10 | 3.62E-10 | 2.51E-04 | Esophagus_Mucosa | ZNF143 | 11_9432090 |
| ENSG00000166478.9 | 6319 | 1.04 | 459.851 | 104.178 | 9.31955e-08 | chr11_9459030_G_A_b38 | -1673 | 20 | 22 | 0.0700637 | 1 | 1.51E-08 | 0.674272 | 0.110809 | 1.00E-04 | 2.82E-05 | 7.21E-05 | 1.15E-04 | Brain_Cerebellar_Hemisphere | ZNF143 | 11_9480577 |
| ENSG00000166478.9 | 6319 | 1.05 | 411.961 | 224.055 | 4.86727e-07 | chr11_9465700_C_T_b38 | 4997 | 48 | 52 | 0.0822785 | 1 | 5.59E-08 | 0.386143 | 0.0690271 | 2.00E-04 | 1.23E-04 | 2.14E-04 | 1.62E-04 | Heart_Atrial_Appendage | ZNF143 | 11_9487247 |
| ENSG00000166478.9 | 6319 | 1.02 | 485.319 | 105.361 | 1.8352e-07 | chr11_9465700_C_T_b38 | 4997 | 24 | 24 | 0.0851064 | 1 | 4.41E-08 | 0.793329 | 0.135366 | 1.00E-04 | 7.37E-05 | 4.49E-04 | 4.44E-05 | Small_Intestine_Terminal_Ileum | ZNF143 | 11_9487247 |
| ENSG00000166478.9 | 6319 | 1.04 | 405.172 | 415.845 | 1.30246e-14 | chr11_9465700_C_T_b38 | 4997 | 82 | 88 | 0.078853 | 1 | 6.94E-17 | 0.309939 | 0.0357901 | 1.00E-04 | 2.02E-12 | 3.63E-12 | 2.65E-04 | Whole_Blood | ZNF143 | 11_9487247 |
| ENSG00000166478.9 | 6319 | 1.03 | 410.199 | 206.505 | 2.05753e-07 | chr11_9482935_T_G_b38 | 22232 | 39 | 40 | 0.0680272 | 1 | 2.29E-08 | 0.294639 | 0.0509513 | 1.00E-04 | 6.26E-05 | 1.34E-04 | 1.33E-04 | Colon_Transverse | ZNF143 | 11_9504482 |
| ENSG00000166478.9 | 6319 | 1.06 | 391.495 | 185.885 | 2.58474e-08 | chr11_9482935_T_G_b38 | 22232 | 35 | 36 | 0.0654545 | 1 | 1.38E-09 | 0.456593 | 0.0721608 | 1.00E-04 | 4.70E-06 | 1.19E-05 | 1.47E-04 | Esophagus_Gastroesophageal_Junction | ZNF143 | 11_9504482 |
| ENSG00000166478.9 | 6319 | 1.05 | 412.804 | 270.293 | 1.58518e-11 | chr11_9482935_T_G_b38 | 22232 | 52 | 54 | 0.0701299 | 1 | 3.42E-13 | 0.426106 | 0.0560653 | 1.00E-04 | 2.55E-09 | 4.65E-09 | 2.33E-04 | Esophagus_Muscularis | ZNF143 | 11_9504482 |
| ENSG00000166478.9 | 6319 | 1.01 | 420.182 | 356.248 | 3.32635e-32 | chr11_9482935_T_G_b38 | 22232 | 69 | 71 | 0.0736515 | 1 | 5.52E-37 | 0.814631 | 0.0579077 | 1.00E-04 | 7.81E-30 | 2.94E-29 | 2.79E-04 | Thyroid | ZNF143 | 11_9504482 |
| ENSG00000166478.9 | 6319 | 1.06 | 411.586 | 313.718 | 7.50415e-17 | chr11_9486860_G_A_b38 | 26157 | 56 | 59 | 0.0673516 | 1 | 1.68E-19 | 0.553033 | 0.0578235 | 1.00E-04 | 4.99E-15 | 9.24E-15 | 3.29E-04 | Nerve_Tibial | ZNF143 | 11_9508407 |
| ENSG00000166478.9 | 6319 | 1.03 | 471.033 | 124.608 | 1.73803e-22 | chr11_9486860_G_A_b38 | 26157 | 25 | 25 | 0.0698324 | 1 | 6.62E-25 | 1.20805 | 0.0954384 | 1.00E-04 | 2.63E-20 | 4.35E-19 | 1.04E-04 | Spleen | ZNF143 | 11_9508407 |
| ENSG00000166478.9 | 6319 | 1.05 | 415.717 | 235.758 | 3.53097e-14 | chr11_9487805_G_C_b38 | 27102 | 46 | 48 | 0.0729483 | 1 | 3.09E-16 | 0.646699 | 0.0743132 | 1.00E-04 | 4.61E-12 | 1.57E-11 | 1.74E-04 | Artery_Aorta | ZNF143 | 11_9509352 |
| ENSG00000166478.9 | 6319 | 1.06 | 419.935 | 116.337 | 3.7455e-09 | chr11_9494639_T_C_b38 | 33936 | 30 | 33 | 0.0942857 | 1 | 2.12E-10 | 0.531511 | 0.0773434 | 1.00E-04 | 6.52E-07 | 4.86E-06 | 6.44E-05 | Artery_Coronary | ZNF143 | 11_9516186 |
| ENSG00000166478.9 | 6319 | 1.04 | 449.313 | 130.54 | 3.97984e-06 | chr11_9515452_G_A_b38 | 54749 | 22 | 23 | 0.0611702 | 1 | 9.00E-07 | 0.617646 | 0.120428 | 8.00E-04 | 1.36E-03 | 2.01E-03 | 1.40E-04 | Brain_Cerebellum | ZNF143 | 11_9536999 |
| ENSG00000166478.9 | 6319 | 1.06 | 396.149 | 153.704 | 2.53624e-06 | chr11_9525993_T_C_b38 | 65290 | 25 | 28 | 0.0639269 | 1 | 3.80E-07 | 0.619264 | 0.117392 | 3.00E-04 | 6.57E-04 | 1.50E-03 | 1.16E-04 | Pituitary | ZNF143 | 11_9547540 |
| ENSG00000196387.9 | 4717 | 1.03 | 477.694 | 117.042 | 1.31806e-05 | chr12_133173626_C_T_b38 | 93788 | 70 | 78 | 0.226744 | 1 | 3.69E-06 | 0.331329 | 0.0685658 | 4.60E-03 | 5.31E-03 | 1.08E-02 | 8.05E-05 | Brain_Caudate_basal_ganglia | ZNF140 | 12_133750212 |
| ENSG00000256223.5 | 4599 | 1.03 | 479.695 | 180.684 | 1.838e-06 | chr12_133231543_C_T_b38 | 100968 | 114 | 133 | 0.255769 | 1 | 3.73E-07 | -0.186946 | 0.0355867 | 4.00E-04 | 7.03E-04 | 1.78E-03 | 7.73E-05 | Stomach | ZNF10 | 12_133808129 |
| ENSG00000170581.13 | 3991 | 1.03 | 487.906 | 108.19 | 1.03015e-08 | chr12_56259776_A_G_b38 | -100350 | 31 | 33 | 0.105096 | 1 | 2.60E-09 | 0.520675 | 0.0807006 | 1.00E-04 | 3.44E-06 | 1.02E-05 | 1.05E-04 | Brain_Cerebellar_Hemisphere | STAT2 | 12_56653560 |
| ENSG00000170581.13 | 3991 | 1.03 | 440.508 | 326.546 | 3.60545e-17 | chr12_56259776_A_G_b38 | -100350 | 73 | 78 | 0.0890411 | 1 | 3.73E-19 | -0.427451 | 0.0451825 | 1.00E-04 | 7.09E-15 | 1.30E-14 | 2.83E-04 | Nerve_Tibial | STAT2 | 12_56653560 |
| ENSG00000170581.13 | 3991 | 1.05 | 461.509 | 319.377 | 3.44783e-05 | chr12_56259776_A_G_b38 | -100350 | 60 | 64 | 0.0744186 | 1 | 1.09E-05 | -0.247727 | 0.0555383 | 1.19E-02 | 1.25E-02 | 9.87E-03 | 2.09E-04 | Skin_Not_Sun_Exposed_Suprapubic | STAT2 | 12_56653560 |
| ENSG00000170581.13 | 3991 | 1.03 | 457.418 | 124.688 | 1.20394e-18 | chr12_56302761_T_G_b38 | -57365 | 17 | 18 | 0.0545455 | 1 | 1.78E-20 | 1.01551 | 0.0927267 | 1.00E-04 | 1.79E-16 | 4.40E-15 | 5.31E-05 | Liver | STAT2 | 12_56696545 |
| ENSG00000170581.13 | 3991 | 1.04 | 480.23 | 367.982 | 3.48106e-19 | chr12_56352774_C_T_b38 | -7352 | 73 | 76 | 0.0793319 | 1 | 3.77E-21 | -0.47175 | 0.047253 | 1.00E-04 | 3.30E-17 | 9.16E-17 | 2.23E-04 | Adipose_Subcutaneous | STAT2 | 12_56746558 |
| ENSG00000170581.13 | 3991 | 1.04 | 464.338 | 288.471 | 1.19561e-09 | chr12_56352774_C_T_b38 | -7352 | 56 | 57 | 0.0725191 | 1 | 1.23E-10 | -0.392885 | 0.0590595 | 1.00E-04 | 3.19E-07 | 6.89E-07 | 1.49E-04 | Adipose_Visceral_Omentum | STAT2 | 12_56746558 |
| ENSG00000170581.13 | 3991 | 1.04 | 462.085 | 361.859 | 8.12308e-06 | chr12_56360038_C_A_b38 | -88 | 70 | 75 | 0.0787815 | 1 | 2.28E-06 | -0.20343 | 0.0424222 | 2.80E-03 | 3.01E-03 | 2.24E-03 | 2.38E-04 | Artery_Tibial | STAT2 | 12_56753822 |
| ENSG00000127337.6 | 6766 | 1.04 | 599.765 | 100.405 | 1.35837e-10 | chr12_69275233_G_A_b38 | -84470 | 21 | 22 | 0.0718954 | 1 | 9.55E-12 | 0.882641 | 0.116193 | 1.00E-04 | 4.19E-08 | 3.12E-07 | 5.65E-05 | Brain_Putamen_basal_ganglia | YEATS4 | 12_69669013 |
| ENSG00000127337.6 | 6766 | 1.05 | 554.906 | 66.7093 | 3.7855e-06 | chr12_69275233_G_A_b38 | -84470 | 11 | 11 | 0.055 | 1 | 9.41E-07 | 1.13955 | 0.21322 | 1.10E-03 | 1.51E-03 | 1.14E-02 | 2.25E-05 | Brain_Substantia_nigra | YEATS4 | 12_69669013 |
| ENSG00000122034.14 | 7500 | 1.03 | 892.961 | 247.424 | 2.16631e-05 | chr13_27385844_G_A_b38 | -38700 | 97 | 109 | 0.165653 | 1 | 7.83E-06 | -0.163247 | 0.0358254 | 1.63E-02 | 1.65E-02 | 1.64E-02 | 7.80E-05 | Artery_Aorta | GTF3A | 13_27959981 |
| ENSG00000122034.14 | 7500 | 1.03 | 927.663 | 369.617 | 1.879e-05 | chr13_27391627_G_T_b38 | -32917 | 110 | 120 | 0.12605 | 1 | 7.30E-06 | -0.137906 | 0.030351 | 1.43E-02 | 1.52E-02 | 9.96E-03 | 1.16E-04 | Artery_Tibial | GTF3A | 13_27965764 |
| ENSG00000122034.14 | 7500 | 1.03 | 881.35 | 330.945 | 1.57435e-09 | chr13_27424509_C_A_b38 | -35 | 106 | 120 | 0.137615 | 1 | 2.17E-10 | -0.212246 | 0.0324923 | 1.00E-04 | 9.54E-07 | 1.62E-06 | 9.02E-05 | Lung | GTF3A | 13_27998646 |
| ENSG00000185650.9 | 5182 | 1.06 | 433.236 | 268.939 | 7.80518e-05 | chr14_68442854_T_C_b38 | -353399 | 171 | 205 | 0.266234 | 1 | 1.90E-05 | -0.179939 | 0.0414402 | 2.33E-02 | 2.60E-02 | 1.90E-02 | 2.31E-04 | Esophagus_Muscularis | ZFP36L1 | 14_68909571 |
| ENSG00000185650.9 | 5182 | 1.06 | 453.857 | 285.727 | 7.41031e-05 | chr14_68589026_C_T_b38 | -207227 | 174 | 199 | 0.246898 | 1 | 1.88E-05 | 0.196991 | 0.0453757 | 2.30E-02 | 2.61E-02 | 1.32E-02 | 3.30E-04 | Cells_Cultured_fibroblasts | ZFP36L1 | 14_69055743 |
| ENSG00000119725.18 | 5811 | 1.06 | 356.272 | 371.495 | 1.80972e-05 | chr14_73737983_G_C_b38 | -148845 | 276 | 335 | 0.329724 | 1 | 3.23E-06 | 0.0995043 | 0.0210969 | 2.50E-03 | 4.68E-03 | 3.24E-03 | 3.44E-04 | Skin_Sun_Exposed_Lower_leg | ZNF410 | 14_74204686 |
| ENSG00000119715.14 | 6449 | 1.04 | 825.727 | 459.864 | 1.0464e-58 | chr14_76314472_C_T_b38 | 3858 | 159 | 173 | 0.147109 | 1 | 6.64E-66 | -0.624824 | 0.0313891 | 1.00E-04 | 8.62E-58 | 1.13E-56 | 1.31E-04 | Muscle_Skeletal | ESRRB | 14_76780815 |
| ENSG00000166716.9 | 5271 | 1.05 | 305.436 | 230.893 | 2.31393e-05 | chr15_84660473_G_C_b38 | -88162 | 153 | 172 | 0.261398 | 1 | 4.00E-06 | -0.197331 | 0.041927 | 4.40E-03 | 5.47E-03 | 6.09E-03 | 2.37E-04 | Artery_Aorta | ZNF592 | 15_85203704 |
| ENSG00000122386.10 | 5174 | 1.04 | 812.683 | 82.561 | 1.00163e-05 | chr16_3480658_C_G_b38 | 368098 | 30 | 33 | 0.144737 | 1 | 4.50E-06 | -0.495405 | 0.101376 | 6.70E-03 | 6.57E-03 | 2.94E-02 | 1.98E-05 | Minor_Salivary_Gland | ZNF205 | 16_3530658 |
| ENSG00000198429.9 | 5587 | 1.07 | 521.93 | 454.356 | 3.67202e-05 | chr19_11841496_A_T_b38 | -46288 | 39 | 42 | 0.0357143 | 1 | 1.05E-05 | -0.476481 | 0.107053 | 1.18E-02 | 1.42E-02 | 9.66E-03 | 2.24E-04 | Muscle_Skeletal | ZNF69 | 19_11952311 |
| ENSG00000198429.9 | 5587 | 1.03 | 521.456 | 431.709 | 0.00012325 | chr19_11887941_C_T_b38 | 157 | 14 | 14 | 0.0125448 | 1 | 4.46E-05 | -0.815907 | 0.198048 | 5.41E-02 | 5.62E-02 | 3.27E-02 | 2.04E-04 | Whole_Blood | ZNF69 | 19_11998756 |
| ENSG00000060566.13 | 7138 | 1.01 | 1180.13 | 330.048 | 2.88683e-05 | chr19_3793151_G_A_b38 | -360450 | 37 | 39 | 0.0447248 | 1 | 1.06E-05 | 0.639422 | 0.143157 | 3.08E-02 | 3.17E-02 | 2.55E-02 | 6.49E-05 | Lung | CREB3L3 | 19_3793149 |
| ENSG00000104804.7 | 6707 | 1.04 | 1063.86 | 241.421 | 2.72736e-05 | chr19_48858184_T_G_b38 | -40549 | 104 | 114 | 0.18038 | 1 | 1.29E-05 | 0.406089 | 0.0913102 | 2.27E-02 | 2.44E-02 | 2.57E-02 | 5.98E-05 | Heart_Atrial_Appendage | TULP2 | 19_49361441 |
| ENSG00000104804.7 | 6707 | 1.03 | 912.427 | 246.914 | 1.21409e-07 | chr19_48879712_G_A_b38 | -19021 | 107 | 123 | 0.18693 | 1 | 2.48E-08 | 0.442633 | 0.0770877 | 1.00E-04 | 8.09E-05 | 1.22E-04 | 7.61E-05 | Artery_Aorta | TULP2 | 19_49382969 |
| ENSG00000104804.7 | 6707 | 1.02 | 1000.69 | 129.009 | 3.6662e-24 | chr19_48888513_G_A_b38 | -10220 | 58 | 60 | 0.168539 | 1 | 9.73E-26 | 1.50253 | 0.115547 | 1.00E-04 | 1.25E-21 | 5.70E-20 | 2.34E-05 | Liver | TULP2 | 19_49391770 |
| ENSG00000104804.7 | 6707 | 1.04 | 933.342 | 329.808 | 1.16255e-21 | chr19_48888513_G_A_b38 | -10220 | 133 | 143 | 0.163991 | 1 | 7.27E-24 | 0.717426 | 0.0662965 | 1.00E-04 | 2.50E-19 | 1.19E-18 | 8.73E-05 | Lung | TULP2 | 19_49391770 |
| ENSG00000161551.14 | 9052 | 1.05 | 1277.62 | 163.383 | 1.77929e-07 | chr19_51883933_G_C_b38 | -4043 | 3 | 3 | 0.00684932 | 1 | 4.61E-08 | -1.72481 | 0.302054 | 2.00E-04 | 1.51E-04 | 3.92E-04 | 3.45E-05 | Pituitary | ZNF577 | 19_52387186 |
| ENSG00000180228.12 | 6135 | 1.05 | 374.536 | 187.011 | 7.52486e-05 | chr2_178497181_G_C_b38 | 45746 | 11 | 11 | 0.0202206 | 1 | 1.75E-05 | 0.389569 | 0.0887056 | 1.99E-02 | 2.26E-02 | 2.83E-02 | 1.47E-04 | Esophagus_Gastroesophageal_Junction | PRKRA | 2_179361908 |
| ENSG00000124831.18 | 7667 | 1.03 | 884.791 | 127.61 | 2.58155e-05 | chr2_237739878_C_T_b38 | 112302 | 18 | 18 | 0.0502793 | 1 | 1.12E-05 | 0.634134 | 0.139139 | 1.83E-02 | 2.00E-02 | 2.65E-02 | 5.62E-05 | Spleen | LRRFIP1 | 2_238648521 |
| ENSG00000115355.15 | 7119 | 1.04 | 572.317 | 104.353 | 3.43095e-10 | chr2_55454381_G_A_b38 | 34460 | 8 | 8 | 0.0254777 | 1 | 2.95E-11 | 1.34232 | 0.18268 | 1.00E-04 | 1.08E-07 | 4.06E-07 | 9.16E-05 | Brain_Cerebellar_Hemisphere | CCDC88A | 2_55681517 |
| ENSG00000115355.15 | 7119 | 1.04 | 501.699 | 127.728 | 6.66989e-17 | chr2_55454381_G_A_b38 | 34460 | 14 | 14 | 0.037234 | 1 | 2.39E-19 | 1.17366 | 0.112895 | 1.00E-04 | 8.19E-15 | 5.12E-14 | 1.27E-04 | Brain_Cerebellum | CCDC88A | 2_55681517 |
| ENSG00000115956.9 | 6927 | 1.03 | 707.587 | 105.988 | 3.24397e-06 | chr2_68248463_C_A_b38 | -116710 | 95 | 109 | 0.347134 | -1 | 1.00E-06 | 0.458411 | 0.0887581 | 2.00E-03 | 1.88E-03 | 3.35E-03 | 7.23E-05 | Brain_Cerebellar_Hemisphere | PLEK | 2_68475595 |
| ENSG00000168874.12 | 6082 | 1.02 | 555.104 | 368.952 | 5.59543e-05 | chr2_86396401_G_C_b38 | 645057 | 44 | 46 | 0.0477178 | 1 | 2.06E-05 | 0.276545 | 0.0641923 | 2.67E-02 | 2.80E-02 | 1.54E-02 | 2.19E-04 | Thyroid | ATOH8 | 2_86623524 |
| ENSG00000185513.14 | 6107 | 1.04 | 844.754 | 370.533 | 4.83968e-07 | chr20_43512293_T_C_b38 | 4613 | 27 | 28 | 0.0290456 | 1 | 1.11E-07 | 0.305203 | 0.0564937 | 3.00E-04 | 3.06E-04 | 2.27E-04 | 1.49E-04 | Thyroid | L3MBTL1 | 20_42140933 |
| ENSG00000177732.8 | 5086 | 1.05 | 892.735 | 302.793 | 1.21761e-05 | chr20_776848_A_G_b38 | 451253 | 140 | 158 | 0.192214 | 1 | 3.44E-06 | 0.183175 | 0.0388041 | 6.80E-03 | 8.26E-03 | 6.51E-03 | 1.15E-04 | Esophagus_Mucosa | SOX12 | 20_757491 |
| ENSG00000141956.13 | 8869 | 1.03 | 1410.06 | 371.034 | 1.60297e-28 | chr21_41800951_G_A_b38 | -78531 | 80 | 83 | 0.0866388 | 1 | 2.84E-31 | 0.913051 | 0.0720513 | 1.00E-04 | 3.21E-26 | 1.45E-25 | 7.40E-05 | Adipose_Subcutaneous | PRDM15 | 21_43221111 |

**Table S2: GTEx cis-eQTLs (FDR<0.05) between aSNPs and TFs**

| phenotype_id | num_var | beta_shape1 | beta_shape2 | true_df | pval_true_df | variant_id | tss_distance | ma_samples | ma_count | maf | ref_factor | pval_nominal | slope | slope_se | pval_perm | pval_beta | qval | pval_nominal_threshold | tissue | gene | hg19 |
|---|---|---|---|---|---|---|---|---|---|---|---|---|---|---|---|---|---|---|---|---|---|
| ENSG00000141956.13 | 8869 | 1.02 | 1496.74 | 194.106 | 1.61106e-11 | chr21_41800951_G_A_b38 | -78531 | 56 | 59 | 0.110902 | 1 | 2.14E-12 | 0.632665 | 0.0847325 | 1.00E-04 | 1.62E-08 | 5.97E-08 | 3.16E-05 | Colon_Sigmoid | PRDM15 | 21_43221111 |
| ENSG00000141956.13 | 8869 | 1.06 | 1378.27 | 307.348 | 7.25607e-20 | chr21_41800951_G_A_b38 | -78531 | 73 | 74 | 0.0900243 | 1 | 7.18E-22 | 0.673959 | 0.0653783 | 1.00E-04 | 1.27E-17 | 4.22E-17 | 7.47E-05 | Esophagus_Mucosa | PRDM15 | 21_43221111 |
| ENSG00000141956.13 | 8869 | 1.01 | 1493.81 | 335.968 | 5.14614e-18 | chr21_41800951_G_A_b38 | -78531 | 77 | 78 | 0.0894495 | 1 | 1.73E-19 | 0.576428 | 0.0602768 | 1.00E-04 | 5.42E-15 | 1.94E-14 | 5.06E-05 | Lung | PRDM15 | 21_43221111 |
| ENSG00000141956.13 | 8869 | 1.02 | 1462.01 | 82.9803 | 2.43504e-08 | chr21_41800951_G_A_b38 | -78531 | 31 | 32 | 0.140351 | 1 | 7.57E-09 | 0.844431 | 0.132248 | 1.00E-04 | 2.99E-05 | 2.83E-04 | 9.87E-06 | Minor_Salivary_Gland | PRDM15 | 21_43221111 |
| ENSG00000141956.13 | 8869 | 1.04 | 1405.74 | 325.711 | 4.15154e-26 | chr21_41800951_G_A_b38 | -78531 | 72 | 75 | 0.0872093 | 1 | 1.10E-28 | 0.734031 | 0.060432 | 1.00E-04 | 7.93E-24 | 3.88E-23 | 6.62E-05 | Skin_Not_Sun_Exposed_Suprapubic | PRDM15 | 21_43221111 |
| ENSG00000141956.13 | 8869 | 1.02 | 1505.61 | 378.63 | 1.43584e-30 | chr21_41800951_G_A_b38 | -78531 | 84 | 86 | 0.0892116 | 1 | 4.05E-33 | 0.80693 | 0.0614849 | 1.00E-04 | 7.56E-28 | 2.60E-27 | 7.98E-05 | Thyroid | PRDM15 | 21_43221111 |
| ENSG00000141956.13 | 8869 | 1.04 | 1368.19 | 216.36 | 1.62181e-10 | chr21_41809755_T_C_b38 | -69727 | 54 | 54 | 0.0918367 | 1 | 1.81E-11 | 0.63186 | 0.0895152 | 1.00E-04 | 1.26E-07 | 3.89E-07 | 4.05E-05 | Colon_Transverse | PRDM15 | 21_43230111 |
| ENSG00000141956.13 | 8869 | 1.03 | 1508.94 | 108.49 | 1.72325e-08 | chr21_41809755_T_C_b38 | -69727 | 22 | 24 | 0.0851064 | 1 | 5.53E-09 | 0.586819 | 0.0931444 | 1.00E-04 | 1.94E-05 | 1.34E-04 | 1.47E-05 | Small_Intestine_Terminal_Ileum | PRDM15 | 21_43230111 |
| ENSG00000141956.13 | 8869 | 1.05 | 1375.75 | 184.889 | 2.58004e-06 | chr21_41809755_T_C_b38 | -69727 | 38 | 38 | 0.0730769 | 1 | 7.37E-07 | 0.628485 | 0.122996 | 1.90E-03 | 2.55E-03 | 5.71E-03 | 2.94E-05 | Stomach | PRDM15 | 21_43230111 |
| ENSG00000275004.3 | 8424 | 1.03 | 943.242 | 467.475 | 6.68656e-05 | chr22_22468169_G_A_b38 | -40985 | 44 | 44 | 0.037415 | 1 | 2.70E-05 | 0.438463 | 0.103525 | 5.22E-02 | 5.49E-02 | 3.20E-02 | 1.14E-04 | Muscle_Skeletal | ZNF280B | 22_22822506 |
| ENSG00000181722.16 | 4852 | 1.03 | 584.758 | 296.594 | 1.52544e-13 | chr3_114336583_T_C_b38 | -810688 | 17 | 17 | 0.0210918 | 1 | 5.05E-15 | 0.76808 | 0.0935954 | 1.00E-04 | 4.53E-11 | 6.07E-11 | 2.36E-04 | Cells_Cultured_fibroblasts | ZBTB20 | 3_114055430 |
| ENSG00000114126.17 | 5585 | 1.04 | 544.934 | 86.2 | 2.86545e-05 | chr3_142172543_C_A_b38 | 23006 | 22 | 24 | 0.10084 | 1 | 1.24E-05 | 0.509703 | 0.110442 | 1.13E-02 | 1.30E-02 | 4.71E-02 | 3.12E-05 | Brain_Amygdala | TFDP2 | 3_141891385 |
| ENSG00000114166.7 | 6461 | 1.02 | 546.649 | 316.571 | 0.000151811 | chr3_20010528_C_T_b38 | -29495 | 212 | 257 | 0.298837 | 1 | 5.35E-05 | -0.137457 | 0.0336156 | 7.32E-02 | 7.55E-02 | 4.78E-02 | 1.60E-04 | Skin_Not_Sun_Exposed_Suprapubic | KAT2B | 3_20052020 |
| ENSG00000114166.7 | 6461 | 1.03 | 508.937 | 227.116 | 1.08077e-09 | chr3_20063346_C_G_b38 | 23323 | 144 | 159 | 0.251582 | 1 | 6.28E-11 | 0.225598 | 0.0330749 | 1.00E-04 | 3.36E-07 | 8.37E-07 | 1.23E-04 | Heart_Atrial_Appendage | KAT2B | 3_20104838 |
| ENSG00000174738.12 | 7073 | 1.06 | 639.314 | 312.895 | 0.000148026 | chr3_23066257_C_T_b38 | -879003 | 53 | 54 | 0.0627907 | 1 | 4.69E-05 | 0.333582 | 0.080955 | 7.66E-02 | 7.71E-02 | 4.86E-02 | 1.53E-04 | Skin_Not_Sun_Exposed_Suprapubic | NR1D2 | 3_23107748 |
| ENSG00000151090.17 | 7337 | 1.06 | 800.997 | 453.703 | 2.18714e-06 | chr3_24458364_G_C_b38 | -36918 | 29 | 29 | 0.0246599 | 1 | 4.19E-07 | -0.509456 | 0.0993908 | 1.10E-03 | 1.20E-03 | 9.88E-04 | 1.42E-04 | Muscle_Skeletal | THRB | 3_24499855 |
| ENSG00000138738.10 | 7670 | 1.04 | 546.358 | 104.684 | 1.06299e-06 | chr4_120722571_C_T_b38 | -200299 | 57 | 64 | 0.203822 | 1 | 2.47E-07 | 0.396235 | 0.0722963 | 4.00E-04 | 4.25E-04 | 1.41E-03 | 6.14E-05 | Brain_Frontal_Cortex_BA9 | PRDM5 | 4_121643726 |
| ENSG00000138738.10 | 7670 | 1.04 | 526.889 | 122.838 | 1.08186e-12 | chr4_120782415_T_C_b38 | -140455 | 86 | 98 | 0.273743 | 1 | 3.58E-14 | 0.569275 | 0.0673967 | 1.00E-04 | 2.22E-10 | 1.23E-09 | 9.91E-05 | Spleen | PRDM5 | 4_121703570 |
| ENSG00000173545.4 | 6281 | 1.01 | 839.141 | 247.485 | 2.28931e-05 | chr5_16496741_C_T_b38 | 30949 | 26 | 26 | 0.0397554 | 1 | 9.00E-06 | -0.233452 | 0.0515828 | 1.56E-02 | 1.79E-02 | 2.18E-02 | 6.21E-05 | Heart_Left_Ventricle | ZNF622 | 5_16496850 |
| ENSG00000112365.4 | 5788 | 1.03 | 509.026 | 327.646 | 8.17732e-05 | chr6_108828668_G_A_b38 | -654569 | 22 | 22 | 0.0252294 | 1 | 3.13E-05 | 0.353212 | 0.0837714 | 3.69E-02 | 3.71E-02 | 2.92E-02 | 1.56E-04 | Lung | ZBTB24 | 6_109149871 |
| ENSG00000137273.3 | 7478 | 1.03 | 1205.32 | 374.503 | 1.01897e-05 | chr6_1364236_A_C_b38 | -25598 | 107 | 115 | 0.120042 | 1 | 3.99E-06 | 0.3091 | 0.066108 | 1.14E-02 | 1.07E-02 | 7.68E-03 | 8.50E-05 | Adipose_Subcutaneous | FOXF2 | 6_1364471 |
| ENSG00000137273.3 | 7478 | 1.05 | 1111.61 | 330.332 | 8.1625e-05 | chr6_1367348_G_A_b38 | -22486 | 84 | 89 | 0.101598 | 1 | 3.20E-05 | 0.302924 | 0.0719414 | 7.17E-02 | 7.53E-02 | 3.61E-02 | 1.20E-04 | Nerve_Tibial | FOXF2 | 6_1367583 |
| ENSG00000054598.6 | 7491 | 1.05 | 1157.05 | 186.129 | 4.46967e-05 | chr6_1623392_G_C_b38 | 13420 | 73 | 79 | 0.162551 | 1 | 2.02E-05 | 0.420223 | 0.0962488 | 4.13E-02 | 4.24E-02 | 4.94E-02 | 4.54E-05 | Pancreas | FOXC1 | 6_1623627 |
| ENSG00000172201.11 | 7223 | 1.05 | 776.957 | 465.656 | 1.38123e-05 | chr6_18921711_T_A_b38 | -915675 | 91 | 94 | 0.079932 | 1 | 4.54E-06 | -0.283548 | 0.0611877 | 7.70E-03 | 8.46E-03 | 6.03E-03 | 1.43E-04 | Muscle_Skeletal | ID4 | 6_18921942 |
| ENSG00000187626.8 | 6761 | 1.03 | 136.837 | 117.128 | 4.03862e-05 | chr6_27393463_T_C_b38 | -858761 | 27 | 28 | 0.0765027 | 1 | 5.68E-06 | -0.710562 | 0.150706 | 3.90E-03 | 4.74E-03 | 8.76E-03 | 3.12E-04 | Brain_Cortex | ZKSCAN4 | 6_27361242 |
| ENSG00000112561.17 | 6196 | 1.03 | 849.276 | 301.952 | 0.000140338 | chr6_41705329_T_C_b38 | -30930 | 92 | 107 | 0.132754 | 1 | 6.37E-05 | -0.218867 | 0.0540354 | 1.00E-01 | 1.03E-01 | 4.39E-02 | 1.65E-04 | Cells_Cultured_fibroblasts | TFEB | 6_41673067 |
| ENSG00000112561.17 | 6196 | 1.02 | 911.443 | 375.71 | 2.64276e-06 | chr6_41733120_T_C_b38 | -3139 | 61 | 66 | 0.0688935 | 1 | 9.55E-07 | -0.288933 | 0.0580575 | 2.40E-03 | 2.14E-03 | 1.74E-03 | 1.10E-04 | Adipose_Subcutaneous | TFEB | 6_41700858 |
| ENSG00000112561.17 | 6196 | 1.04 | 856.563 | 331.809 | 3.49157e-05 | chr6_41733120_T_C_b38 | -3139 | 49 | 53 | 0.0607798 | 1 | 1.38E-05 | -0.282543 | 0.064114 | 2.62E-02 | 2.48E-02 | 2.06E-02 | 9.76E-05 | Lung | TFEB | 6_41700858 |
| ENSG00000171467.15 | 5401 | 1.03 | 728.959 | 304.843 | 5.60647e-09 | chr6_43403177_C_T_b38 | 33699 | 86 | 90 | 0.109489 | 1 | 7.07E-10 | 0.284432 | 0.0448261 | 1.00E-04 | 2.66E-06 | 3.45E-06 | 1.33E-04 | Esophagus_Mucosa | ZNF318 | 6_43370915 |
| ENSG00000198517.9 | 8431 | 1.04 | 781.036 | 377.933 | 1.7786e-08 | chr7_1556748_G_A_b38 | 26034 | 65 | 65 | 0.0639764 | 1 | 1.33E-09 | -0.275494 | 0.0444554 | 1.00E-04 | 8.86E-06 | 8.72E-06 | 1.49E-04 | Skin_Sun_Exposed_Lower_leg | MAFK | 7_1596384 |
| ENSG00000106031.7 | 5864 | 1.02 | 533.848 | 459.407 | 0.000126495 | chr7_27895635_A_C_b38 | 695529 | 83 | 88 | 0.0748299 | -1 | 4.69E-05 | 0.322241 | 0.0784936 | 6.07E-02 | 6.16E-02 | 3.54E-02 | 1.92E-04 | Muscle_Skeletal | HOXA13 | 7_27935254 |
| ENSG00000106571.12 | 6151 | 1.04 | 616.979 | 320.217 | 1.13259e-05 | chr7_43000485_A_G_b38 | 763426 | 16 | 16 | 0.0186047 | 1 | 3.24E-06 | -0.623612 | 0.131853 | 5.80E-03 | 5.50E-03 | 4.70E-03 | 1.53E-04 | Skin_Not_Sun_Exposed_Suprapubic | GLI3 | 7_43040084 |
| ENSG00000122515.14 | 5973 | 1.04 | 674.589 | 128.092 | 1.10156e-07 | chr7_44751322_T_C_b38 | 2741 | 55 | 58 | 0.162011 | 1 | 3.18E-08 | -0.365983 | 0.0624361 | 1.00E-04 | 5.22E-05 | 1.31E-04 | 7.54E-05 | Spleen | ZMIZ2 | 7_44790921 |
| ENSG00000122515.14 | 5973 | 1.03 | 606.622 | 364.235 | 1.09951e-06 | chr7_44759748_T_C_b38 | 11167 | 127 | 137 | 0.143006 | 1 | 2.42E-07 | -0.20528 | 0.0390807 | 4.00E-04 | 5.46E-04 | 4.84E-04 | 1.68E-04 | Adipose_Subcutaneous | ZMIZ2 | 7_44799347 |
| ENSG00000122515.14 | 5973 | 1.02 | 561.98 | 243.07 | 1.44988e-08 | chr7_44759748_T_C_b38 | 11167 | 88 | 97 | 0.147416 | 1 | 1.77E-09 | -0.215123 | 0.0345397 | 1.00E-04 | 6.33E-06 | 1.70E-05 | 8.51E-05 | Breast_Mammary_Tissue | ZMIZ2 | 7_44799347 |
| ENSG00000122515.14 | 5973 | 1.04 | 586.873 | 327.4 | 7.99597e-17 | chr7_44759748_T_C_b38 | 11167 | 112 | 122 | 0.139269 | 1 | 1.01E-18 | -0.308515 | 0.0330669 | 1.00E-04 | 1.36E-14 | 2.43E-14 | 2.21E-04 | Nerve_Tibial | ZMIZ2 | 7_44799347 |
| ENSG00000122515.14 | 5973 | 1.05 | 630.425 | 183.706 | 1.11739e-10 | chr7_44759748_T_C_b38 | 11167 | 55 | 61 | 0.125514 | 1 | 1.19E-11 | -0.360939 | 0.0501738 | 1.00E-04 | 3.16E-08 | 1.12E-07 | 8.24E-05 | Pancreas | ZMIZ2 | 7_44799347 |
| ENSG00000122515.14 | 5973 | 1.02 | 587.599 | 320.537 | 2.56603e-14 | chr7_44759748_T_C_b38 | 11167 | 108 | 118 | 0.137209 | 1 | 6.77E-16 | -0.265375 | 0.0313658 | 1.00E-04 | 8.31E-12 | 1.85E-11 | 1.52E-04 | Skin_Not_Sun_Exposed_Suprapubic | ZMIZ2 | 7_44799347 |
| ENSG00000122515.14 | 5973 | 1.05 | 578.804 | 387.186 | 9.18828e-15 | chr7_44759748_T_C_b38 | 11167 | 140 | 153 | 0.150591 | 1 | 1.67E-16 | -0.24213 | 0.0282234 | 1.00E-04 | 1.50E-12 | 2.65E-12 | 2.07E-04 | Skin_Sun_Exposed_Lower_leg | ZMIZ2 | 7_44799347 |
| ENSG00000122515.14 | 5973 | 1.02 | 602.307 | 435.225 | 4.66113e-16 | chr7_44759748_T_C_b38 | 11167 | 146 | 162 | 0.145161 | 1 | 8.06E-18 | -0.150245 | 0.0168062 | 1.00E-04 | 1.49E-13 | 2.91E-13 | 1.71E-04 | Whole_Blood | ZMIZ2 | 7_44799347 |
| ENSG00000122515.14 | 5973 | 1.04 | 580.11 | 285.474 | 7.61524e-08 | chr7_44762083_T_C_b38 | 13502 | 104 | 112 | 0.142494 | 1 | 1.07E-08 | -0.214475 | 0.0365272 | 2.00E-04 | 2.87E-05 | 4.81E-05 | 1.21E-04 | Adipose_Visceral_Omentum | ZMIZ2 | 7_44801682 |
| ENSG00000122515.14 | 5973 | 1.04 | 614.942 | 184.617 | 1.5559e-05 | chr7_44764345_G_T_b38 | 15764 | 73 | 76 | 0.146154 | 1 | 5.28E-06 | -0.221052 | 0.0472631 | 6.70E-03 | 7.82E-03 | 1.53E-02 | 6.22E-05 | Stomach | ZMIZ2 | 7_44803944 |
| ENSG00000122515.14 | 5973 | 1.02 | 617.604 | 370.213 | 4.63201e-23 | chr7_44764345_G_T_b38 | 15764 | 125 | 138 | 0.143154 | 1 | 1.75E-25 | -0.283097 | 0.0253459 | 1.00E-04 | 1.08E-20 | 2.63E-20 | 1.97E-04 | Thyroid | ZMIZ2 | 7_44803944 |
| ENSG00000178665.14 | 7709 | 1.05 | 469.144 | 421.222 | 8.13487e-09 | chr7_55864680_T_C_b38 | -22795 | 49 | 50 | 0.0448029 | 1 | 5.42E-10 | 0.76875 | 0.121365 | 1.00E-04 | 2.09E-06 | 2.33E-06 | 2.36E-04 | Whole_Blood | ZNF713 | 7_55932373 |
| ENSG00000198039.11 | 6666 | 1.05 | 223.332 | 119.642 | 1.79373e-06 | chr7_64861244_C_T_b38 | -8928 | 22 | 22 | 0.0607735 | 1 | 2.15E-07 | 0.394419 | 0.0723233 | 4.00E-04 | 2.60E-04 | 7.73E-04 | 1.83E-04 | Brain_Nucleus_accumbens_basal_ganglia | ZNF273 | 7_64321622 |
| ENSG00000197008.9 | 6773 | 1.05 | 253.18 | 173.089 | 4.56061e-08 | chr7_64925965_A_C_b38 | 131577 | 24 | 24 | 0.0512821 | 1 | 3.17E-09 | -0.778455 | 0.125639 | 1.00E-04 | 6.55E-06 | 1.74E-05 | 2.06E-04 | Pancreas | ZNF138 | 7_64386343 |
| ENSG00000197008.9 | 6773 | 1.05 | 245.828 | 271.188 | 6.20277e-10 | chr7_64948473_C_A_b38 | 154085 | 40 | 40 | 0.0508906 | 1 | 1.47E-11 | -0.535213 | 0.076434 | 1.00E-04 | 7.08E-08 | 1.64E-07 | 2.90E-04 | Adipose_Visceral_Omentum | ZNF138 | 7_64408851 |
| ENSG00000152926.14 | 6632 | 1.06 | 209.787 | 182.285 | 8.84469e-09 | chr7_65312360_G_C_b38 | 321552 | 27 | 27 | 0.0496324 | 1 | 3.15E-10 | -0.47115 | 0.0714301 | 1.00E-04 | 7.87E-07 | 1.42E-06 | 3.96E-04 | Testis | ZNF117 | 7_64777273 |
| ENSG00000152926.14 | 6632 | 1.05 | 211.705 | 150.62 | 2.95869e-10 | chr7_65375489_C_A_b38 | 384681 | 46 | 48 | 0.109589 | 1 | 6.39E-12 | -0.54106 | 0.0735034 | 1.00E-04 | 2.55E-08 | 1.16E-07 | 2.13E-04 | Pituitary | ZNF117 | 7_64840402 |
| ENSG00000236609.3 | 9079 | 1.06 | 750.362 | 444.207 | 7.52723e-06 | chr7_6499492_A_G_b38 | -116125 | 43 | 43 | 0.0365646 | 1 | 1.32E-06 | -0.300177 | 0.0613383 | 3.10E-03 | 3.92E-03 | 2.98E-03 | 1.55E-04 | Muscle_Skeletal | ZNF853 | 7_6539123 |
| ENSG00000136870.10 | 7467 | 1.03 | 609.122 | 364.089 | 4.66335e-15 | chr9_101403009_C_A_b38 | 4136 | 67 | 70 | 0.0730689 | 1 | 9.94E-17 | 0.463077 | 0.0533867 | 1.00E-04 | 1.24E-12 | 2.54E-12 | 1.70E-04 | Adipose_Subcutaneous | ZNF189 | 9_104165291 |
| ENSG00000136870.10 | 7467 | 1.04 | 570.594 | 283.199 | 5.01757e-09 | chr9_101403009_C_A_b38 | 4136 | 59 | 62 | 0.0788804 | 1 | 4.25E-10 | 0.47422 | 0.0736007 | 1.00E-04 | 1.61E-06 | 3.18E-06 | 1.24E-04 | Adipose_Visceral_Omentum | ZNF189 | 9_104165291 |
| ENSG00000136870.10 | 7467 | 1.05 | 564.822 | 193.548 | 3.2685e-07 | chr9_101403009_C_A_b38 | 4136 | 40 | 42 | 0.0763636 | 1 | 5.15E-08 | 0.464117 | 0.0822682 | 2.00E-04 | 1.20E-04 | 2.47E-04 | 9.62E-05 | Esophagus_Gastroesophageal_Junction | ZNF189 | 9_104165291 |
| ENSG00000136870.10 | 7467 | 1.04 | 543.315 | 274.24 | 1.53021e-18 | chr9_101403009_C_A_b38 | 4136 | 57 | 60 | 0.0779221 | 1 | 4.56E-21 | 0.602882 | 0.0595044 | 1.00E-04 | 1.85E-16 | 5.98E-16 | 1.74E-04 | Esophagus_Muscularis | ZNF189 | 9_104165291 |
| ENSG00000136870.10 | 7467 | 1.05 | 593.382 | 322.53 | 2.87985e-23 | chr9_101403009_C_A_b38 | 4136 | 67 | 73 | 0.0837156 | 1 | 3.45E-26 | 0.538833 | 0.0470204 | 1.00E-04 | 1.86E-21 | 1.01E-20 | 1.43E-04 | Lung | ZNF189 | 9_104165291 |
| ENSG00000136870.10 | 7467 | 1.05 | 543.384 | 312.119 | 2.71201e-10 | chr9_101403009_C_A_b38 | 4136 | 67 | 71 | 0.0825581 | 1 | 1.18E-11 | 0.329454 | 0.0469989 | 1.00E-04 | 6.87E-08 | 1.08E-07 | 1.75E-04 | Skin_Not_Sun_Exposed_Suprapubic | ZNF189 | 9_104165291 |
| ENSG00000136870.10 | 7467 | 1.04 | 560.941 | 381.167 | 2.32947e-14 | chr9_101403009_C_A_b38 | 4136 | 77 | 81 | 0.0797244 | 1 | 2.82E-16 | 0.40108 | 0.0471378 | 1.00E-04 | 4.31E-12 | 7.37E-12 | 2.11E-04 | Skin_Sun_Exposed_Lower_leg | ZNF189 | 9_104165291 |
| ENSG00000136870.10 | 7467 | 1.04 | 583.812 | 361.237 | 3.12469e-07 | chr9_101403009_C_A_b38 | 4136 | 70 | 74 | 0.0767635 | 1 | 4.66E-08 | 0.343264 | 0.0616513 | 2.00E-04 | 1.24E-04 | 9.67E-05 | 2.20E-04 | Thyroid | ZNF189 | 9_104165291 |
| ENSG00000136870.10 | 7467 | 1.03 | 625.682 | 138.324 | 1.41883e-05 | chr9_101409574_C_T_b38 | 10701 | 26 | 27 | 0.0695876 | 1 | 4.62E-06 | 0.544493 | 0.114633 | 7.20E-03 | 7.42E-03 | 1.55E-02 | 5.67E-05 | Adrenal_Gland | ZNF189 | 9_104171856 |
| ENSG00000136870.10 | 7467 | 1.05 | 574.72 | 240.433 | 5.48284e-06 | chr9_101409574_C_T_b38 | 10701 | 48 | 50 | 0.0759878 | 1 | 1.22E-06 | 0.310813 | 0.0626158 | 2.30E-03 | 2.30E-03 | 2.76E-03 | 1.28E-04 | Artery_Aorta | ZNF189 | 9_104171856 |
| ENSG00000136870.10 | 7467 | 1.04 | 635.081 | 361.107 | 1.10437e-15 | chr9_101409574_C_T_b38 | 10701 | 69 | 73 | 0.0766807 | 1 | 1.91E-17 | 0.437754 | 0.0492013 | 1.00E-04 | 2.18E-13 | 4.32E-13 | 1.75E-04 | Artery_Tibial | ZNF189 | 9_104171856 |
| ENSG00000136870.10 | 7467 | 1.07 | 548.965 | 237.618 | 2.19925e-07 | chr9_101409574_C_T_b38 | 10701 | 56 | 59 | 0.0896657 | 1 | 2.62E-08 | 0.394818 | 0.0688858 | 2.00E-04 | 6.52E-05 | 1.49E-04 | 1.01E-04 | Breast_Mammary_Tissue | ZNF189 | 9_104171856 |
| ENSG00000136870.10 | 7467 | 1.02 | 600.586 | 296.125 | 3.26032e-06 | chr9_101409574_C_T_b38 | 10701 | 54 | 57 | 0.0707196 | 1 | 8.00E-07 | 0.279975 | 0.0556505 | 1.40E-03 | 1.67E-03 | 1.05E-03 | 2.27E-04 | Cells_Cultured_fibroblasts | ZNF189 | 9_104171856 |
| ENSG00000136870.10 | 7467 | 1.05 | 605.593 | 212.488 | 7.69066e-07 | chr9_101409574_C_T_b38 | 10701 | 46 | 48 | 0.0816327 | 1 | 1.58E-07 | 0.394922 | 0.0730876 | 2.00E-04 | 3.18E-04 | 6.02E-04 | 9.49E-05 | Colon_Transverse | ZNF189 | 9_104171856 |

**Table S2: GTEx cis-eQTLs (FDR<0.05) between aSNPs and TFs**

| phenotype_id | num_var | beta_shape1 | beta_shape2 | true_df | pval_true_df | variant_id | tss_distance | ma_samples | ma_count | maf | ref_factor | pval_nominal | slope | slope_se | pval_perm | pval_beta | qval | pval_nominal_threshold | tissue | gene | hg19 |
|---|---|---|---|---|---|---|---|---|---|---|---|---|---|---|---|---|---|---|---|---|---|
| ENSG00000136870.10 | 7467 | 1.07 | 561.345 | 228.607 | 6.15072e-05 | chr9_101409574_C_T_b38 | 10701 | 52 | 55 | 0.0870253 | 1 | 1.85E-05 | 0.297815 | 0.0682651 | 2.42E-02 | 2.63E-02 | 2.73E-02 | 1.23E-04 | Heart_Atrial_Appendage | ZNF189 | 9_104171856 |
| ENSG00000136870.10 | 7467 | 1.04 | 567.645 | 426.882 | 4.9763e-07 | chr9_101409574_C_T_b38 | 10701 | 78 | 84 | 0.0752688 | 1 | 7.64E-08 | 0.209794 | 0.0384315 | 2.00E-04 | 1.98E-04 | 1.78E-04 | 1.92E-04 | Whole_Blood | ZNF189 | 9_104171856 |
| ENSG00000173258.12 | 7063 | 1.04 | 544.724 | 355.686 | 1.24308e-16 | chr9_111475730_T_C_b38 | -49429 | 64 | 64 | 0.0668058 | 1 | 6.78E-19 | 0.626996 | 0.067201 | 1.00E-04 | 1.96E-14 | 4.59E-14 | 1.95E-04 | Adipose_Subcutaneous | ZNF483 | 9_114238010 |
| ENSG00000173258.12 | 7063 | 1.03 | 623.141 | 133.429 | 4.57347e-05 | chr9_111560219_G_A_b38 | 35060 | 23 | 25 | 0.0664894 | 1 | 1.75E-05 | 0.429562 | 0.0967661 | 2.10E-02 | 2.47E-02 | 2.64E-02 | 9.84E-05 | Brain_Cerebellum | ZNF483 | 9_114322499 |
| ENSG00000167081.16 | 4839 | 1.04 | 513.072 | 127.714 | 1.54258e-05 | chr9_126173459_C_T_b38 | 426114 | 48 | 56 | 0.153005 | 1 | 4.77E-06 | 0.398109 | 0.0837021 | 4.50E-03 | 6.31E-03 | 1.12E-02 | 8.78E-05 | Brain_Cortex | PBX3 | 9_128935738 |

**Table S3: Functional enrichment for a-cTFs**

| #category | term ID | term description | observed gene count | background gene count | strength | false discovery rate | matching proteins in your network (labels) |
|---|---|---|---|---|---|---|---|
| GO Function | GO:0003700 | DNA-binding transcription factor activity | 39 | 1238 | 1.01 | 1.03E-28 | CREB3L3,HOXA13,ZBTB24,PRRX1,ZNF10,FOXF2,PRDM5,ZNF117,NHLH1,ATOH8,NR1D2, ZNF483,STAT2,PGR,LMX1A,MAFK,ZNF69,SOX12,ZNF140,TFEB,PROX1,JUN,PBX3,ATOH7, ZKSCAN4,ID4,GATA3,FOXC1,LRRFIP1,GLI3,ZNF143,THRB,ZFP36L1,ZFP69B,ZBTB20,DP2, ESRRB,ZNF205,ZNF280B |
| GO Function | GO:0000976 | Transcription regulatory region sequence-specific dna binding | 33 | 1028 | 1.02 | 1.10E-23 | CREB3L3,ZBTB24,PRRX1,FOXF2,KAT2B,RBBP5,PRDM5,PRDM15,ZNF117,NHLH1,ZNF138, ATOH8,NR1D2,STAT2,PGR,LMX1A,MAFK,SOX12,TFEB,PROX1,JUN,PBX3,GATA3,FOXC1, LRRFIP1,GLI3,ZNF143,THRB,ZNF713,ZNF273,ZBTB20,ESRRB,ZNF205 |
| GO Function | GO:0000981 | DNA-binding transcription factor activity, RNA polymerase II-specific | 32 | 1022 | 1.01 | 1.27E-22 | CREB3L3,HOXA13,ZBTB24,PRRX1,FOXF2,PRDM5,NHLH1,ATOH8,NR1D2,STAT2,PGR,LMX1A, MAFK,SOX12,ZNF140,TFEB,PROX1,JUN,PBX3,ATOH7,ID4,GATA3,FOXC1,LRRFIP1,GLI3,ZNF143, THRB,ZFP36L1,ZBTB20,DP2,ESRRB,ZNF205 |
| GO Function | GO:0000977 | RNA polymerase II transcription regulatory region sequence-specific DNA binding | 29 | 878 | 1.03 | 1.05E-20 | CREB3L3,PRRX1,FOXF2,KAT2B,PRDM5,PRDM15,ZNF117,NHLH1,ZNF138,ATOH8,NR1D2,STAT2, PGR,LMX1A,MAFK,PROX1,JUN,PBX3,GATA3,FOXC1,LRRFIP1,GLI3,ZNF143,THRB,ZNF713, ZNF273,ZBTB20,ESRRB,ZNF205 |
| GO Function | GO:0000978 | RNA polymerase II cis-regulatory region sequence-specific DNA binding | 22 | 672 | 1.03 | 7.27E-15 | CREB3L3,PRRX1,PRDM5,PRDM15,ZNF117,ZNF138,ATOH8,NR1D2,STAT2,PGR,MAFK,PROX1, JUN,PBX3,GATA3,LRRFIP1,GLI3,ZNF143,ZNF713,ZNF273,ZBTB20,ESRRB |
| GO Function | GO:0001227 | DNA-binding transcription repressor activity, RNA polymerase II-specific | 10 | 260 | 1.1 | 1.19E-06 | PRRX1,PRDM5,NR1D2,MAFK,ZNF140,PROX1,GATA3,LRRFIP1,ZBTB20,ZNF205 |
| GO Function | GO:0004879 | Nuclear receptor activity | 4 | 48 | 1.43 | 2.30E-03 | NR1D2,PGR,THRB,ESRRB |
| GO Function | GO:0071837 | HMG box domain binding | 3 | 16 | 1.79 | 2.80E-03 | PRRX1,JUN,GATA3 |
| GO Function | GO:0003707 | Steroid hormone receptor activity | 4 | 55 | 1.38 | 3.40E-03 | NR1D2,PGR,THRB,ESRRB |
| GO Process | GO:0007623 | Circadian rhythm | 5 | 142 | 1.06 | 1.52E-02 | NR1D2,PROX1,JUN,ATOH7,ID4 |
| GO Process | GO:1902107 | Positive regulation of leukocyte differentiation | 5 | 156 | 1.02 | 2.30E-02 | SOX12,JUN,GATA3,GLI3,ZFP36L1 |
| GO Process | GO:0021766 | Hippocampus development | 4 | 81 | 1.21 | 2.30E-02 | LMX1A,PROX1,ID4,GLI3 |
| GO Process | GO:0006367 | Transcription initiation from rna polymerase ii promoter | 5 | 162 | 1.0 | 2.59E-02 | KAT2B,NR1D2,PGR,THRB,ESRRB |
| GO Process | GO:0060840 | Artery development | 4 | 88 | 1.17 | 2.90E-02 | PRRX1,PROX1,FOXC1,GLI3 |
| GO Process | GO:0110111 | Negative regulation of animal organ morphogenesis | 3 | 34 | 1.46 | 3.07E-02 | GATA3,FOXC1,THRB |
| GO Process | GO:0022612 | Gland morphogenesis | 4 | 99 | 1.12 | 3.97E-02 | PGR,PROX1,ID4,GLI3 |
| GO Process | GO:1902895 | Positive regulation of pri-mirna transcription by rna polymerase ii | 3 | 40 | 1.39 | 4.34E-02 | ATOH8,JUN,GATA3 |
| GO Process | GO:0001709 | Cell fate determination | 3 | 42 | 1.37 | 4.70E-02 | PRRX1,PROX1,GATA3 |
| Reactome | HSA-383280 | Nuclear Receptor transcription pathway | 4 | 53 | 1.39 | 1.50E-02 | NR1D2,PGR,THRB,ESRRB |

**Table S4: TF target information**

| Database | Number of tested TFs | Total target genes | GTEx<br>Number of overlapping Neandertal regulated TFs | GTEx<br>Target genes for Neandertal TFs | GTEx<br>Neandertal target genes in deserts |
|---|---|---|---|---|---|
| TRANSFAC predicted targets | 158 | 17,656 | 3 | 4,572 | 56 |
| TRANSFAC curated targets | 201 | 11,024 | 10 | 3,384 | 51 |
| MotifMap predicted targets | 207 | 16,258 | 1 | 189 | 4 |
| JASPAR predicted targets | 111 | 17,630 | 6 | 12,874 | 183 |
| ENCODE | 181 | 17,854 | 8 | 17,086 | 227 |
| CHEA | 199 | 17,634 | 5 | 4,595 | 79 |
| Human TFTG DB | 671 | 17,878 | 18 | 13,667 | 189 |
| unique | 870 (of 1795) | 18,895 | 27 (of 60) | 18,415 | 244 (of 248/257) |

**Table S5: Overlap between prediction databases**

Percentages in () are calculated based on the total number in database in row (below diagonal) and column (above diagnonal)

| | CHEA | ENCODE | TRANSFAC_Cur | TRANSFAC_Pred | JASPAR_Pred | MotifMap_Pred | Human_TFTG_DB | Animal_TFDB |
|---|---|---|---|---|---|---|---|---|
| CHEA | 199 | 70 (35.18) | 72 (36.18) | 54 (27.14) | 54 (27.14) | 63 (31.66) | 136 (68.34) | 154 (77.39) |
| ENCODE | 70 (38.67) | 181 | 54 (29.83) | 48 (26.52) | 50 (27.62) | 52 (28.73) | 100 (55.25) | 122 (67.4) |
| TRANSFAC_Cur | 72 (35.82) | 54 (26.87) | 201 | 72 (35.82) | 61 (30.35) | 92 (45.77) | 180 (89.55) | 186 (92.54) |
| TRANSFAC_Pred | 54 (34.18) | 48 (30.38) | 72 (45.57) | 158 | 44 (27.85) | 62 (39.24) | 124 (78.48) | 141 (89.24) |
| JASPAR_Pred | 54 (48.65) | 50 (45.05) | 61 (54.95) | 44 (39.64) | 111 | 50 (45.05) | 105 (94.59) | 110 (99.1) |
| MotifMap_Pred | 63 (31.03) | 52 (25.62) | 92 (45.32) | 62 (30.54) | 50 (24.63) | 203 | 176 (86.7) | 179 (88.18) |
| Human_TFTG_DB | 136 (20.27) | 100 (14.9) | 180 (26.83) | 124 (18.48) | 105 (15.65) | 176 (26.23) | 671 | 634 (94.49) |
| Animal_TFDB | 154 (9.4) | 122 (7.44) | 186 (11.35) | 141 (8.6) | 110 (6.71) | 179 (10.92) | 634 (38.68) | 1639 |

**Table S6: GTEx a-cTF phenotype associations**

| GTEx a-cTF | cTF aSNPs | phenotype | P value | beta | GWAS |
|---|---|---|---|---|---|
| STAT2 | 12_56746558; 12_56753822; 12_56653560; 12_56696545 | Mean platelet (thrombocyte) volume | 9.50E-19 | -0.0455 | UK Biobank Neale Lab GWAS |
| | | Standing height | 3.00E-18 | 0.259 | UK Biobank Neale Lab GWAS |
| | | Forced vital capacity (fvc), best measure | 1.80E-12 | 0.0263 | UK Biobank Neale Lab GWAS |
| | | Forced expiratory volume in 1-second (fev1), predicted | 1.60E-12 | 0.0126 | UK Biobank Neale Lab GWAS |
| | | Trunk predicted mass | 3.20E-10 | 0.0922 | UK Biobank Neale Lab GWAS |
| | | Trunk fat-free mass | 4.00E-10 | 0.0956 | UK Biobank Neale Lab GWAS |
| | | Forced vital capacity (fvc) | 1.00E-09 | 0.0224 | UK Biobank Neale Lab GWAS |
| | | Forced expiratory volume in 1-second (fev1), best measure | 7.40E-09 | 0.0176 | UK Biobank Neale Lab GWAS |
| | | Height | 1.00E-13 | 0.054 | https://doi.org/10.1038/nature09410 |
| | | Psoriasis | 5.00E-12 | NA | https://doi.org/10.1038/ncomms7916 |
| | | Psoriasis vulgaris | 5.00E-17 | NA | https://doi.org/10.1016/j.ajhg.2015.10.019 |
| | | Psoriasis | 4.00E-15 | NA | https://doi.org/10.1038/ng.2467 |
| | | Mean platelet volume | 2.40E-08 | -0.0403 | https://doi.org/10.1016/j.cell.2016.10.042 |
| | | Whole body fat-free mass | 4.70E-08 | 0.162 | UK Biobank Neale Lab GWAS |
| TFEB | 6_41700858; 6_41673067 | Mean sphered cell volume | 3.40E-29 | 0.287 | UK Biobank Neale Lab GWAS |
| | | Mean corpuscular volume | 6.40E-29 | 0.235 | UK Biobank Neale Lab GWAS |
| | | Mean corpuscular haemoglobin | 2.70E-22 | 0.0852 | UK Biobank Neale Lab GWAS |
| | | Mean reticulocyte volume | 5.00E-22 | 0.365 | UK Biobank Neale Lab GWAS |
| | | Mean corpuscular volume | 1.10E-16 | 0.0593 | https://doi.org/10.1016/j.cell.2016.10.042 |
| | | Mean corpuscular hemoglobin | 2.20E-11 | 0.048 | https://doi.org/10.1016/j.cell.2016.10.042 |
| TULP2 | 19_49382969; 19_49361441; 19_49391770 | Reticulocyte fraction of red cells | 4.60E-12 | 0.0325 | https://doi.org/10.1016/j.cell.2016.10.042 |
| | | High light scatter reticulocyte count | 7.70E-12 | 0.00022 | UK Biobank Neale Lab GWAS |
| | | Reticulocyte count | 2.70E-10 | 0.0297 | https://doi.org/10.1016/j.cell.2016.10.042 |
| | | High light scatter reticulocyte percentage of red cells | 2.30E-09 | 0.0281 | https://doi.org/10.1016/j.cell.2016.10.042 |
| | | High light scatter reticulocyte count | 1.40E-08 | 0.0266 | https://doi.org/10.1016/j.cell.2016.10.042 |
| | | Hand grip strength (right) | 4.80E-08 | -0.128 | UK Biobank Neale Lab GWAS |
| CCDC88A | 2_55681517 | Standing height | 2.40E-08 | -0.191 | UK Biobank Neale Lab GWAS |
| ZNF483 | 9_114238010; 9_114322499 | Monocyte count | 1.90E-17 | -0.0623 | https://doi.org/10.1016/j.cell.2016.10.042 |
| | | Monocyte percentage | 7.00E-15 | -0.102 | UK Biobank Neale Lab GWAS |
| | | Monocyte percentage of white cells | 2.10E-11 | -0.049 | https://doi.org/10.1016/j.cell.2016.10.042 |
| | | Monocyte count | 5.40E-09 | -0.00626 | UK Biobank Neale Lab GWAS |
| | | Granulocyte percentage of myeloid white cells | 4.90E-08 | 0.04 | https://doi.org/10.1016/j.cell.2016.10.042 |
| ZFP36L1 | 14_69055743; 14_68909571 | Breast cancer | 4.70E-11 | -0.0487 | https://doi.org/10.1038/nature24284 |
| FOXC1 | 6_1623627 | Standing height | 9.10E-12 | 0.151 | UK Biobank Neale Lab GWAS |
| | | Sitting height | 5.00E-10 | 0.0794 | UK Biobank Neale Lab GWAS |
| HOXA13 | 7_27935254 | Standing height | 3.20E-09 | 0.179 | UK Biobank Neale Lab GWAS |
| PLEK | 2_68475595 | Varicose veins of lower extremity | 7.40E-28 | 0.158 | UKB SAIGE |
| | | Varicose veins | 2.60E-27 | 0.154 | UKB SAIGE |

**Table S6: GTEx a-cTF phenotype associations**

| GTEx a-cTF | cTF aSNPs | phenotype | P value | beta | GWAS |
|---|---|---|---|---|---|
| ZNF592 | 15_85203704 | Standing height | 5.00E-14 | 0.134 | UK Biobank Neale Lab GWAS |
| | | Comparative height size at age 10 | 2.40E-10 | 0.0122 | UK Biobank Neale Lab GWAS |
| ATOH7 | 10_70028315 | Sitting height | 1.10E-19 | -0.107 | UK Biobank Neale Lab GWAS |
| | | Standing height | 1.30E-14 | -0.157 | UK Biobank Neale Lab GWAS |
| ZBTB20 | 3_114055430 | Pattern 1 \| hair/balding pattern | 4.10E-12 | 0.163 | UK Biobank Neale Lab GWAS |
| ZKSCAN4 | 6_27361242 | Mean corpuscular volume | 1.50E-24 | -0.199 | UK Biobank Neale Lab GWAS |
| | | Mean corpuscular haemoglobin | 2.40E-24 | -0.0825 | UK Biobank Neale Lab GWAS |
| | | Standing height | 4.40E-23 | -0.275 | UK Biobank Neale Lab GWAS |
| | | Mean corpuscular hemoglobin | 3.60E-18 | -0.0577 | https://doi.org/10.1016/j.cell.2016.10.042 |
| | | Comparative height size at age 10 | 3.40E-16 | -0.0245 | UK Biobank Neale Lab GWAS |
| | | Sitting height | 1.30E-15 | -0.129 | UK Biobank Neale Lab GWAS |
| | | Mean corpuscular volume | 2.50E-14 | -0.0505 | https://doi.org/10.1016/j.cell.2016.10.042 |
| | | Trunk fat mass | 1.50E-12 | -0.161 | UK Biobank Neale Lab GWAS |
| | | Weight | 3.60E-12 | -0.433 | UK Biobank Neale Lab GWAS |
| | | Mean sphered cell volume | 7.10E-12 | -0.162 | UK Biobank Neale Lab GWAS |
| | | Hip circumference | 6.00E-11 | -0.263 | UK Biobank Neale Lab GWAS |
| | | Basal metabolic rate | 9.10E-11 | -24 | UK Biobank Neale Lab GWAS |
| | | Arm fat-free mass (right) | 2.40E-10 | -0.0123 | UK Biobank Neale Lab GWAS |
| | | Arm fat-free mass (left) | 2.40E-10 | -0.0132 | UK Biobank Neale Lab GWAS |
| | | Arm predicted mass (right) | 2.90E-10 | -0.0115 | UK Biobank Neale Lab GWAS |
| | | Whole body fat mass | 3.80E-10 | -0.256 | UK Biobank Neale Lab GWAS |
| | | Whole body water mass | 4.00E-10 | -0.128 | UK Biobank Neale Lab GWAS |
| | | Whole body fat-free mass | 4.40E-10 | -0.173 | UK Biobank Neale Lab GWAS |
| | | Trunk fat-free mass | 2.00E-09 | -0.0857 | UK Biobank Neale Lab GWAS |
| | | Trunk predicted mass | 2.60E-09 | -0.0816 | UK Biobank Neale Lab GWAS |
| | | Leg predicted mass (left) | 2.90E-09 | -0.0298 | UK Biobank Neale Lab GWAS |
| | | Leg fat-free mass (left) | 3.40E-09 | -0.0316 | UK Biobank Neale Lab GWAS |
| | | Arm predicted mass (left) | 3.90E-09 | -0.0115 | UK Biobank Neale Lab GWAS |
| | | Trunk fat percentage | 5.00E-09 | -0.187 | UK Biobank Neale Lab GWAS |
| | | Leg fat-free mass (right) | 5.10E-09 | -0.0312 | UK Biobank Neale Lab GWAS |
| | | Leg predicted mass (right) | 5.20E-09 | -0.0292 | UK Biobank Neale Lab GWAS |

**Table S7: Significant phenotype associations for trans-eQTL aSNPs**

| trans-eQTL aSNPs | phenotype association | P value | Beta | Publication |
|---|---|---|---|---|
| rs10919070 rs10919071 | QT interval | 7.00E-17 | 2.39E+00 | https://doi.org/10.1161/CIRCGEN.117.001758 |
| | QT interval | 3.00E-30 | 1.37E+00 | https://doi.org/10.1038/ng.362 |
| | QT interval | 5.00E-17 | 2.41E+00 | https://doi.org/10.1038/s41586-019-1310-4 |
| | QT interval | 1.00E-31 | 1.68E+00 | https://doi.org/10.1038/ng.3014 |
| | Prothrombin time | 4.70E-27 | 1.50E-01 | https://doi.org/10.1038/s41588-018-0047-6 |
| | Activated partial thromboplastin time | 9.00E-13 | 1.20E-01 | https://doi.org/10.1038/s41588-018-0047-6 |
| rs13063635 rs13098911 | Monocyte percentage | 1.70E-30 | -1.24E-01 | UK Biobank Neale Lab GWAS |
| | Monocyte percentage of white cells | 3.10E-27 | -6.48E-02 | https://doi.org/10.1016/j.cell.2016.10.042 |
| | Monocyte count | 1.50E-22 | -5.87E-02 | https://doi.org/10.1016/j.cell.2016.10.042 |
| | Mouth ulcers \| mouth/teeth dental problems | 3.30E-21 | -1.39E-01 | UK Biobank Neale Lab GWAS |
| | Monocyte count | 5.90E-21 | -9.26E-03 | UK Biobank Neale Lab GWAS |
| | Granulocyte percentage of myeloid white cells | 8.60E-21 | 5.62E-02 | https://doi.org/10.1016/j.cell.2016.10.042 |
| | Celiac disease | 2.50E-11 | 2.78E-01 | https://doi.org/10.1038/ng.543 |
| | Malabsorption/coeliac disease \| non-cancer illness code, self-reported | 6.00E-11 | 4.42E-01 | UK Biobank Neale Lab GWAS |
| | Macrophage inflammatory protein 1b levels | 5.00E-09 | -1.26E-01 | https://doi.org/10.1016/j.ajhg.2016.11.007 |
| | Monocyte chemoattractant protein-1 levels | 1.60E-10 | 1.33E-01 | https://doi.org/10.1016/j.ajhg.2016.11.007 |
| | Eosinophil percentage of granulocytes | 4.30E-09 | -3.54E-02 | https://doi.org/10.1016/j.cell.2016.10.042 |
| | Intestinal malabsorption (non-celiac) | 8.00E-08 | 3.24E-01 | SAIGE UKB |
| | Neutrophil percentage of granulocytes | 2.00E-08 | 3.38E-02 | https://doi.org/10.1016/j.cell.2016.10.042 |
| | Malabsorption/coeliac disease \| non-cancer illness code, self-reported | 2.80E-08 | 3.36E-01 | UK Biobank Neale Lab GWAS |
| rs56032325 | Mean platelet volume | 2.40E-16 | -5.84E-02 | https://doi.org/10.1016/j.cell.2016.10.042 |
| | Mean platelet (thrombocyte) volume | 3.10E-15 | -4.01E-02 | UK Biobank Neale Lab GWAS |
| | Platelet count | 3.50E-09 | 1.62E+00 | UK Biobank Neale Lab GWAS |
| | Platelet count | 6.20E-09 | 4.17E-02 | https://doi.org/10.1016/j.cell.2016.10.042 |
| | Hypospadias | 2.00E-14 | NA | https://doi.org/10.1038/ng.3063 |
| | Standing height | 4.10E-18 | -2.27E-01 | UK Biobank Neale Lab GWAS |
| | Sitting height | 4.20E-16 | -1.23E-01 | UK Biobank Neale Lab GWAS |
| | Trunk predicted mass | 1.50E-11 | -8.69E-02 | UK Biobank Neale Lab GWAS |

**Table S7: Significant phenotype associations for trans-eQTL aSNPs**

| trans-eQTL aSNPs | phenotype association | P value | Beta | Publication |
|---|---|---|---|---|
| rs7811653 | Forced vital capacity (fvc) | 2.70E-11 | -2.15E-02 | UK Biobank Neale Lab GWAS |
| | Trunk fat-free mass | 3.20E-11 | -8.91E-02 | UK Biobank Neale Lab GWAS |
| | Whole body fat-free mass | 3.00E-10 | -1.64E-01 | UK Biobank Neale Lab GWAS |
| | Whole body water mass | 3.10E-10 | -1.21E-01 | UK Biobank Neale Lab GWAS |
| | Forced vital capacity (fvc), best measure | 4.90E-10 | -2.04E-02 | UK Biobank Neale Lab GWAS |
| | Forced expiratory volume in 1-second (fev1) | 1.10E-09 | -1.52E-02 | UK Biobank Neale Lab GWAS |
| | Arm predicted mass (right) | 2.60E-09 | -1.02E-02 | UK Biobank Neale Lab GWAS |
| | Arm fat-free mass (right) | 6.30E-09 | -1.06E-02 | UK Biobank Neale Lab GWAS |
| | Forced expiratory volume in 1-second (fev1), best measure | 1.00E-08 | -1.54E-02 | UK Biobank Neale Lab GWAS |
| | Basal metabolic rate | 1.80E-08 | -1.96E+01 | UK Biobank Neale Lab GWAS |
| | Leg predicted mass (right) | 2.10E-08 | -2.63E-02 | UK Biobank Neale Lab GWAS |
| | Arm predicted mass (left) | 2.30E-08 | -1.03E-02 | UK Biobank Neale Lab GWAS |
| | Arm fat-free mass (left) | 2.50E-08 | -1.09E-02 | UK Biobank Neale Lab GWAS |
| | Leg fat-free mass (right) | 2.90E-08 | -2.78E-02 | UK Biobank Neale Lab GWAS |
| rs72996113 | Mean sphered cell volume | 1.00E-47 | -3.10E-01 | UK Biobank Neale Lab GWAS |
| | High light scatter reticulocyte count | 1.30E-25 | 4.35E-04 | UK Biobank Neale Lab GWAS |
| | Reticulocyte count | 1.10E-18 | 1.40E-03 | UK Biobank Neale Lab GWAS |
| | Reticulocyte count | 9.70E-18 | 5.20E-02 | https://doi.org/10.1016/j.cell.2016.10.042 |
| | Mean corpuscular volume | 1.50E-17 | -1.50E-01 | UK Biobank Neale Lab GWAS |
| | Reticulocyte fraction of red cells | 2.30E-17 | 5.14E-02 | https://doi.org/10.1016/j.cell.2016.10.042 |
| | Red blood cell (erythrocyte) distribution width | 1.20E-16 | -3.17E-02 | UK Biobank Neale Lab GWAS |
| | Reticulocyte percentage | 1.30E-14 | 2.85E-02 | UK Biobank Neale Lab GWAS |
| | Mean reticulocyte volume | 2.00E-13 | -2.32E-01 | UK Biobank Neale Lab GWAS |
| | Platelet distribution width | 2.40E-13 | 1.52E-02 | UK Biobank Neale Lab GWAS |
| | Mean corpuscular hemoglobin concentration | 7.00E-36 | 4.00E-02 | https://doi.org/10.1016/j.cell.2020.06.045 |
| | Platelet crit | 4.30E-13 | -1.37E-03 | UK Biobank Neale Lab GWAS |
| | Red cell distribution width | 4.10E-12 | -4.14E-02 | https://doi.org/10.1016/j.cell.2016.10.042 |
| | Mean corpuscular haemoglobin concentration | 1.40E-11 | 2.89E-02 | UK Biobank Neale Lab GWAS |

**Table S7: Significant phenotype associations for trans-eQTL aSNPs**

| trans-eQTL aSNPs | phenotype association | P value | Beta | Publication |
|---|---|---|---|---|
| | Lymphocyte percentage | 5.40E-11 | -1.92E-01 | UK Biobank Neale Lab GWAS |
| | High light scatter reticulocyte count | 6.30E-10 | 3.74E-02 | https://doi.org/10.1016/j.cell.2016.10.042 |
| | Plateletcrit | 8.00E-10 | -3.79E-02 | https://doi.org/10.1016/j.cell.2016.10.042 |
| | Lymphocyte counts | 9.40E-10 | -3.72E-02 | https://doi.org/10.1016/j.cell.2016.10.042 |
| | High light scatter reticulocyte percentage of red cells | 1.50E-09 | 3.66E-02 | https://doi.org/10.1016/j.cell.2016.10.042 |
| | High light scatter reticulocyte percentage | 2.80E-09 | 8.21E-03 | UK Biobank Neale Lab GWAS |
| | Mean corpuscular hemoglobin concentration | 1.50E-08 | 3.31E-02 | https://doi.org/10.1016/j.cell.2016.10.042 |
| | Neutrophill percentage | 2.40E-08 | 1.89E-01 | UK Biobank Neale Lab GWAS |
| rs2066807<br>rs2066819 | Mean platelet (thrombocyte) volume | 9.50E-19 | -4.55E-02 | UK Biobank Neale Lab GWAS |
| | Standing height | 3.00E-18 | 2.59E-01 | UK Biobank Neale Lab GWAS |
| | Forced vital capacity (fvc), best measure | 1.80E-12 | 2.63E-02 | UK Biobank Neale Lab GWAS |
| | Forced expiratory volume in 1-second (fev1), predicted | 1.60E-12 | 1.26E-02 | UK Biobank Neale Lab GWAS |
| | Trunk predicted mass | 3.20E-10 | 9.22E-02 | UK Biobank Neale Lab GWAS |
| | Trunk fat-free mass | 4.00E-10 | 9.56E-02 | UK Biobank Neale Lab GWAS |
| | Forced vital capacity (fvc) | 1.00E-09 | 2.24E-02 | UK Biobank Neale Lab GWAS |
| | Forced expiratory volume in 1-second (fev1), best measure | 7.40E-09 | 1.76E-02 | UK Biobank Neale Lab GWAS |
| | Height | 1.00E-13 | 5.40E-02 | https://doi.org/10.1038/nature09410 |
| | Psoriasis | 5.00E-12 | NA | https://doi.org/10.1038/ncomms7916 |
| | Psoriasis vulgaris | 5.00E-17 | NA | https://doi.org/10.1016/j.ajhg.2015.10.019 |
| | Psoriasis | 4.00E-15 | NA | https://doi.org/10.1038/ng.2467 |
| | Mean platelet volume | 2.40E-08 | -4.03E-02 | https://doi.org/10.1016/j.cell.2016.10.042 |
| | Whole body fat-free mass | 4.70E-08 | 1.62E-01 | UK Biobank Neale Lab GWAS |
| rs12603526 | Colorectal cancer | 3.00E-08 | NA | https://doi.org/10.1038/ng.2985 |
| | Colorectal cancer | 3.00E-08 | NA | https://doi.org/10.1053/j.gastro.2016.02.076 |
| rs11650665 | Immature reticulocyte fraction | 6.80E-19 | -1.42E-03 | UK Biobank Neale Lab GWAS |
| | Immature fraction of reticulocytes | 1.30E-17 | -3.32E-02 | https://doi.org/10.1016/j.cell.2016.10.042 |
| | High light scatter reticulocyte percentage of red cells | 2.80E-14 | -2.98E-02 | https://doi.org/10.1016/j.cell.2016.10.042 |
| | High light scatter reticulocyte count | 1.90E-13 | -2.88E-02 | https://doi.org/10.1016/j.cell.2016.10.042 |

**Table S7: Significant phenotype associations for trans-eQTL aSNPs**

| trans-eQTL aSNPs | phenotype association | P value | Beta | Publication |
|---|---|---|---|---|
| | Eosinophill count | 2.30E-09 | -1.15E-02 | UK Biobank Neale Lab GWAS |
| | High light scatter reticulocyte count | 8.90E-09 | -1.55E-04 | UK Biobank Neale Lab GWAS |
| rs72973711 | Monocyte percentage | 1.40E-26 | 1.36E-01 | UK Biobank Neale Lab GWAS |
| | Monocyte percentage of white cells | 2.60E-18 | 6.29E-02 | https://doi.org/10.1016/j.cell.2016.10.042 |
| | Neutrophill count | 6.50E-16 | -5.43E-02 | UK Biobank Neale Lab GWAS |
| | Granulocyte percentage of myeloid white cells | 6.90E-15 | -5.63E-02 | https://doi.org/10.1016/j.cell.2016.10.042 |
| | White blood cell (leukocyte) count | 2.30E-12 | -6.82E-02 | UK Biobank Neale Lab GWAS |
| | Neutrophill percentage | 2.60E-10 | -2.53E-01 | UK Biobank Neale Lab GWAS |
| | Neutrophil count | 1.00E-11 | NA | https://doi.org/10.1016/j.cell.2020.06.045 |
| | Red cell distribution width | 6.00E-21 | 3.60E-02 | https://doi.org/10.1016/j.cell.2020.06.045 |
| | White blood cell count | 1.00E-17 | 3.20E-02 | https://doi.org/10.1016/j.cell.2020.06.045 |
| | White blood cell count | 4.00E-16 | NA | https://doi.org/10.1016/j.cell.2020.06.045 |
| rs4805834 | Creatinine levels | 5.00E-11 | -1.00E-02 | https://doi.org/10.1038/ng.566 |
| | Red blood cell (erythrocyte) count | 2.60E-10 | 7.43E-03 | UK Biobank Neale Lab GWAS |
| | Haematocrit percentage | 5.90E-10 | 6.00E-02 | UK Biobank Neale Lab GWAS |
| | Haemoglobin concentration | 6.70E-10 | 2.03E-02 | UK Biobank Neale Lab GWAS |
| rs16997087 | Brain connectivity | 1.00E-10 | NA | https://doi.org/10.1073/pnas.1216206110 |
| rs12908161 | Standing height | 2.20E-27 | 1.60E-02 | UK Biobank Neale Lab GWAS |
| | Sitting height | 4.80E-12 | 1.17E-02 | UK Biobank Neale Lab GWAS |
| | Comparative height size at age 10 | 4.94E-11 | 1.23E-02 | UK Biobank Neale Lab GWAS |
| | Creatinine | 1.06E-10 | -1.42E-02 | UK Biobank Neale Lab GWAS |
| | Trunk fat mass | 1.50E-10 | 1.46E-02 | UK Biobank Neale Lab GWAS |
| | Schizophrenia | 9.41E-10 | -7.41E-02 | https://doi.org/10.1038/nature13595 |
| | Trunk fat percentage | 1.20E-09 | 1.28E-02 | UK Biobank Neale Lab GWAS |
| | Forced expiratory volume in 1-second (FEV1) | 6.40E-09 | 1.14E-02 | UK Biobank Neale Lab GWAS |
| | Weight | 1.90E-08 | 1.11E-02 | UK Biobank Neale Lab GWAS |
| | Weight | 3.50E-08 | 1.08E-02 | UK Biobank Neale Lab GWAS |
| | Forced expiratory volume in 1-second (FEV1), predicted | 4.80E-08 | 1.43E-02 | UK Biobank Neale Lab GWAS |

**Table S7: Significant phenotype associations for trans-eQTL aSNPs**

| trans-eQTL aSNPs | phenotype association | P value | Beta | Publication |
|---|---|---|---|---|
| rs13043612 | Reticulocyte percentage | 3.38E-100 | -5.43E-02 | UK Biobank Neale Lab GWAS |
| | High light scatter reticulocyte percentage | 9.56E-99 | -5.39E-02 | UK Biobank Neale Lab GWAS |
| | High light scatter reticulocyte count | 7.19E-96 | -5.27E-02 | UK Biobank Neale Lab GWAS |
| | Reticulocyte count | 2.45E-94 | -5.20E-02 | UK Biobank Neale Lab GWAS |
| | Reticulocyte fraction of red cells | 2.16E-62 | -6.38E-02 | https://doi.org/10.1016/j.cell.2016.10.042 |
| | Reticulocyte count | 2.07E-57 | -6.12E-02 | https://doi.org/10.1016/j.cell.2016.10.042 |
| | High light scatter reticulocyte percentage of red cells | 3.77E-45 | -5.39E-02 | https://doi.org/10.1016/j.cell.2016.10.042 |
| | High light scatter reticulocyte count | 4.20E-43 | -5.27E-02 | https://doi.org/10.1016/j.cell.2016.10.042 |
| | Immature reticulocyte fraction | 2.86E-34 | -3.12E-02 | UK Biobank Neale Lab GWAS |
| | Immature fraction of reticulocytes | 1.24E-11 | -2.57E-02 | https://doi.org/10.1016/j.cell.2016.10.042 |
| | Lymphocyte count | 5.43E-11 | 1.66E-02 | UK Biobank Neale Lab GWAS |
| | lymphocyte cell count | 6.87E-11 | 1.33E-02 | ieu-b-32 |
| rs6784615 | CD80 on monocyte | 2.46E-47 | 6.59E-01 | https://doi.org/10.1038/s41588-020-0684-4 |
| | CD80 on CD62L+ myeloid Dendritic Cell | 3.72E-24 | 4.63E-01 | https://doi.org/10.1038/s41588-020-0684-4 |
| | CD80 on myeloid Dendritic Cell | 1.22E-18 | 4.03E-01 | https://doi.org/10.1038/s41588-020-0684-4 |
| | HDL cholesterol | 4.50E-17 | -3.36E-02 | https://doi.org/10.1371/journal.pmed.1003062 |
| | triglycerides | 1.50E-16 | 3.45E-02 | https://doi.org/10.1371/journal.pmed.1003062 |
| | HDL cholesterol | 8.67E-15 | -1.42E-02 | UK Biobank Neale Lab GWAS |
| | Triglycerides | 3.98E-14 | 3.68E-02 | UK Biobank Neale Lab GWAS |
| | apolipoprotein A-I | 6.30E-14 | -3.10E-02 | https://doi.org/10.1371/journal.pmed.1003062 |
| | HDL cholesterol | 1.09E-13 | -3.55E-02 | UK Biobank Neale Lab GWAS |
| | Apoliprotein A | 3.75E-11 | -8.71E-03 | UK Biobank Neale Lab GWAS |
| | Apoliprotein A | 7.21E-11 | -3.16E-02 | UK Biobank Neale Lab GWAS |
| | Triglycerides | 4.37E-10 | 3.14E-02 | UK Biobank Neale Lab GWAS |
| rs72647484 | Heel bone mineral density | 3.70E-09 | 2.16E-02 | https://doi.org/10.1038/s41588-018-0302-x |

**Table S8: Frequencies of a-cTF aSNPs in present-day populations**

| | 1000 Genomes | | | | | | | | | | | | | | | SGDP |
|---|---|---|---|---|---|---|---|---|---|---|---|---|---|---|---|---|
| | FIN | IBS | CEU | GBR | TSI | CHS | CHB | CDX | JPT | KHV | BEB | STU | GIH | ITU | PJL | PAP |
| 10_70028315 | 0.237 / 0.974 | 0.159 / 0.935 | 0.126 / 0.889 | 0.154 / 0.92 | 0.136 / 0.912 | 0.171 / 0.896 | 0.165 / 0.892 | 0.145 / 0.858 | 0.12 / 0.802 | 0.106 / 0.808 | 0.279 / 0.996 | 0.23 / 0.991 | 0.218 / 0.986 | 0.196 / 0.981 | 0.224 / 0.989 | - / - |
| 10_8561207 | 0.121 / 0.879 | 0.187 / 0.955 | 0.177 / 0.941 | 0.17 / 0.937 | 0.154 / 0.93 | - / - | - / - | - / - | - / - | 0.005 / 0.194 | 0.099 / 0.901 | 0.078 / 0.848 | 0.112 / 0.913 | 0.074 / 0.831 | 0.12 / 0.932 | - / - |
| 11_101133512 | 0.167 / 0.933 | 0.196 / 0.959 | 0.207 / 0.959 | 0.198 / 0.955 | 0.164 / 0.937 | 0.029 / 0.425 | 0.019 / 0.365 | 0.027 / 0.436 | 0.034 / 0.436 | 0.02 / 0.427 | 0.11 / 0.922 | 0.103 / 0.91 | 0.083 / 0.849 | 0.083 / 0.863 | 0.083 / 0.863 | - / - |
| 11_9409633 | 0.056 / 0.674 | 0.051 / 0.683 | 0.066 / 0.728 | 0.077 / 0.765 | 0.065 / 0.746 | - / - | - / - | - / - | - / - | - / - | 0.041 / 0.67 | 0.059 / 0.77 | 0.083 / 0.849 | 0.049 / 0.714 | 0.068 / 0.815 | - / - |
| 11_9432090 | 0.061 / 0.7 | 0.051 / 0.683 | 0.066 / 0.728 | 0.077 / 0.765 | 0.065 / 0.746 | - / - | - / - | - / - | - / - | - / - | 0.041 / 0.67 | 0.059 / 0.77 | 0.083 / 0.849 | 0.049 / 0.714 | 0.068 / 0.815 | - / - |
| 11_9480577 | 0.061 / 0.7 | 0.051 / 0.683 | 0.071 / 0.751 | 0.082 / 0.784 | 0.07 / 0.767 | - / - | - / - | - / - | - / - | - / - | 0.035 / 0.621 | 0.054 / 0.744 | 0.083 / 0.849 | 0.044 / 0.685 | 0.052 / 0.74 | - / - |
| 11_9487247 | 0.076 / 0.766 | 0.056 / 0.708 | 0.076 / 0.772 | 0.082 / 0.784 | 0.075 / 0.783 | - / - | - / - | - / - | - / - | - / - | 0.035 / 0.621 | 0.054 / 0.744 | 0.083 / 0.849 | 0.044 / 0.685 | 0.073 / 0.833 | - / - |
| 11_9504482 | 0.076 / 0.766 | 0.056 / 0.708 | 0.076 / 0.772 | 0.082 / 0.784 | 0.07 / 0.767 | - / - | - / - | - / - | - / - | 0.005 / 0.194 | 0.035 / 0.621 | 0.054 / 0.744 | 0.083 / 0.849 | 0.044 / 0.685 | 0.073 / 0.833 | - / - |
| 11_9508407 | 0.076 / 0.766 | 0.056 / 0.708 | 0.076 / 0.772 | 0.082 / 0.784 | 0.07 / 0.767 | - / - | - / - | - / - | - / - | - / - | 0.035 / 0.621 | 0.054 / 0.744 | 0.083 / 0.849 | 0.044 / 0.685 | 0.073 / 0.833 | - / - |
| 11_9509352 | 0.076 / 0.766 | 0.056 / 0.708 | 0.076 / 0.772 | 0.082 / 0.784 | 0.07 / 0.767 | - / - | - / - | - / - | - / - | 0.005 / 0.194 | 0.035 / 0.621 | 0.054 / 0.744 | 0.083 / 0.849 | 0.044 / 0.685 | 0.073 / 0.833 | - / - |
| 11_9516186 | 0.076 / 0.766 | 0.056 / 0.708 | 0.076 / 0.772 | 0.082 / 0.784 | 0.07 / 0.767 | - / - | - / - | - / - | - / - | - / - | 0.035 / 0.621 | 0.054 / 0.744 | 0.083 / 0.849 | 0.044 / 0.685 | 0.073 / 0.833 | - / - |
| 11_9536999 | 0.076 / 0.766 | 0.056 / 0.708 | 0.076 / 0.772 | 0.082 / 0.784 | 0.07 / 0.767 | - / - | - / - | - / - | - / - | - / - | 0.035 / 0.621 | 0.054 / 0.744 | 0.083 / 0.849 | 0.044 / 0.685 | 0.068 / 0.815 | - / - |
| 11_9547540 | 0.076 / 0.766 | 0.056 / 0.708 | 0.076 / 0.772 | 0.082 / 0.784 | 0.07 / 0.767 | - / - | - / - | - / - | - / - | 0.005 / 0.194 | 0.035 / 0.621 | 0.054 / 0.744 | 0.083 / 0.849 | 0.044 / 0.685 | 0.068 / 0.815 | - / - |
| 12_133750212 | 0.242 / 0.976 | 0.243 / 0.977 | 0.288 / 0.987 | 0.22 / 0.966 | 0.234 / 0.975 | 0.133 / 0.837 | 0.107 / 0.789 | 0.091 / 0.741 | 0.077 / 0.67 | 0.152 / 0.888 | 0.233 / 0.992 | 0.235 / 0.992 | 0.248 / 0.99 | 0.294 / 0.997 | 0.266 / 0.994 | - / - |
| 12_133808129 | 0.177 / 0.942 | 0.21 / 0.965 | 0.258 / 0.98 | 0.17 / 0.937 | 0.201 / 0.961 | 0.143 / 0.854 | 0.097 / 0.763 | 0.097 / 0.757 | 0.067 / 0.627 | 0.146 / 0.881 | 0.221 / 0.99 | 0.196 / 0.984 | 0.204 / 0.981 | 0.275 / 0.995 | 0.203 / 0.985 | - / - |
| 12_56653560 | 0.076 / 0.766 | 0.047 / 0.656 | 0.076 / 0.772 | 0.055 / 0.665 | 0.079 / 0.799 | 0.024 / 0.387 | 0.029 / 0.447 | 0.027 / 0.436 | 0.058 / 0.582 | 0.04 / 0.567 | 0.023 / 0.501 | 0.025 / 0.509 | 0.024 / 0.488 | 0.01 / 0.275 | 0.016 / 0.384 | - / - |
| 12_56696545 | 0.081 / 0.785 | 0.051 / 0.683 | 0.086 / 0.806 | 0.055 / 0.665 | 0.079 / 0.799 | 0.024 / 0.387 | 0.029 / 0.447 | 0.027 / 0.436 | 0.058 / 0.582 | 0.04 / 0.567 | 0.029 / 0.566 | 0.025 / 0.509 | 0.024 / 0.488 | 0.01 / 0.275 | 0.016 / 0.384 | - / - |
| 12_56746558 | 0.071 / 0.745 | 0.042 / 0.625 | 0.076 / 0.772 | 0.055 / 0.665 | 0.079 / 0.799 | 0.024 / 0.387 | 0.024 / 0.405 | 0.032 / 0.479 | 0.053 / 0.555 | 0.04 / 0.567 | 0.023 / 0.501 | 0.025 / 0.509 | 0.024 / 0.488 | 0.01 / 0.275 | 0.016 / 0.384 | - / - |
| 12_56753822 | 0.071 / 0.745 | 0.042 / 0.625 | 0.076 / 0.772 | 0.049 / 0.629 | 0.079 / 0.799 | 0.024 / 0.387 | 0.024 / 0.405 | 0.032 / 0.479 | 0.053 / 0.555 | 0.04 / 0.567 | 0.023 / 0.501 | 0.025 / 0.509 | 0.024 / 0.488 | 0.01 / 0.275 | 0.016 / 0.384 | 0.5 / 0.949 |
| 12_69669013 | 0.03 / 0.505 | 0.051 / 0.683 | 0.076 / 0.772 | 0.049 / 0.629 | 0.065 / 0.746 | - / - | - / - | - / - | - / - | - / - | 0.006 / 0.182 | 0.015 / 0.372 | 0.044 / 0.665 | 0.005 / 0.163 | 0.021 / 0.464 | - / - |
| 13_27959981 | 0.167 / 0.933 | 0.187 / 0.955 | 0.167 / 0.934 | 0.148 / 0.912 | 0.178 / 0.948 | - / - | - / - | - / - | - / - | - / - | 0.023 / 0.501 | 0.01 / 0.281 | 0.029 / 0.54 | 0.02 / 0.445 | 0.031 / 0.582 | - / - |
| 13_27965764 | 0.162 / 0.928 | 0.187 / 0.955 | 0.167 / 0.934 | 0.148 / 0.912 | 0.168 / 0.94 | - / - | - / - | - / - | - / - | - / - | 0.023 / 0.501 | 0.01 / 0.281 | 0.029 / 0.54 | 0.02 / 0.445 | 0.031 / 0.582 | - / - |
| 13_27998646 | 0.167 / 0.933 | 0.182 / 0.95 | 0.172 / 0.938 | 0.148 / 0.912 | 0.159 / 0.934 | - / - | - / - | - / - | - / - | - / - | 0.023 / 0.501 | 0.015 / 0.372 | 0.044 / 0.665 | 0.02 / 0.445 | 0.031 / 0.582 | - / - |
| 14_68909571 | 0.212 / 0.963 | 0.229 / 0.973 | 0.237 / 0.973 | 0.242 / 0.975 | 0.173 / 0.944 | 0.048 / 0.55 | 0.058 / 0.62 | 0.016 / 0.331 | 0.034 / 0.436 | 0.01 / 0.293 | 0.035 / 0.621 | 0.078 / 0.848 | 0.087 / 0.861 | 0.093 / 0.888 | 0.156 / 0.966 | 0.036 / 0.196 |
| 14_69055743 | 0.141 / 0.908 | 0.266 / 0.984 | 0.283 / 0.985 | 0.154 / 0.92 | 0.234 / 0.975 | 0.095 / 0.751 | 0.092 / 0.748 | 0.054 / 0.595 | 0.13 / 0.823 | 0.051 / 0.626 | 0.023 / 0.501 | 0.059 / 0.77 | 0.068 / 0.795 | 0.088 / 0.876 | 0.109 / 0.919 | 0.25 / 0.771 |
| 14_74204686 | 0.288 / 0.987 | 0.276 / 0.986 | 0.343 / 0.993 | 0.324 / 0.992 | 0.294 / 0.989 | - / - | 0.005 / 0.17 | - / - | - / - | 0.01 / 0.293 | 0.064 / 0.8 | 0.064 / 0.793 | 0.107 / 0.906 | 0.049 / 0.714 | 0.073 / 0.833 | - / - |
| 14_76780815 | 0.212 / 0.963 | 0.154 / 0.931 | 0.192 / 0.951 | 0.121 / 0.877 | 0.187 / 0.953 | - / - | 0.005 / 0.17 | 0.005 / 0.155 | - / - | - / - | 0.047 / 0.709 | 0.049 / 0.714 | 0.058 / 0.75 | 0.034 / 0.608 | 0.115 / 0.927 | - / - |
| 15_85203704 | 0.212 / 0.963 | 0.266 / 0.984 | 0.258 / 0.98 | 0.214 / 0.964 | 0.224 / 0.972 | 0.067 / 0.645 | 0.063 / 0.639 | 0.054 / 0.595 | 0.038 / 0.47 | 0.076 / 0.727 | 0.151 / 0.964 | 0.137 / 0.954 | 0.121 / 0.928 | 0.176 / 0.974 | 0.135 / 0.95 | - / - |
| 16_3530658 | 0.182 / 0.945 | 0.164 / 0.938 | 0.222 / 0.966 | 0.187 / 0.95 | 0.262 / 0.982 | 0.029 / 0.425 | 0.049 / 0.573 | 0.011 / 0.254 | 0.053 / 0.555 | 0.056 / 0.651 | 0.076 / 0.845 | 0.093 / 0.889 | 0.058 / 0.75 | 0.098 / 0.898 | 0.109 / 0.919 | - / - |
| 19_11952311 | 0.015 / 0.329 | 0.065 / 0.753 | 0.03 / 0.505 | 0.022 / 0.395 | 0.056 / 0.704 | 0.071 / 0.666 | 0.083 / 0.715 | 0.102 / 0.772 | 0.072 / 0.651 | 0.056 / 0.651 | 0.035 / 0.621 | 0.02 / 0.447 | 0.01 / 0.265 | 0.039 / 0.65 | 0.031 / 0.582 | 0.179 / 0.662 |
| 19_11998756 | - / - | 0.047 / 0.656 | 0.02 / 0.398 | 0.005 / 0.146 | 0.037 / 0.593 | 0.01 / 0.222 | 0.01 / 0.253 | 0.011 / 0.254 | - / - | - / - | 0.023 / 0.501 | 0.01 / 0.281 | 0.024 / 0.488 | 0.029 / 0.562 | 0.031 / 0.582 | - / - |
| 19_3793149 | 0.005 / 0.145 | 0.028 / 0.516 | 0.04 / 0.586 | 0.027 / 0.45 | 0.047 / 0.652 | - / - | - / - | - / - | - / - | - / - | 0.047 / 0.709 | 0.049 / 0.714 | 0.01 / 0.265 | 0.034 / 0.608 | 0.094 / 0.889 | - / - |
| 19_49361441 | 0.167 / 0.933 | 0.168 / 0.941 | 0.172 / 0.938 | 0.198 / 0.955 | 0.14 / 0.917 | - / - | - / - | - / - | - / - | - / - | 0.041 / 0.67 | 0.025 / 0.509 | 0.087 / 0.861 | 0.054 / 0.742 | 0.099 / 0.899 | - / - |
| 19_49382969 | 0.167 / 0.933 | 0.164 / 0.938 | 0.167 / 0.934 | 0.176 / 0.942 | 0.154 / 0.93 | - / - | 0.005 / 0.17 | - / - | - / - | - / - | 0.047 / 0.709 | 0.074 / 0.831 | 0.112 / 0.913 | 0.069 / 0.813 | 0.115 / 0.927 | - / - |
| 19_49391770 | 0.111 / 0.862 | 0.164 / 0.938 | 0.157 / 0.925 | 0.159 / 0.926 | 0.121 / 0.897 | - / - | 0.005 / 0.17 | - / - | - / - | - / - | 0.047 / 0.709 | 0.074 / 0.831 | 0.097 / 0.886 | 0.059 / 0.767 | 0.099 / 0.899 | - / - |
| 19_52387186 | - / - | 0.033 / 0.561 | 0.005 / 0.152 | 0.011 / 0.243 | 0.019 / 0.405 | - / - | 0.01 / 0.253 | 0.005 / 0.155 | - / - | 0.005 / 0.194 | 0.017 / 0.419 | 0.005 / 0.166 | 0.015 / 0.354 | 0.02 / 0.445 | 0.01 / 0.285 | - / - |
| 1_159888701 | 0.005 / 0.145 | 0.014 / 0.341 | 0.005 / 0.152 | 0.022 / 0.395 | 0.009 / 0.258 | 0.01 / 0.222 | 0.01 / 0.253 | 0.005 / 0.155 | 0.019 / 0.308 | 0.01 / 0.293 | 0.099 / 0.901 | 0.127 / 0.944 | 0.068 / 0.795 | 0.137 / 0.952 | 0.125 / 0.939 | - / - |
| 1_165316371 | 0.086 / 0.8 | 0.103 / 0.862 | 0.086 / 0.806 | 0.093 / 0.817 | 0.061 / 0.726 | 0.11 / 0.787 | 0.17 / 0.898 | 0.075 / 0.688 | 0.178 / 0.896 | 0.096 / 0.782 | 0.233 / 0.992 | 0.176 / 0.977 | 0.184 / 0.975 | 0.127 / 0.942 | 0.13 / 0.945 | - / - |
| 1_170634895 | 0.121 / 0.879 | 0.131 / 0.904 | 0.101 / 0.845 | 0.093 / 0.817 | 0.089 / 0.827 | - / - | - / - | - / - | - / - | - / - | 0.041 / 0.67 | 0.02 / 0.447 | 0.044 / 0.665 | 0.044 / 0.685 | 0.036 / 0.628 | - / - |
| 1_205456751 | 0.051 / 0.647 | 0.061 / 0.731 | 0.081 / 0.789 | 0.066 / 0.72 | 0.037 / 0.593 | 0.024 / 0.387 | 0.034 / 0.484 | 0.011 / 0.254 | 0.019 / 0.308 | 0.035 / 0.538 | 0.093 / 0.889 | 0.034 / 0.606 | 0.058 / 0.75 | 0.059 / 0.767 | 0.031 / 0.582 | - / - |
| 1_213470400 | 0.01 / 0.245 | 0.093 / 0.838 | 0.04 / 0.586 | 0.044 / 0.593 | 0.047 / 0.652 | 0.033 / 0.46 | 0.063 / 0.639 | 0.027 / 0.436 | 0.014 / 0.251 | 0.025 / 0.471 | 0.012 / 0.315 | 0.005 / 0.166 | 0.015 / 0.354 | - / - | 0.01 / 0.285 | - / - |
| 1_41347586 | 0.091 / 0.816 | 0.089 / 0.827 | 0.106 / 0.855 | 0.082 / 0.784 | 0.07 / 0.767 | 0.033 / 0.46 | 0.068 / 0.661 | 0.097 / 0.757 | 0.067 / 0.627 | 0.04 / 0.567 | 0.052 / 0.743 | 0.034 / 0.606 | 0.024 / 0.488 | 0.015 / 0.369 | 0.078 / 0.849 | - / - |
| 1_58900316 | 0.146 / 0.913 | 0.238 / 0.976 | 0.273 / 0.983 | 0.258 / 0.981 | 0.271 / 0.984 | 0.095 / 0.751 | 0.092 / 0.748 | 0.129 / 0.829 | 0.067 / 0.627 | 0.141 / 0.873 | 0.087 / 0.876 | 0.098 / 0.901 | 0.102 / 0.896 | 0.088 / 0.876 | 0.141 / 0.955 | 0.077 / 0.403 |

**Table S8: Frequencies of a-cTF aSNPs in present-day populations**

| | 1000 Genomes | | | | | | | | | | | | | | | SGDP |
|---|---|---|---|---|---|---|---|---|---|---|---|---|---|---|---|---|
| | FIN | IBS | CEU | GBR | TSI | CHS | CHB | CDX | JPT | KHV | BEB | STU | GIH | ITU | PJL | PAP |
| 20_42140933 | 0.025 / 0.453 | 0.009 / 0.258 | 0.04 / 0.586 | 0.022 / 0.395 | 0.014 / 0.333 | - / - | - / - | - / - | - / - | - / - | 0.006 / 0.182 | - / - | 0.005 / 0.155 | - / - | 0.005 / 0.172 | - / - |
| 20_757491 | 0.172 / 0.937 | 0.121 / 0.891 | 0.162 / 0.93 | 0.165 / 0.931 | 0.159 / 0.934 | 0.581 / 0.999 | 0.456 / 0.996 | 0.624 / 0.999 | 0.49 / 0.997 | 0.646 / 1 | 0.215 / 0.989 | 0.162 / 0.97 | 0.204 / 0.981 | 0.172 / 0.973 | 0.245 / 0.992 | - / - |
| 21_43221111 | 0.071 / 0.745 | 0.112 / 0.877 | 0.071 / 0.751 | 0.088 / 0.804 | 0.079 / 0.799 | 0.005 / 0.136 | - / - | 0.005 / 0.155 | - / - | 0.015 / 0.369 | 0.157 / 0.968 | 0.127 / 0.944 | 0.112 / 0.913 | 0.118 / 0.93 | 0.104 / 0.909 | - / - |
| 21_43230111 | 0.071 / 0.745 | 0.107 / 0.869 | 0.066 / 0.728 | 0.088 / 0.804 | 0.079 / 0.799 | 0.005 / 0.136 | - / - | 0.005 / 0.155 | - / - | 0.015 / 0.369 | 0.157 / 0.968 | 0.127 / 0.944 | 0.112 / 0.913 | 0.113 / 0.923 | 0.099 / 0.899 | - / - |
| 22_22822506 | 0.02 / 0.396 | 0.033 / 0.561 | 0.045 / 0.62 | 0.016 / 0.329 | 0.051 / 0.68 | - / - | - / - | - / - | - / - | - / - | - / - | 0.015 / 0.372 | 0.01 / 0.265 | 0.01 / 0.275 | 0.026 / 0.526 | - / - |
| 2_179361908 | 0.051 / 0.647 | 0.019 / 0.408 | 0.04 / 0.586 | 0.033 / 0.505 | 0.019 / 0.405 | 0.052 / 0.575 | 0.063 / 0.639 | 0.038 / 0.513 | 0.019 / 0.308 | 0.035 / 0.538 | 0.006 / 0.182 | - / - | - / - | - / - | - / - | - / - |
| 2_238648521 | - / - | 0.103 / 0.862 | 0.035 / 0.545 | 0.049 / 0.629 | 0.051 / 0.68 | 0.057 / 0.6 | 0.063 / 0.639 | 0.054 / 0.595 | 0.062 / 0.604 | 0.061 / 0.672 | 0.047 / 0.709 | 0.044 / 0.682 | 0.019 / 0.426 | 0.025 / 0.503 | 0.052 / 0.74 | 0.1 / 0.469 |
| 2_55681517 | 0.025 / 0.453 | 0.051 / 0.683 | 0.051 / 0.649 | 0.066 / 0.72 | 0.084 / 0.812 | - / - | - / - | - / - | - / - | - / - | 0.058 / 0.773 | 0.044 / 0.682 | 0.112 / 0.913 | 0.069 / 0.813 | 0.073 / 0.833 | - / - |
| 2_68475595 | 0.242 / 0.975 | 0.388 / 0.997 | 0.409 / 0.997 | 0.429 / 0.998 | 0.355 / 0.996 | 0.348 / 0.986 | 0.291 / 0.975 | 0.403 / 0.993 | 0.433 / 0.994 | 0.328 / 0.985 | 0.244 / 0.993 | 0.206 / 0.986 | 0.15 / 0.954 | 0.172 / 0.97 | 0.115 / 0.92 | - / - |
| 2_86623524 | 0.04 / 0.59 | 0.019 / 0.408 | 0.035 / 0.545 | 0.022 / 0.395 | 0.023 / 0.466 | 0.024 / 0.387 | 0.029 / 0.447 | 0.065 / 0.646 | 0.135 / 0.831 | 0.04 / 0.567 | 0.023 / 0.501 | 0.02 / 0.447 | 0.019 / 0.426 | 0.015 / 0.369 | 0.021 / 0.464 | - / - |
| 3_114055430 | 0.015 / 0.329 | 0.028 / 0.516 | 0.03 / 0.505 | 0.011 / 0.243 | 0.019 / 0.405 | 0.01 / 0.222 | 0.005 / 0.17 | - / - | 0.005 / 0.113 | 0.005 / 0.194 | 0.064 / 0.8 | 0.078 / 0.848 | 0.053 / 0.726 | 0.049 / 0.714 | 0.073 / 0.833 | - / - |
| 3_141891385 | 0.071 / 0.745 | 0.089 / 0.827 | 0.116 / 0.873 | 0.099 / 0.833 | 0.107 / 0.873 | 0.057 / 0.6 | 0.053 / 0.597 | 0.065 / 0.646 | 0.048 / 0.528 | 0.04 / 0.567 | 0.041 / 0.67 | 0.005 / 0.166 | 0.034 / 0.589 | 0.039 / 0.65 | 0.073 / 0.833 | - / - |
| 3_20052020 | 0.283 / 0.986 | 0.304 / 0.991 | 0.303 / 0.99 | 0.302 / 0.989 | 0.285 / 0.988 | 0.014 / 0.282 | 0.01 / 0.253 | - / - | 0.005 / 0.113 | 0.005 / 0.194 | 0.099 / 0.901 | 0.103 / 0.91 | 0.146 / 0.954 | 0.137 / 0.952 | 0.172 / 0.973 | - / - |
| 3_20104838 | 0.237 / 0.974 | 0.262 / 0.983 | 0.278 / 0.984 | 0.28 / 0.986 | 0.154 / 0.93 | 0.005 / 0.136 | 0.019 / 0.365 | 0.011 / 0.254 | - / - | 0.005 / 0.194 | 0.07 / 0.822 | 0.049 / 0.714 | 0.107 / 0.906 | 0.098 / 0.898 | 0.146 / 0.958 | - / - |
| 3_23107748 | 0.04 / 0.59 | 0.065 / 0.753 | 0.025 / 0.451 | 0.055 / 0.665 | 0.028 / 0.516 | 0.105 / 0.774 | 0.136 / 0.851 | 0.075 / 0.688 | 0.173 / 0.891 | 0.141 / 0.873 | 0.058 / 0.773 | 0.034 / 0.606 | 0.039 / 0.631 | 0.054 / 0.742 | 0.062 / 0.793 | - / - |
| 3_24499855 | 0.015 / 0.329 | 0.009 / 0.258 | 0.035 / 0.545 | 0.038 / 0.552 | 0.028 / 0.516 | 0.081 / 0.7 | 0.097 / 0.763 | 0.097 / 0.757 | 0.101 / 0.751 | 0.111 / 0.819 | 0.093 / 0.889 | 0.098 / 0.901 | 0.112 / 0.913 | 0.069 / 0.813 | 0.068 / 0.815 | 0.067 / 0.333 |
| 4_121643726 | 0.141 / 0.908 | 0.252 / 0.98 | 0.192 / 0.951 | 0.28 / 0.986 | 0.238 / 0.976 | 0.095 / 0.751 | 0.097 / 0.763 | 0.043 / 0.543 | 0.091 / 0.721 | 0.086 / 0.756 | 0.07 / 0.822 | 0.049 / 0.714 | 0.044 / 0.665 | 0.074 / 0.831 | 0.078 / 0.849 | - / - |
| 4_121703570 | 0.212 / 0.963 | 0.318 / 0.992 | 0.217 / 0.963 | 0.313 / 0.991 | 0.276 / 0.986 | 0.205 / 0.929 | 0.228 / 0.948 | 0.156 / 0.876 | 0.188 / 0.91 | 0.162 / 0.901 | 0.076 / 0.845 | 0.054 / 0.744 | 0.058 / 0.75 | 0.113 / 0.923 | 0.104 / 0.909 | 0.033 / 0.121 |
| 5_16496850 | 0.01 / 0.245 | 0.028 / 0.516 | 0.04 / 0.586 | 0.038 / 0.552 | 0.033 / 0.558 | - / - | - / - | - / - | - / - | - / - | - / - | 0.015 / 0.372 | 0.024 / 0.488 | - / - | 0.005 / 0.172 | - / - |
| 6_109149871 | 0.045 / 0.621 | 0.023 / 0.467 | 0.035 / 0.545 | 0.033 / 0.505 | 0.028 / 0.516 | - / - | - / - | - / - | - / - | - / - | 0.023 / 0.501 | 0.015 / 0.372 | 0.01 / 0.265 | 0.015 / 0.369 | 0.042 / 0.668 | - / - |
| 6_1364471 | 0.101 / 0.839 | 0.121 / 0.891 | 0.111 / 0.863 | 0.077 / 0.765 | 0.079 / 0.799 | - / - | - / - | - / - | 0.005 / 0.113 | - / - | 0.017 / 0.419 | 0.025 / 0.509 | 0.029 / 0.54 | 0.02 / 0.445 | 0.026 / 0.526 | - / - |
| 6_1367583 | 0.076 / 0.766 | 0.089 / 0.827 | 0.106 / 0.855 | 0.06 / 0.694 | 0.061 / 0.726 | - / - | - / - | - / - | 0.005 / 0.113 | - / - | 0.012 / 0.315 | 0.039 / 0.645 | 0.019 / 0.426 | 0.01 / 0.275 | 0.026 / 0.526 | - / - |
| 6_1623627 | 0.061 / 0.7 | 0.154 / 0.931 | 0.136 / 0.902 | 0.137 / 0.899 | 0.159 / 0.934 | - / - | - / - | - / - | - / - | 0.005 / 0.194 | 0.041 / 0.67 | 0.01 / 0.281 | 0.039 / 0.631 | 0.054 / 0.742 | 0.031 / 0.582 | - / - |
| 6_18921942 | 0.051 / 0.647 | 0.07 / 0.77 | 0.086 / 0.806 | 0.088 / 0.804 | 0.075 / 0.783 | - / - | - / - | 0.005 / 0.155 | - / - | - / - | 0.035 / 0.621 | 0.044 / 0.682 | 0.058 / 0.75 | 0.025 / 0.503 | 0.089 / 0.876 | - / - |
| 6_27361242 | 0.061 / 0.7 | 0.07 / 0.77 | 0.091 / 0.82 | 0.082 / 0.784 | 0.047 / 0.652 | - / - | 0.005 / 0.17 | - / - | - / - | 0.005 / 0.194 | 0.017 / 0.419 | 0.01 / 0.281 | 0.024 / 0.488 | 0.029 / 0.562 | 0.01 / 0.285 | - / - |
| 6_41673067 | 0.157 / 0.924 | 0.145 / 0.922 | 0.116 / 0.873 | 0.143 / 0.906 | 0.145 / 0.921 | 0.005 / 0.136 | - / - | - / - | - / - | 0.005 / 0.194 | 0.035 / 0.621 | 0.078 / 0.848 | 0.063 / 0.774 | 0.054 / 0.742 | 0.052 / 0.74 | 0.1 / 0.469 |
| 6_41700858 | 0.096 / 0.829 | 0.075 / 0.788 | 0.051 / 0.649 | 0.082 / 0.784 | 0.056 / 0.704 | - / - | - / - | - / - | - / - | 0.005 / 0.194 | 0.035 / 0.621 | 0.049 / 0.714 | 0.053 / 0.726 | 0.054 / 0.742 | 0.036 / 0.628 | 0.115 / 0.515 |
| 6_43370915 | 0.035 / 0.552 | 0.131 / 0.904 | 0.096 / 0.832 | 0.099 / 0.833 | 0.089 / 0.827 | 0.005 / 0.136 | - / - | - / - | - / - | 0.025 / 0.471 | 0.105 / 0.913 | 0.083 / 0.863 | 0.112 / 0.913 | 0.059 / 0.767 | 0.073 / 0.833 | 0.033 / 0.121 |
| 7_1596384 | 0.035 / 0.552 | 0.061 / 0.731 | 0.101 / 0.845 | 0.066 / 0.72 | 0.056 / 0.704 | - / - | - / - | - / - | - / - | 0.005 / 0.194 | 0.012 / 0.315 | - / - | 0.01 / 0.265 | 0.005 / 0.163 | 0.005 / 0.172 | - / - |
| 7_27935254 | 0.035 / 0.553 | 0.131 / 0.898 | 0.061 / 0.68 | 0.071 / 0.744 | 0.098 / 0.854 | - / - | - / - | - / - | - / - | 0.01 / 0.295 | 0.006 / 0.183 | 0.039 / 0.646 | 0.019 / 0.355 | 0.005 / 0.164 | 0.005 / 0.001 | - / - |
| 7_43040084 | 0.04 / 0.59 | 0.019 / 0.408 | 0.01 / 0.251 | 0.044 / 0.593 | 0.023 / 0.466 | 0.114 / 0.797 | 0.117 / 0.811 | 0.118 / 0.808 | 0.082 / 0.688 | 0.111 / 0.819 | 0.081 / 0.862 | 0.083 / 0.863 | 0.053 / 0.726 | 0.103 / 0.908 | 0.078 / 0.849 | - / - |
| 7_44790921 | 0.061 / 0.7 | 0.15 / 0.926 | 0.116 / 0.873 | 0.17 / 0.937 | 0.112 / 0.882 | 0.057 / 0.6 | 0.039 / 0.519 | 0.016 / 0.331 | 0.019 / 0.308 | 0.02 / 0.427 | 0.047 / 0.709 | 0.078 / 0.848 | 0.112 / 0.913 | 0.059 / 0.767 | 0.057 / 0.768 | 0.036 / 0.196 |
| 7_44799347 | 0.056 / 0.674 | 0.121 / 0.891 | 0.116 / 0.873 | 0.181 / 0.946 | 0.112 / 0.882 | 0.057 / 0.6 | 0.039 / 0.519 | 0.016 / 0.331 | 0.019 / 0.308 | 0.02 / 0.427 | 0.041 / 0.67 | 0.069 / 0.814 | 0.107 / 0.906 | 0.059 / 0.767 | 0.057 / 0.768 | 0.033 / 0.121 |
| 7_44801682 | 0.056 / 0.674 | 0.121 / 0.891 | 0.116 / 0.873 | 0.181 / 0.946 | 0.112 / 0.882 | 0.057 / 0.6 | 0.039 / 0.519 | 0.016 / 0.331 | 0.019 / 0.308 | 0.02 / 0.427 | 0.041 / 0.67 | 0.069 / 0.814 | 0.107 / 0.906 | 0.059 / 0.767 | 0.057 / 0.768 | 0.045 / 0.249 |
| 7_44803944 | 0.056 / 0.674 | 0.121 / 0.891 | 0.116 / 0.873 | 0.181 / 0.946 | 0.112 / 0.882 | 0.057 / 0.6 | 0.039 / 0.519 | 0.016 / 0.331 | 0.019 / 0.308 | 0.02 / 0.427 | 0.041 / 0.67 | 0.069 / 0.814 | 0.107 / 0.906 | 0.059 / 0.767 | 0.057 / 0.768 | 0.033 / 0.121 |
| 7_55932373 | 0.071 / 0.745 | 0.037 / 0.594 | 0.04 / 0.586 | 0.049 / 0.629 | 0.07 / 0.767 | 0.024 / 0.387 | 0.029 / 0.447 | 0.022 / 0.39 | 0.062 / 0.604 | 0.015 / 0.369 | 0.041 / 0.67 | 0.108 / 0.919 | 0.044 / 0.665 | 0.049 / 0.714 | 0.047 / 0.706 | - / - |
| 7_64321622 | 0.035 / 0.552 | 0.047 / 0.656 | 0.04 / 0.586 | 0.049 / 0.629 | 0.023 / 0.466 | 0.119 / 0.806 | 0.15 / 0.873 | 0.113 / 0.796 | 0.125 / 0.814 | 0.111 / 0.819 | 0.029 / 0.566 | 0.02 / 0.447 | 0.015 / 0.354 | 0.029 / 0.562 | 0.005 / 0.172 | - / - |
| 7_64386343 | 0.035 / 0.552 | 0.047 / 0.656 | 0.04 / 0.586 | 0.049 / 0.629 | 0.023 / 0.466 | 0.119 / 0.806 | 0.155 / 0.879 | 0.118 / 0.808 | 0.12 / 0.802 | 0.126 / 0.849 | 0.029 / 0.566 | 0.015 / 0.372 | 0.015 / 0.354 | 0.029 / 0.562 | 0.005 / 0.172 | - / - |
| 7_64408851 | 0.035 / 0.552 | 0.047 / 0.656 | 0.04 / 0.586 | 0.049 / 0.629 | 0.023 / 0.466 | 0.114 / 0.797 | 0.16 / 0.886 | 0.118 / 0.808 | 0.12 / 0.802 | 0.131 / 0.856 | 0.029 / 0.566 | 0.039 / 0.645 | 0.024 / 0.488 | 0.039 / 0.65 | 0.016 / 0.384 | - / - |
| 7_64777273 | 0.04 / 0.59 | 0.037 / 0.594 | 0.02 / 0.398 | 0.044 / 0.593 | 0.033 / 0.558 | 0.129 / 0.828 | 0.155 / 0.879 | 0.151 / 0.867 | 0.173 / 0.891 | 0.146 / 0.881 | 0.035 / 0.621 | 0.034 / 0.606 | 0.024 / 0.488 | 0.034 / 0.608 | 0.01 / 0.285 | - / - |
| 7_64840402 | 0.106 / 0.85 | 0.084 / 0.815 | 0.071 / 0.751 | 0.099 / 0.833 | 0.098 / 0.853 | 0.114 / 0.797 | 0.141 / 0.858 | 0.129 / 0.829 | 0.149 / 0.857 | 0.121 / 0.839 | 0.041 / 0.67 | 0.029 / 0.562 | 0.034 / 0.589 | 0.049 / 0.714 | 0.031 / 0.582 | - / - |
| 7_6539123 | 0.015 / 0.329 | 0.056 / 0.708 | 0.061 / 0.705 | 0.049 / 0.629 | 0.042 / 0.627 | - / - | - / - | - / - | - / - | - / - | 0.012 / 0.315 | 0.005 / 0.166 | 0.034 / 0.589 | 0.039 / 0.65 | 0.016 / 0.384 | - / - |
| 9_104165291 | 0.051 / 0.647 | 0.098 / 0.85 | 0.066 / 0.728 | 0.104 / 0.844 | 0.07 / 0.767 | - / - | - / - | 0.005 / 0.155 | - / - | 0.01 / 0.293 | 0.047 / 0.709 | 0.083 / 0.863 | 0.083 / 0.849 | 0.044 / 0.685 | 0.104 / 0.909 | - / - |

**Table S8: Frequencies of a-cTF aSNPs in present-day populations**

| | 1000 Genomes | | | | | | | | | | | | | | | SGDP |
|---|---|---|---|---|---|---|---|---|---|---|---|---|---|---|---|---|
| | FIN | IBS | CEU | GBR | TSI | CHS | CHB | CDX | JPT | KHV | BEB | STU | GIH | ITU | PJL | PAP |
| 9_104171856 | 0.051 / 0.647 | 0.098 / 0.85 | 0.071 / 0.751 | 0.104 / 0.844 | 0.07 / 0.767 | - / - | - / - | 0.005 / 0.155 | - / - | 0.005 / 0.194 | 0.047 / 0.709 | 0.083 / 0.863 | 0.083 / 0.849 | 0.039 / 0.65 | 0.104 / 0.909 | - / - |
| 9_114238010 | 0.076 / 0.766 | 0.065 / 0.753 | 0.071 / 0.751 | 0.027 / 0.45 | 0.089 / 0.827 | 0.252 / 0.96 | 0.165 / 0.892 | 0.167 / 0.893 | 0.231 / 0.947 | 0.187 / 0.925 | 0.145 / 0.96 | 0.113 / 0.926 | 0.073 / 0.813 | 0.078 / 0.848 | 0.13 / 0.945 | - / - |
| 9_114322499 | 0.086 / 0.8 | 0.056 / 0.708 | 0.086 / 0.806 | 0.027 / 0.45 | 0.075 / 0.783 | 0.348 / 0.986 | 0.243 / 0.956 | 0.188 / 0.916 | 0.284 / 0.97 | 0.217 / 0.949 | 0.151 / 0.964 | 0.103 / 0.91 | 0.087 / 0.861 | 0.083 / 0.863 | 0.135 / 0.95 | - / - |
| 9_128935738 | 0.116 / 0.87 | 0.145 / 0.922 | 0.131 / 0.896 | 0.187 / 0.95 | 0.107 / 0.873 | 0.005 / 0.136 | 0.005 / 0.17 | - / - | - / - | 0.01 / 0.293 | 0.128 / 0.945 | 0.103 / 0.91 | 0.228 / 0.988 | 0.113 / 0.923 | 0.172 / 0.973 | - / - |

**Table S9: Frequencies of trans-eQTL aSNPs in present-day populations**

| | 1000 Genomes | | | | | | | | | | | | | | | SGDP |
|---|---|---|---|---|---|---|---|---|---|---|---|---|---|---|---|---|
| | FIN | IBS | CEU | GBR | TSI | CHS | CHB | CDX | JPT | KHV | BEB | STU | GIH | ITU | PJL | PAP |
| 11_100453046 | 0.051 / 0.647 | 0.131 / 0.904 | 0.131 / 0.896 | 0.104 / 0.844 | 0.107 / 0.873 | 0.371 / 0.989 | 0.277 / 0.97 | 0.376 / 0.99 | 0.13 / 0.823 | 0.268 / 0.973 | 0.099 / 0.901 | 0.132 / 0.949 | 0.073 / 0.813 | 0.118 / 0.93 | 0.089 / 0.876 | 0.133 / 0.56 |
| 12_56740682 | 0.071 / 0.745 | 0.042 / 0.625 | 0.076 / 0.772 | 0.055 / 0.665 | 0.089 / 0.827 | 0.024 / 0.387 | 0.029 / 0.447 | 0.027 / 0.436 | 0.058 / 0.582 | 0.04 / 0.567 | 0.023 / 0.501 | 0.025 / 0.509 | 0.024 / 0.488 | 0.01 / 0.275 | 0.016 / 0.384 | 0.567 / 0.967 |
| 12_56750204 | 0.071 / 0.745 | 0.042 / 0.625 | 0.076 / 0.772 | 0.055 / 0.665 | 0.079 / 0.799 | 0.024 / 0.387 | 0.024 / 0.405 | 0.032 / 0.479 | 0.053 / 0.555 | 0.04 / 0.567 | 0.023 / 0.501 | 0.025 / 0.509 | 0.024 / 0.488 | 0.01 / 0.275 | 0.016 / 0.384 | 0.567 / 0.967 |
| 15_85207825 | 0.237 / 0.974 | 0.299 / 0.99 | 0.278 / 0.984 | 0.231 / 0.971 | 0.271 / 0.984 | 0.162 / 0.882 | 0.189 / 0.918 | 0.124 / 0.82 | 0.096 / 0.736 | 0.116 / 0.829 | 0.163 / 0.971 | 0.137 / 0.954 | 0.146 / 0.954 | 0.181 / 0.976 | 0.151 / 0.962 | - / - |
| 17_76259847 | 0.242 / 0.976 | 0.304 / 0.991 | 0.242 / 0.975 | 0.297 / 0.988 | 0.299 / 0.99 | 0.029 / 0.425 | 0.063 / 0.639 | 0.027 / 0.436 | 0.062 / 0.604 | 0.035 / 0.538 | 0.174 / 0.977 | 0.113 / 0.926 | 0.165 / 0.966 | 0.127 / 0.942 | 0.172 / 0.973 | - / - |
| 17_800593 | 0.035 / 0.552 | - / - | 0.01 / 0.251 | 0.005 / 0.146 | 0.005 / 0.154 | 0.233 / 0.95 | 0.238 / 0.953 | 0.059 / 0.623 | 0.317 / 0.978 | 0.131 / 0.856 | 0.041 / 0.67 | - / - | 0.015 / 0.354 | 0.005 / 0.163 | 0.016 / 0.384 | - / - |
| 18_74072245 | 0.051 / 0.647 | 0.07 / 0.77 | 0.071 / 0.751 | 0.06 / 0.694 | 0.079 / 0.799 | - / - | 0.005 / 0.17 | - / - | - / - | - / - | 0.035 / 0.621 | 0.025 / 0.509 | 0.039 / 0.631 | 0.044 / 0.685 | 0.052 / 0.74 | - / - |
| 19_33453659 | 0.106 / 0.84 | 0.154 / 0.927 | 0.111 / 0.856 | 0.159 / 0.921 | 0.192 / 0.955 | 0.076 / 0.666 | 0.097 / 0.748 | 0.118 / 0.808 | 0.058 / 0.556 | 0.157 / 0.888 | 0.035 / 0.622 | 0.044 / 0.683 | 0.029 / 0.49 | 0.034 / 0.609 | 0.068 / 0.794 | - / - |
| 1_169099037 | 0.086 / 0.8 | 0.182 / 0.95 | 0.131 / 0.896 | 0.104 / 0.844 | 0.136 / 0.912 | 0.019 / 0.334 | 0.005 / 0.17 | 0.048 / 0.572 | 0.048 / 0.528 | 0.01 / 0.293 | 0.099 / 0.901 | 0.078 / 0.848 | 0.068 / 0.795 | 0.074 / 0.831 | 0.125 / 0.939 | - / - |
| 1_169099483 | 0.086 / 0.8 | 0.182 / 0.95 | 0.131 / 0.896 | 0.104 / 0.844 | 0.14 / 0.917 | 0.019 / 0.334 | 0.005 / 0.17 | 0.048 / 0.572 | 0.048 / 0.528 | 0.01 / 0.293 | 0.099 / 0.901 | 0.078 / 0.848 | 0.068 / 0.795 | 0.074 / 0.831 | 0.125 / 0.939 | - / - |
| 1_22587728 | 0.091 / 0.816 | 0.051 / 0.683 | 0.106 / 0.855 | 0.082 / 0.784 | 0.093 / 0.84 | - / - | 0.005 / 0.17 | - / - | - / - | - / - | 0.023 / 0.501 | 0.025 / 0.509 | 0.039 / 0.631 | 0.01 / 0.275 | 0.036 / 0.628 | - / - |
| 20_16054982 | 0.061 / 0.7 | 0.042 / 0.625 | 0.066 / 0.728 | 0.071 / 0.744 | 0.098 / 0.853 | 0.095 / 0.751 | 0.083 / 0.715 | 0.161 / 0.884 | 0.062 / 0.604 | 0.182 / 0.92 | 0.047 / 0.709 | 0.034 / 0.606 | 0.068 / 0.795 | 0.059 / 0.767 | 0.047 / 0.706 | - / - |
| 20_4132364 | 0.419 / 0.997 | 0.234 / 0.974 | 0.323 / 0.992 | 0.335 / 0.992 | 0.229 / 0.973 | 0.314 / 0.979 | 0.325 / 0.982 | 0.344 / 0.986 | 0.274 / 0.967 | 0.253 / 0.967 | 0.116 / 0.931 | 0.088 / 0.877 | 0.083 / 0.849 | 0.078 / 0.848 | 0.141 / 0.955 | - / - |
| 3_46173072 | 0.187 / 0.949 | 0.084 / 0.815 | 0.121 / 0.88 | 0.11 / 0.858 | 0.121 / 0.897 | 0.038 / 0.493 | 0.092 / 0.748 | 0.032 / 0.479 | 0.053 / 0.555 | 0.03 / 0.508 | 0.355 / 0.999 | 0.343 / 0.998 | 0.374 / 0.999 | 0.373 / 0.999 | 0.38 / 0.999 | - / - |
| 3_46235201 | 0.141 / 0.908 | 0.061 / 0.731 | 0.101 / 0.845 | 0.099 / 0.833 | 0.075 / 0.783 | 0.043 / 0.523 | 0.073 / 0.679 | 0.027 / 0.436 | 0.048 / 0.528 | 0.03 / 0.508 | 0.209 / 0.988 | 0.284 / 0.997 | 0.325 / 0.998 | 0.275 / 0.995 | 0.359 / 0.999 | - / - |
| 3_52506426 | 0.051 / 0.647 | 0.084 / 0.803 | 0.045 / 0.621 | 0.049 / 0.63 | 0.084 / 0.799 | 0.014 / 0.283 | 0.019 / 0.312 | 0.032 / 0.438 | 0.019 / 0.309 | 0.025 / 0.472 | 0.029 / 0.567 | 0.049 / 0.715 | 0.063 / 0.775 | 0.039 / 0.651 | 0.042 / 0.668 | 0.071 / 0.384 |
| 6_52281072 | 0.015 / 0.329 | 0.098 / 0.85 | 0.081 / 0.789 | 0.071 / 0.744 | 0.079 / 0.799 | 0.019 / 0.334 | 0.029 / 0.447 | 0.032 / 0.479 | 0.024 / 0.355 | 0.02 / 0.427 | 0.134 / 0.951 | 0.113 / 0.926 | 0.184 / 0.975 | 0.093 / 0.888 | 0.083 / 0.863 | - / - |
| 7_46402269 | 0.061 / 0.7 | 0.098 / 0.85 | 0.035 / 0.545 | 0.104 / 0.844 | 0.117 / 0.89 | 0.033 / 0.46 | 0.034 / 0.484 | 0.022 / 0.39 | 0.034 / 0.436 | 0.015 / 0.369 | 0.116 / 0.931 | 0.103 / 0.91 | 0.107 / 0.906 | 0.074 / 0.831 | 0.13 / 0.945 | - / - |

**Table S10: CLUES derived allele frequency estimates over last 500 generations and respective summary statistics for a-cTF aSNPs**

| | | | | | | generations | | | | | | | | | | | | | | | | | | | | | | | | | | | | | | | | | | | | | | | | | | | | | | | | | |
|---|---|---|---|---|---|---|---|---|---|---|---|---|---|---|---|---|---|---|---|---|---|---|---|---|---|---|---|---|---|---|---|---|---|---|---|---|---|---|---|---|---|---|---|---|---|---|---|---|---|---|---|---|---|---|---|
| chr | pos | rs | proxy_for | logLR | s | g0 | g10 | g20 | g30 | g40 | g50 | g60 | g70 | g80 | g90 | g100 | g110 | g120 | g130 | g140 | g150 | g160 | g170 | g180 | g190 | g200 | g210 | g220 | g230 | g240 | g250 | g260 | g270 | g280 | g290 | g300 | g310 | g320 | g330 | g340 | g350 | g360 | g370 | g380 | g390 | g400 | g410 | g420 | g430 | g440 | g450 | g460 | g470 | g480 | g490 |
| 1 | 58900316 | rs77554680 | na | 0.5465 | 0.00151 | 0.185 | 0.185 | 0.1849 | 0.1859 | 0.189 | 0.1934 | 0.1958 | 0.197 | 0.1992 | 0.2015 | 0.2035 | 0.2038 | 0.2029 | 0.2018 | 0.2019 | 0.2023 | 0.2026 | 0.202 | 0.2012 | 0.2004 | 0.1996 | 0.1993 | 0.199 | 0.1981 | 0.1974 | 0.1963 | 0.195 | 0.1929 | 0.1904 | 0.1878 | 0.1854 | 0.1838 | 0.1825 | 0.1817 | 0.1811 | 0.1803 | 0.1795 | 0.1783 | 0.177 | 0.1756 | 0.1743 | 0.173 | 0.1717 | 0.1706 | 0.17 | 0.1692 | 0.1681 | 0.1667 | 0.1653 | 0.1638 |
| 1 | 58900316 | rs77554680 | na | 0.0703 | 0.00056 | 0.1775 | 0.1775 | 0.1778 | 0.1774 | 0.1784 | 0.181 | 0.1848 | 0.1879 | 0.1906 | 0.1925 | 0.1938 | 0.1948 | 0.1954 | 0.1962 | 0.1967 | 0.1962 | 0.1964 | 0.1969 | 0.1969 | 0.1967 | 0.1964 | 0.1958 | 0.1947 | 0.1931 | 0.1911 | 0.1883 | 0.1857 | 0.1837 | 0.1817 | 0.179 | 0.176 | 0.1733 | 0.1703 | 0.1677 | 0.1648 | 0.1617 | 0.1586 | 0.1556 | 0.1529 | 0.1505 | 0.1487 | 0.1468 | 0.145 | 0.1433 | 0.142 | 0.1409 | 0.1396 | 0.1384 | 0.1372 | 0.1362 |
| 1 | 165316371 | rs12748775 | na | 0.8635 | 0.00232 | 0.1316 | 0.1316 | 0.1313 | 0.1309 | 0.1308 | 0.1304 | 0.1287 | 0.1264 | 0.1255 | 0.1261 | 0.1262 | 0.1252 | 0.1233 | 0.1207 | 0.1184 | 0.116 | 0.1146 | 0.1138 | 0.1136 | 0.1133 | 0.1128 | 0.1119 | 0.111 | 0.1105 | 0.11 | 0.1093 | 0.1087 | 0.108 | 0.1068 | 0.1057 | 0.1048 | 0.1044 | 0.104 | 0.1034 | 0.103 | 0.1023 | 0.1012 | 0.1 | 0.0987 | 0.0981 | 0.0977 | 0.0972 | 0.0965 | 0.0956 | 0.0946 | 0.0934 | 0.092 | 0.0904 | 0.0885 | 0.0864 |
| 1 | 165316371 | rs12748775 | na | 2.1736 | 0.00391 | 0.1312 | 0.1312 | 0.1308 | 0.1305 | 0.1299 | 0.1291 | 0.1282 | 0.1276 | 0.1263 | 0.1248 | 0.1249 | 0.1251 | 0.1255 | 0.1257 | 0.125 | 0.1239 | 0.1228 | 0.1213 | 0.1198 | 0.1175 | 0.1152 | 0.1128 | 0.1105 | 0.1086 | 0.1062 | 0.1037 | 0.1014 | 0.0996 | 0.098 | 0.0961 | 0.0939 | 0.0922 | 0.0905 | 0.0891 | 0.0881 | 0.0871 | 0.0863 | 0.0852 | 0.084 | 0.0826 | 0.0809 | 0.0793 | 0.0777 | 0.0762 | 0.0746 | 0.073 | 0.0717 | 0.0702 | 0.0685 | 0.0667 |
| 1 | 170634895 | rs77401317 | na | 5.2013 | 0.00703 | 0.1254 | 0.1254 | 0.1242 | 0.1223 | 0.1192 | 0.1155 | 0.1101 | 0.1045 | 0.1008 | 0.0991 | 0.0974 | 0.0946 | 0.0912 | 0.0887 | 0.0851 | 0.0816 | 0.0788 | 0.0775 | 0.0758 | 0.0741 | 0.0722 | 0.0698 | 0.0671 | 0.0645 | 0.062 | 0.0592 | 0.0563 | 0.0528 | 0.0498 | 0.0473 | 0.0452 | 0.0432 | 0.0411 | 0.0393 | 0.0376 | 0.036 | 0.0346 | 0.0332 | 0.0318 | 0.0302 | 0.0286 | 0.0275 | 0.0265 | 0.0254 | 0.0243 | 0.023 | 0.0216 | 0.0205 | 0.0193 | 0.0181 |
| 1 | 170634895 | rs77401317 | na | 4.0468 | 0.00601 | 0.1361 | 0.1361 | 0.135 | 0.1329 | 0.1298 | 0.1283 | 0.1282 | 0.1283 | 0.1265 | 0.1242 | 0.1223 | 0.1206 | 0.1182 | 0.1171 | 0.1168 | 0.1166 | 0.1156 | 0.1147 | 0.1136 | 0.1129 | 0.112 | 0.1111 | 0.1098 | 0.1084 | 0.1063 | 0.104 | 0.1015 | 0.0994 | 0.0971 | 0.0948 | 0.0927 | 0.0908 | 0.089 | 0.0872 | 0.085 | 0.0828 | 0.0808 | 0.0789 | 0.0769 | 0.0748 | 0.0726 | 0.0709 | 0.0688 | 0.0668 | 0.0648 | 0.0627 | 0.0603 | 0.0583 | 0.0562 | 0.0539 |
| 1 | 205456751 | rs77115904 | na | 0.1566 | 0.00132 | 0.0702 | 0.0702 | 0.07 | 0.0702 | 0.0718 | 0.072 | 0.0722 | 0.0721 | 0.072 | 0.0723 | 0.0727 | 0.0728 | 0.0727 | 0.0734 | 0.0746 | 0.0754 | 0.0758 | 0.0758 | 0.0758 | 0.076 | 0.0763 | 0.0766 | 0.0771 | 0.0775 | 0.0777 | 0.0776 | 0.0773 | 0.0771 | 0.0771 | 0.0768 | 0.0763 | 0.0756 | 0.0746 | 0.0735 | 0.0729 | 0.0723 | 0.0718 | 0.0712 | 0.0705 | 0.0698 | 0.0693 | 0.0688 | 0.0681 | 0.0674 | 0.0665 | 0.0659 | 0.0651 | 0.0642 | 0.0632 | 0.0621 |
| 1 | 205456751 | rs77115904 | na | 0.5498 | 0.00244 | 0.0773 | 0.0773 | 0.0773 | 0.0767 | 0.0754 | 0.0729 | 0.0695 | 0.0662 | 0.0634 | 0.0623 | 0.0616 | 0.0611 | 0.0599 | 0.059 | 0.0584 | 0.0578 | 0.0573 | 0.0564 | 0.0556 | 0.055 | 0.0546 | 0.054 | 0.0534 | 0.0525 | 0.0513 | 0.0498 | 0.0483 | 0.0469 | 0.0454 | 0.0445 | 0.0438 | 0.0434 | 0.0429 | 0.0422 | 0.0415 | 0.0406 | 0.0396 | 0.0385 | 0.0372 | 0.0358 | 0.0344 | 0.0331 | 0.0316 | 0.0307 | 0.0298 | 0.0289 | 0.0278 | 0.0271 | 0.0263 | 0.0255 |
| 2 | 68475595 | rs4671887 | na | 3.8693 | 0.00339 | 0.3612 | 0.3612 | 0.3608 | 0.3595 | 0.3573 | 0.353 | 0.35 | 0.3486 | 0.3484 | 0.3482 | 0.3481 | 0.3469 | 0.3463 | 0.3446 | 0.3426 | 0.3405 | 0.3377 | 0.3345 | 0.3312 | 0.3279 | 0.3247 | 0.321 | 0.3167 | 0.3126 | 0.3091 | 0.3055 | 0.3018 | 0.2978 | 0.2943 | 0.2915 | 0.2887 | 0.286 | 0.2832 | 0.2805 | 0.2783 | 0.2759 | 0.2739 | 0.2721 | 0.27 | 0.2678 | 0.2653 | 0.2624 | 0.2595 | 0.2565 | 0.2535 | 0.2503 | 0.2467 | 0.243 | 0.2392 | 0.235 |
| 2 | 68475595 | rs4671887 | na | 2.1699 | 0.00271 | 0.3589 | 0.3589 | 0.3583 | 0.3567 | 0.3548 | 0.351 | 0.3513 | 0.3521 | 0.353 | 0.3531 | 0.3508 | 0.3481 | 0.3466 | 0.3456 | 0.3435 | 0.3411 | 0.3388 | 0.3363 | 0.3334 | 0.331 | 0.328 | 0.3239 | 0.3189 | 0.3136 | 0.3084 | 0.3029 | 0.2977 | 0.2931 | 0.2885 | 0.2835 | 0.2787 | 0.2744 | 0.2704 | 0.2661 | 0.2622 | 0.2595 | 0.2571 | 0.2546 | 0.2525 | 0.2501 | 0.248 | 0.2461 | 0.244 | 0.2421 | 0.2399 | 0.2377 | 0.2351 | 0.2323 | 0.2292 | 0.2257 |
| 2 | 86623524 | rs75988363 | na | 0.0096 | 0.00039 | 0.0542 | 0.0542 | 0.054 | 0.0528 | 0.0508 | 0.0488 | 0.0476 | 0.0458 | 0.0438 | 0.0427 | 0.042 | 0.0413 | 0.0403 | 0.0394 | 0.0382 | 0.0375 | 0.0365 | 0.0357 | 0.0349 | 0.0348 | 0.0353 | 0.0359 | 0.0364 | 0.0369 | 0.0372 | 0.0374 | 0.0375 | 0.0378 | 0.038 | 0.0382 | 0.0383 | 0.0383 | 0.0383 | 0.0384 | 0.0385 | 0.0385 | 0.0385 | 0.0384 | 0.0382 | 0.038 | 0.0378 | 0.0378 | 0.0377 | 0.0377 | 0.0375 | 0.0374 | 0.0372 | 0.0369 | 0.0366 | 0.0362 |
| 2 | 86623524 | rs75988363 | na | 0.2575 | 0.00205 | 0.0556 | 0.0556 | 0.0554 | 0.0555 | 0.0552 | 0.0538 | 0.0516 | 0.0495 | 0.0481 | 0.0473 | 0.0466 | 0.0458 | 0.0453 | 0.0448 | 0.0443 | 0.0447 | 0.045 | 0.045 | 0.0449 | 0.0449 | 0.0447 | 0.0447 | 0.0448 | 0.0447 | 0.0444 | 0.0441 | 0.0437 | 0.043 | 0.0425 | 0.0421 | 0.0418 | 0.0417 | 0.0414 | 0.0411 | 0.0407 | 0.0401 | 0.0394 | 0.0386 | 0.0379 | 0.0374 | 0.0369 | 0.0364 | 0.0358 | 0.0352 | 0.0344 | 0.0335 | 0.033 | 0.0324 | 0.0317 | 0.0309 |
| 2 | 238648521 | rs3769071 | na | 0.0003 | 5E-05 | 0.0513 | 0.0513 | 0.0508 | 0.0501 | 0.0496 | 0.0509 | 0.0525 | 0.055 | 0.0564 | 0.058 | 0.059 | 0.0594 | 0.0602 | 0.0604 | 0.0605 | 0.0606 | 0.0612 | 0.0619 | 0.0625 | 0.0629 | 0.063 | 0.0629 | 0.0633 | 0.0636 | 0.0644 | 0.0652 | 0.0657 | 0.0662 | 0.0668 | 0.0673 | 0.0676 | 0.0678 | 0.068 | 0.0681 | 0.0686 | 0.0689 | 0.0693 | 0.0695 | 0.0695 | 0.0697 | 0.0699 | 0.07 | 0.07 | 0.0699 | 0.0697 | 0.0694 | 0.0693 | 0.069 | 0.0687 | 0.0683 |
| 2 | 238648521 | rs3769071 | na | 0.1073 | -0.00112 | 0.0497 | 0.0497 | 0.0495 | 0.048 | 0.0467 | 0.0463 | 0.0465 | 0.0475 | 0.0483 | 0.0491 | 0.0501 | 0.0504 | 0.0507 | 0.0513 | 0.0518 | 0.0518 | 0.0514 | 0.0512 | 0.0505 | 0.0508 | 0.0513 | 0.0518 | 0.0522 | 0.0525 | 0.0525 | 0.0525 | 0.0526 | 0.0528 | 0.053 | 0.0539 | 0.0548 | 0.0556 | 0.0563 | 0.057 | 0.0577 | 0.0584 | 0.059 | 0.0595 | 0.0601 | 0.0606 | 0.061 | 0.0613 | 0.0616 | 0.0617 | 0.0618 | 0.0618 | 0.0617 | 0.0619 | 0.062 | 0.062 |
| 3 | 20041386 | rs62241326 | 20052020 | 0.9059 | 0.00188 | 0.2227 | 0.2227 | 0.2227 | 0.2223 | 0.2201 | 0.2198 | 0.2206 | 0.2205 | 0.2197 | 0.2199 | 0.2194 | 0.2187 | 0.2173 | 0.2149 | 0.2124 | 0.2116 | 0.2118 | 0.2121 | 0.2114 | 0.211 | 0.2099 | 0.2086 | 0.2067 | 0.2032 | 0.1998 | 0.1962 | 0.1925 | 0.1886 | 0.1853 | 0.182 | 0.1787 | 0.1757 | 0.1728 | 0.1699 | 0.1672 | 0.1643 | 0.1615 | 0.1585 | 0.1555 | 0.1525 | 0.1494 | 0.1465 | 0.1436 | 0.1408 | 0.1382 | 0.1355 | 0.1332 | 0.1309 | 0.1285 | 0.1262 |
| 3 | 20041386 | rs62241326 | 20052020 | 0.9256 | 0.00195 | 0.2228 | 0.2228 | 0.2229 | 0.2238 | 0.2261 | 0.2256 | 0.2239 | 0.2207 | 0.2201 | 0.2202 | 0.2189 | 0.2173 | 0.2166 | 0.2162 | 0.2163 | 0.216 | 0.2142 | 0.2116 | 0.2092 | 0.2069 | 0.2045 | 0.2025 | 0.2001 | 0.1974 | 0.1934 | 0.1899 | 0.187 | 0.1847 | 0.1827 | 0.1806 | 0.1783 | 0.1761 | 0.1742 | 0.1727 | 0.1707 | 0.1684 | 0.1659 | 0.1632 | 0.1611 | 0.1594 | 0.1578 | 0.1561 | 0.1544 | 0.1527 | 0.1508 | 0.1487 | 0.1465 | 0.1441 | 0.1422 | 0.1401 |
| 3 | 20104838 | rs13066876 | na | 0.1021 | 0.00059 | 0.2422 | 0.2422 | 0.243 | 0.2476 | 0.2559 | 0.2616 | 0.2662 | 0.2698 | 0.2715 | 0.2694 | 0.2668 | 0.2635 | 0.2597 | 0.2568 | 0.2541 | 0.2524 | 0.2505 | 0.2486 | 0.2466 | 0.2442 | 0.2423 | 0.2407 | 0.2389 | 0.2377 | 0.2364 | 0.2353 | 0.2342 | 0.233 | 0.2315 | 0.2299 | 0.228 | 0.2257 | 0.2242 | 0.2231 | 0.2221 | 0.2205 | 0.219 | 0.2173 | 0.2159 | 0.2147 | 0.2135 | 0.2125 | 0.2111 | 0.2097 | 0.2083 | 0.207 | 0.2058 | 0.2045 | 0.2036 | 0.2028 |
| 3 | 20104838 | rs13066876 | na | 0.0099 | 0.00017 | 0.2381 | 0.2381 | 0.2385 | 0.2415 | 0.2452 | 0.2482 | 0.2514 | 0.2515 | 0.2515 | 0.2533 | 0.2548 | 0.2551 | 0.255 | 0.2547 | 0.2543 | 0.2547 | 0.2544 | 0.254 | 0.253 | 0.2521 | 0.2505 | 0.2485 | 0.2465 | 0.2444 | 0.2417 | 0.2387 | 0.2362 | 0.2336 | 0.2308 | 0.2282 | 0.2256 | 0.2225 | 0.22 | 0.2183 | 0.2167 | 0.2153 | 0.2138 | 0.2124 | 0.2109 | 0.2095 | 0.2084 | 0.2075 | 0.2069 | 0.2064 | 0.2059 | 0.2055 | 0.2051 | 0.2046 | 0.2042 | 0.204 |
| 3 | 141891385 | rs73236075 | na | 0.2166 | 0.00125 | 0.1084 | 0.1084 | 0.1084 | 0.1091 | 0.1113 | 0.1133 | 0.1139 | 0.1146 | 0.1139 | 0.1137 | 0.1124 | 0.11 | 0.1063 | 0.103 | 0.1006 | 0.0994 | 0.0979 | 0.0964 | 0.0951 | 0.0941 | 0.0939 | 0.0938 | 0.0939 | 0.094 | 0.0942 | 0.0941 | 0.0939 | 0.0935 | 0.0931 | 0.093 | 0.0931 | 0.093 | 0.0929 | 0.093 | 0.0929 | 0.0927 | 0.0922 | 0.0915 | 0.0907 | 0.0898 | 0.0888 | 0.0877 | 0.0863 | 0.085 | 0.0837 | 0.0825 | 0.0815 | 0.0804 | 0.0791 | 0.078 |
| 3 | 141891385 | rs73236075 | na | 0.8667 | 0.00234 | 0.1133 | 0.1133 | 0.1135 | 0.1151 | 0.1181 | 0.1218 | 0.1244 | 0.1258 | 0.1257 | 0.1259 | 0.1236 | 0.1208 | 0.1191 | 0.1165 | 0.1141 | 0.1125 | 0.1107 | 0.1095 | 0.1081 | 0.1064 | 0.1043 | 0.1017 | 0.099 | 0.0966 | 0.0947 | 0.0927 | 0.0907 | 0.0887 | 0.0867 | 0.0849 | 0.0827 | 0.0804 | 0.0783 | 0.0757 | 0.0729 | 0.0696 | 0.0663 | 0.0632 | 0.0604 | 0.0576 | 0.0549 | 0.0522 | 0.0493 | 0.047 | 0.0453 | 0.0441 | 0.0432 | 0.0423 | 0.0413 | 0.0402 |
| 4 | 121643726 | rs13110077 | na | 4.6381 | 0.00452 | 0.2442 | 0.2442 | 0.2441 | 0.2447 | 0.2413 | 0.2306 | 0.2227 | 0.2151 | 0.2095 | 0.2041 | 0.199 | 0.1937 | 0.188 | 0.1828 | 0.1785 | 0.1748 | 0.1721 | 0.1704 | 0.1697 | 0.1682 | 0.1664 | 0.1644 | 0.1631 | 0.1618 | 0.1602 | 0.1586 | 0.1573 | 0.1563 | 0.1547 | 0.1533 | 0.1524 | 0.1515 | 0.1506 | 0.1494 | 0.148 | 0.1466 | 0.1454 | 0.144 | 0.1425 | 0.1414 | 0.1403 | 0.1392 | 0.1379 | 0.1364 | 0.1349 | 0.1331 | 0.1309 | 0.1282 | 0.1249 | 0.1213 |
| 4 | 121643726 | rs13110077 | na | 2.7127 | 0.00339 | 0.2405 | 0.2405 | 0.2406 | 0.2415 | 0.2407 | 0.2382 | 0.236 | 0.2337 | 0.2308 | 0.2279 | 0.2236 | 0.219 | 0.2142 | 0.2097 | 0.2046 | 0.1995 | 0.1949 | 0.191 | 0.1866 | 0.1824 | 0.179 | 0.1753 | 0.1716 | 0.1681 | 0.1652 | 0.1622 | 0.1585 | 0.1547 | 0.1507 | 0.147 | 0.1439 | 0.1412 | 0.1391 | 0.1372 | 0.1352 | 0.1334 | 0.1314 | 0.1291 | 0.1273 | 0.1254 | 0.1239 | 0.1222 | 0.1208 | 0.1194 | 0.118 | 0.1164 | 0.1146 | 0.1127 | 0.1106 | 0.1083 |
| 4 | 121703570 | rs75558997 | na | 2.3013 | 0.00273 | 0.3213 | 0.3213 | 0.3211 | 0.32 | 0.3173 | 0.3106 | 0.303 | 0.2954 | 0.2886 | 0.2786 | 0.2686 | 0.2611 | 0.2558 | 0.2504 | 0.2454 | 0.2411 | 0.2364 | 0.2331 | 0.2298 | 0.2265 | 0.2238 | 0.2213 | 0.2193 | 0.2176 | 0.216 | 0.214 | 0.2115 | 0.2092 | 0.2075 | 0.206 | 0.2046 | 0.2034 | 0.2023 | 0.2015 | 0.2007 | 0.1997 | 0.1985 | 0.197 | 0.1952 | 0.1931 | 0.191 | 0.1893 | 0.188 | 0.1865 | 0.1851 | 0.1835 | 0.1814 | 0.179 | 0.1764 | 0.1736 |
| 4 | 121703570 | rs75558997 | na | 1.4786 | 0.0022 | 0.324 | 0.324 | 0.324 | 0.3244 | 0.3235 | 0.3195 | 0.3143 | 0.3101 | 0.3041 | 0.2966 | 0.2893 | 0.282 | 0.2765 | 0.2728 | 0.2712 | 0.2704 | 0.2687 | 0.2674 | 0.2652 | 0.2642 | 0.2642 | 0.2644 | 0.2641 | 0.2633 | 0.263 | 0.2619 | 0.2595 | 0.2569 | 0.2544 | 0.2525 | 0.2511 | 0.25 | 0.2491 | 0.2478 | 0.2466 | 0.2454 | 0.2439 | 0.2421 | 0.2401 | 0.2379 | 0.2357 | 0.2339 | 0.2319 | 0.2297 | 0.2274 | 0.2248 | 0.2218 | 0.2192 | 0.2166 | 0.214 |
| 6 | 1364471 | rs1922929 | na | 0.0073 | 0.00024 | 0.9021 | 0.9021 | 0.9024 | 0.9035 | 0.906 | 0.9082 | 0.9089 | 0.9086 | 0.9081 | 0.9081 | 0.9078 | 0.908 | 0.9078 | 0.9074 | 0.9072 | 0.907 | 0.907 | 0.9071 | 0.9075 | 0.9083 | 0.9087 | 0.9088 | 0.909 | 0.9091 | 0.909 | 0.9088 | 0.9084 | 0.9082 | 0.9083 | 0.9085 | 0.908 | 0.9071 | 0.9063 | 0.9055 | 0.9046 | 0.9037 | 0.9029 | 0.9023 | 0.9017 | 0.9011 | 0.9007 | 0.9003 | 0.9001 | 0.8999 | 0.8999 | 0.8996 | 0.8995 | 0.8995 | 0.8997 | 0.8998 |
| 6 | 1364471 | rs1922929 | na | 0.0069 | -0.0002 | 0.9016 | 0.9016 | 0.9017 | 0.9021 | 0.9032 | 0.9032 | 0.9027 | 0.902 | 0.9023 | 0.9031 | 0.9053 | 0.9077 | 0.9086 | 0.9082 | 0.9077 | 0.9072 | 0.907 | 0.9064 | 0.9051 | 0.9031 | 0.9008 | 0.8984 | 0.8963 | 0.8947 | 0.8935 | 0.8921 | 0.8906 | 0.8891 | 0.8878 | 0.8867 | 0.8858 | 0.8852 | 0.885 | 0.8851 | 0.8849 | 0.8847 | 0.8846 | 0.8845 | 0.8847 | 0.8848 | 0.8849 | 0.8851 | 0.8856 | 0.8863 | 0.8869 | 0.8873 | 0.8873 | 0.8874 | 0.8877 | 0.8878 |
| 6 | 1623627 | rs62390617 | na | 0.014 | 0.00029 | 0.1129 | 0.1129 | 0.1129 | 0.1141 | 0.1163 | 0.1187 | 0.1217 | 0.1229 | 0.1242 | 0.125 | 0.126 | 0.1264 | 0.1263 | 0.1253 | 0.1254 | 0.1253 | 0.1247 | 0.1238 | 0.1225 | 0.1207 | 0.1188 | 0.1174 | 0.1167 | 0.1162 | 0.1153 | 0.114 | 0.1129 | 0.1118 | 0.111 | 0.1099 | 0.1085 | 0.1068 | 0.105 | 0.104 | 0.1028 | 0.1013 | 0.1001 | 0.0992 | 0.0983 | 0.0971 | 0.0956 | 0.0939 | 0.0924 | 0.091 | 0.0902 | 0.0892 | 0.0879 | 0.0867 | 0.0855 | 0.0844 |
| 6 | 1623627 | rs62390617 | na | 0.112 | 0.00088 | 0.1143 | 0.1143 | 0.1143 | 0.1134 | 0.1117 | 0.1131 | 0.1161 | 0.1213 | 0.1253 | 0.1285 | 0.1315 | 0.1329 | 0.133 | 0.134 | 0.1351 | 0.1365 | 0.1378 | 0.1386 | 0.1386 | 0.1382 | 0.1375 | 0.137 | 0.1366 | 0.1357 | 0.135 | 0.134 | 0.1328 | 0.1315 | 0.1301 | 0.1281 | 0.1256 | 0.1234 | 0.1214 | 0.1196 | 0.1183 | 0.1168 | 0.1154 | 0.1139 | 0.1126 | 0.1116 | 0.1108 | 0.1101 | 0.1092 | 0.108 | 0.1067 | 0.1054 | 0.1039 | 0.1023 | 0.1008 | 0.0995 |
| 6 | 18921942 | rs72826453 | na | 0.0312 | 0.00068 | 0.0533 | 0.0533 | 0.0533 | 0.0537 | 0.0539 | 0.0548 | 0.0543 | 0.0544 | 0.055 | 0.055 | 0.0559 | 0.057 | 0.0579 | 0.0592 | 0.0603 | 0.061 | 0.061 | 0.0615 | 0.0621 | 0.0628 | 0.0635 | 0.064 | 0.0643 | 0.0641 | 0.0638 | 0.0634 | 0.063 | 0.0625 | 0.0622 | 0.0616 | 0.0611 | 0.0608 | 0.0604 | 0.0604 | 0.0606 | 0.0608 | 0.0609 | 0.061 | 0.0608 | 0.0605 | 0.0603 | 0.0603 | 0.0601 | 0.0599 | 0.0594 | 0.059 | 0.0588 | 0.0584 | 0.058 | 0.0574 |
| 6 | 18921942 | rs72826453 | na | 0.2691 | 0.0021 | 0.0554 | 0.0554 | 0.0554 | 0.0557 | 0.0571 | 0.0596 | 0.0612 | 0.0622 | 0.0631 | 0.0635 | 0.0629 | 0.0617 | 0.0607 | 0.0597 | 0.0584 | 0.0572 | 0.0563 | 0.0552 | 0.0543 | 0.0527 | 0.051 | 0.0496 | 0.0478 | 0.0457 | 0.0441 | 0.043 | 0.0421 | 0.0411 | 0.0402 | 0.0395 | 0.0388 | 0.0379 | 0.037 | 0.0365 | 0.0361 | 0.0359 | 0.0356 | 0.0353 | 0.0349 | 0.0345 | 0.0339 | 0.0337 | 0.0334 | 0.033 | 0.0326 | 0.0321 | 0.0316 | 0.0311 | 0.0304 | 0.0296 |
| 6 | 27361242 | rs12212348 | na | 0.2728 | 0.00166 | 0.0685 | 0.0685 | 0.0681 | 0.0673 | 0.0661 | 0.0643 | 0.0617 | 0.0604 | 0.0599 | 0.0607 | 0.0605 | 0.0608 | 0.0617 | 0.0628 | 0.0641 | 0.0654 | 0.0666 | 0.0671 | 0.0677 | 0.0681 | 0.0686 | 0.069 | 0.069 | 0.0688 | 0.0681 | 0.0668 | 0.0648 | 0.0628 | 0.0608 | 0.0588 | 0.0568 | 0.0549 | 0.0527 | 0.0505 | 0.0482 | 0.0461 | 0.0444 | 0.043 | 0.0417 | 0.0403 | 0.0392 | 0.038 | 0.0367 | 0.0353 | 0.034 | 0.0331 | 0.0321 | 0.0315 | 0.0308 | 0.0301 |
| 6 | 27361242 | rs12212348 | na | 1.6044 | 0.00483 | 0.0671 | 0.0671 | 0.0663 | 0.0641 | 0.0612 | 0.0607 | 0.0616 | 0.0626 | 0.0651 | 0.0679 | 0.0704 | 0.0723 | 0.0733 | 0.0735 | 0.0737 | 0.0733 | 0.0725 | 0.0712 | 0.0694 | 0.0676 | 0.0652 | 0.063 | 0.0612 | 0.0593 | 0.0575 | 0.0564 | 0.0553 | 0.054 | 0.0524 | 0.0508 | 0.0491 | 0.0471 | 0.0447 | 0.0424 | 0.0402 | 0.0383 | 0.0366 | 0.0353 | 0.034 | 0.0327 | 0.0315 | 0.0306 | 0.0298 | 0.0288 | 0.0279 | 0.0268 | 0.0256 | 0.0243 | 0.0228 | 0.0217 |
| 6 | 41673067 | rs6906513 | na | 0.0009 | -5E-05 | 0.1262 | 0.1262 | 0.1266 | 0.1292 | 0.1336 | 0.1355 | 0.1351 | 0.134 | 0.1353 | 0.1375 | 0.1403 | 0.1429 | 0.1446 | 0.1453 | 0.1467 | 0.1484 | 0.1494 | 0.1496 | 0.1499 | 0.1501 | 0.15 | 0.1498 | 0.1495 | 0.1486 | 0.1478 | 0.1472 | 0.1468 | 0.146 | 0.1451 | 0.1442 | 0.1434 | 0.1427 | 0.1424 | 0.142 | 0.1416 | 0.1412 | 0.1405 | 0.1401 | 0.1395 | 0.1385 | 0.1374 | 0.1363 | 0.1354 | 0.1344 | 0.1336 | 0.1331 | 0.1328 | 0.1327 | 0.1327 | 0.1325 |
| 6 | 41673067 | rs6906513 | na | 0.1039 | -0.00078 | 0.1299 | 0.1299 | 0.1302 | 0.1318 | 0.1346 | 0.137 | 0.1378 | 0.1386 | 0.1391 | 0.1398 | 0.1395 | 0.1389 | 0.1396 | 0.141 | 0.1418 | 0.1429 | 0.1439 | 0.1445 | 0.1453 | 0.1458 | 0.1463 | 0.1473 | 0.1478 | 0.1483 | 0.1485 | 0.1489 | 0.1491 | 0.1499 | 0.151 | 0.1521 | 0.1526 | 0.1533 | 0.1542 | 0.1549 | 0.1553 | 0.1551 | 0.1553 | 0.1552 | 0.1548 | 0.1544 | 0.1537 | 0.1529 | 0.1521 | 0.1512 | 0.1508 | 0.1505 | 0.1501 | 0.1498 | 0.1498 | 0.1498 |
| 6 | 41700858 | rs35088334 | na | 0.2212 | 0.00156 | 0.0747 | 0.0747 | 0.0746 | 0.0746 | 0.0741 | 0.075 | 0.0765 | 0.0782 | 0.0798 | 0.0812 | 0.0819 | 0.0834 | 0.0852 | 0.0867 | 0.0886 | 0.09 | 0.0908 | 0.0911 | 0.0908 | 0.0905 | 0.0908 | 0.0913 | 0.0923 | 0.0932 | 0.0942 | 0.0954 | 0.0965 | 0.0976 | 0.0987 | 0.1 | 0.1012 | 0.1022 | 0.1032 | 0.104 | 0.1046 | 0.105 | 0.1052 | 0.1058 | 0.1064 | 0.1069 | 0.1072 | 0.1074 | 0.1073 | 0.1071 | 0.1067 | 0.1061 | 0.1052 | 0.1042 | 0.1033 | 0.1022 |
| 6 | 41700858 | rs35088334 | na | 0.6205 | 0.00256 | 0.0824 | 0.0824 | 0.0825 | 0.0827 | 0.0839 | 0.0848 | 0.0856 | 0.0864 | 0.0865 | 0.087 | 0.0872 | 0.0882 | 0.0889 | 0.0902 | 0.0914 | 0.0922 | 0.0927 | 0.0926 | 0.093 | 0.0928 | 0.0924 | 0.0917 | 0.0911 | 0.0903 | 0.0895 | 0.0886 | 0.0873 | 0.0862 | 0.0851 | 0.0841 | 0.0827 | 0.0812 | 0.0799 | 0.0784 | 0.077 | 0.0754 | 0.0739 | 0.0721 | 0.0704 | 0.0689 | 0.0676 | 0.0665 | 0.0658 | 0.0651 | 0.0641 | 0.063 | 0.0617 | 0.0602 | 0.0588 | 0.0575 |
| 6 | 43370915 | rs74690967 | na | 0.3166 | 0.00161 | 0.1013 | 0.1013 | 0.1013 | 0.1026 | 0.1049 | 0.1071 | 0.1073 | 0.1055 | 0.1044 | 0.1037 | 0.1038 | 0.1045 | 0.1048 | 0.1045 | 0.105 | 0.1043 | 0.1032 | 0.1023 | 0.1013 | 0.1013 | 0.1017 | 0.1017 | 0.1014 | 0.101 | 0.1008 | 0.1007 | 0.1005 | 0.1006 | 0.1008 | 0.101 | 0.1014 | 0.102 | 0.1022 | 0.1025 | 0.1029 | 0.1031 | 0.1033 | 0.1034 | 0.1033 | 0.1034 | 0.1034 | 0.1032 | 0.1029 | 0.1023 | 0.1016 | 0.1011 | 0.1005 | 0.0996 | 0.0986 | 0.0974 |
| 6 | 43370915 | rs74690967 | na | 0.4065 | 0.00181 | 0.1067 | 0.1067 | 0.1068 | 0.1067 | 0.1054 | 0.1031 | 0.1011 | 0.0993 | 0.0974 | 0.0946 | 0.0918 | 0.0883 | 0.0852 | 0.0824 | 0.0795 | 0.0778 | 0.0759 | 0.0743 | 0.0732 | 0.0721 | 0.0712 | 0.0707 | 0.07 | 0.0695 | 0.0695 | 0.0699 | 0.0704 | 0.0711 | 0.0718 | 0.0723 | 0.0725 | 0.0726 | 0.0727 | 0.0728 | 0.0728 | 0.0727 | 0.0725 | 0.0724 | 0.0721 | 0.0716 | 0.0709 | 0.0701 | 0.069 | 0.0678 | 0.0667 | 0.066 | 0.0656 | 0.065 | 0.0641 | 0.0631 |
| 7 | 27935254 | rs215 | na | 0.0234 | 0.00054 | 0.0511 | 0.0511 | 0.0508 | 0.049 | 0.0463 | 0.0452 | 0.0446 | 0.0445 | 0.0455 | 0.0458 | 0.046 | 0.0464 | 0.0467 | 0.0478 | 0.0487 | 0.0491 | 0.05 | 0.0509 | 0.0514 | 0.0517 | 0.0518 | 0.052 | 0.0525 | 0.0529 | 0.053 | 0.053 | 0.0529 | 0.0528 | 0.0526 | 0.0524 | 0.0521 | 0.0519 | 0.0517 | 0.0517 | 0.0514 | 0.051 | 0.0505 | 0.0501 | 0.0496 | 0.049 | 0.0482 | 0.0478 | 0.0473 | 0.0466 | 0.0459 | 0.045 | 0.0441 | 0.0433 | 0.0423 | 0.0416 |
| 7 | 27935254 | rs215 | na | 0.1382 | 0.00137 | 0.058 | 0.058 | 0.0581 | 0.0585 | 0.0587 | 0.058 | 0.0578 | 0.0571 | 0.0566 | 0.0562 | 0.0551 | 0.0547 | 0.0552 | 0.0554 | 0.0559 | 0.0572 | 0.0583 | 0.0591 | 0.0596 | 0.0598 | 0.0595 | 0.0593 | 0.0592 | 0.059 | 0.0586 | 0.058 | 0.0573 | 0.0563 | 0.056 | 0.0557 | 0.0556 | 0.0556 | 0.0554 | 0.0553 | 0.0551 | 0.0548 | 0.0545 | 0.0543 | 0.0539 | 0.0535 | 0.053 | 0.0523 | 0.0516 | 0.0507 | 0.0498 | 0.049 | 0.0482 | 0.0477 | 0.0471 | 0.0464 |
| 7 | 44790921 | rs55705735 | na | 0.0624 | -0.00063 | 0.1028 | 0.1028 | 0.103 | 0.1045 | 0.1079 | 0.1145 | 0.122 | 0.1284 | 0.1342 | 0.1382 | 0.1407 | 0.1439 | 0.1467 | 0.1499 | 0.153 | 0.1563 | 0.1595 | 0.1626 | 0.1651 | 0.1673 | 0.1696 | 0.1724 | 0.1752 | 0.1779 | 0.18 | 0.1823 | 0.1844 | 0.1861 | 0.1877 | 0.1887 | 0.1894 | 0.19 | 0.1907 | 0.1912 | 0.1915 | 0.1918 | 0.1919 | 0.1918 | 0.1919 | 0.1922 | 0.1927 | 0.1933 | 0.1937 | 0.194 | 0.1944 | 0.1946 | 0.1947 | 0.1948 | 0.1953 | 0.1956 |
| 7 | 44790921 | rs55705735 | na | 0.0429 | 0.00054 | 0.1009 | 0.1009 | 0.1011 | 0.1021 | 0.1049 | 0.1083 | 0.1113 | 0.1135 | 0.1151 | 0.1158 | 0.117 | 0.1178 | 0.1182 | 0.118 | 0.1178 | 0.1174 | 0.1167 | 0.1163 | 0.1161 | 0.1161 | 0.116 | 0.1156 | 0.1145 | 0.1135 | 0.1123 | 0.1118 | 0.1111 | 0.1103 | 0.1097 | 0.109 | 0.108 | 0.1076 | 0.1075 | 0.1076 | 0.1077 | 0.1077 | 0.1077 | 0.1078 | 0.1076 | 0.1072 | 0.1067 | 0.1066 | 0.1064 | 0.1064 | 0.1066 | 0.1066 | 0.1065 | 0.1062 | 0.1059 | 0.1054 |
| 9 | 104165291 | rs75061787 | na | 0.1351 | 0.00142 | 0.0587 | 0.0587 | 0.0586 | 0.0585 | 0.0583 | 0.0578 | 0.0567 | 0.0548 | 0.0539 | 0.0546 | 0.0555 | 0.0565 | 0.057 | 0.0566 | 0.0559 | 0.0555 | 0.0551 | 0.0552 | 0.055 | 0.0546 | 0.0541 | 0.0534 | 0.0526 | 0.0514 | 0.0504 | 0.0494 | 0.0483 | 0.0478 | 0.0471 | 0.0463 | 0.0453 | 0.0446 | 0.0439 | 0.0433 | 0.0429 | 0.0426 | 0.0422 | 0.0417 | 0.041 | 0.0403 | 0.0394 | 0.039 | 0.0387 | 0.0383 | 0.038 | 0.0376 | 0.0371 | 0.0365 | 0.0359 | 0.0353 |
| 9 | 104165291 | rs75061787 | na | 0.1052 | 0.00122 | 0.0569 | 0.0569 | 0.0568 | 0.0563 | 0.0548 | 0.0529 | 0.052 | 0.0528 | 0.0542 | 0.0552 | 0.0564 | 0.0579 | 0.0588 | 0.0591 | 0.0596 | 0.0596 | 0.0595 | 0.0592 | 0.0586 | 0.0578 | 0.057 | 0.0567 | 0.0564 | 0.0563 | 0.0565 | 0.0565 | 0.0564 | 0.0562 | 0.0561 | 0.0562 | 0.0563 | 0.0563 | 0.056 | 0.0557 | 0.0552 | 0.0546 | 0.0538 | 0.0529 | 0.0519 | 0.051 | 0.0501 | 0.0494 | 0.049 | 0.0485 | 0.0483 | 0.048 | 0.0475 | 0.0471 | 0.0465 | 0.0458 |
| 9 | 114238010 | rs2026644 | na | 0.0535 | 0.00063 | 0.1043 | 0.1043 | 0.1043 | 0.1037 | 0.1025 | 0.1008 | 0.0971 | 0.094 | 0.0909 | 0.0862 | 0.0819 | 0.0792 | 0.0776 | 0.0771 | 0.0778 | 0.0788 | 0.0799 | 0.081 | 0.0819 | 0.0827 | 0.0835 | 0.0839 | 0.0843 | 0.0844 | 0.0842 | 0.0838 | 0.0832 | 0.0829 | 0.083 | 0.0829 | 0.0827 | 0.0825 | 0.0822 | 0.0819 | 0.0814 | 0.0808 | 0.08 | 0.0795 | 0.0791 | 0.0786 | 0.0784 | 0.0779 | 0.0777 | 0.0777 | 0.0776 | 0.0774 | 0.0772 | 0.0768 | 0.0763 | 0.0757 |
| 9 | 114238010 | rs2026644 | na | 0.0046 | -0.00017 | 0.1095 | 0.1095 | 0.1094 | 0.1091 | 0.1097 | 0.1095 | 0.1089 | 0.1091 | 0.1101 | 0.1114 | 0.1129 | 0.1145 | 0.1159 | 0.1165 | 0.1172 | 0.1186 | 0.1194 | 0.1197 | 0.1203 | 0.1216 | 0.1226 | 0.1233 | 0.1233 | 0.1236 | 0.1239 | 0.1242 | 0.1244 | 0.1247 | 0.1251 | 0.1252 | 0.1252 | 0.1251 | 0.1251 | 0.125 | 0.125 | 0.1254 | 0.1258 | 0.126 | 0.1262 | 0.1264 | 0.1266 | 0.1269 | 0.127 | 0.1272 | 0.1272 | 0.1272 | 0.1272 | 0.1273 | 0.1273 | 0.1273 |
| 9 | 128935738 | rs1983983 | na | 1.9758 | 0.00396 | 0.125 | 0.125 | 0.1242 | 0.121 | 0.1148 | 0.1079 | 0.0992 | 0.093 | 0.0887 | 0.0846 | 0.0816 | 0.0793 | 0.0771 | 0.0752 | 0.0737 | 0.0717 | 0.0702 | 0.0689 | 0.068 | 0.0675 | 0.067 | 0.0662 | 0.0656 | 0.0651 | 0.0649 | 0.0648 | 0.0646 | 0.0643 | 0.0641 | 0.0638 | 0.0634 | 0.0629 | 0.0622 | 0.0614 | 0.0602 | 0.0589 | 0.0573 | 0.0558 | 0.0541 | 0.0524 | 0.0505 | 0.0484 | 0.046 | 0.0437 | 0.0413 | 0.0388 | 0.0369 | 0.0351 | 0.0334 | 0.0321 |
| 9 | 128935738 | rs1983983 | na | 1.7915 | 0.00361 | 0.1357 | 0.1357 | 0.1356 | 0.1349 | 0.132 | 0.1304 | 0.1293 | 0.1281 | 0.1267 | 0.1261 | 0.1257 | 0.1259 | 0.1264 | 0.1255 | 0.1236 | 0.122 | 0.1203 | 0.1187 | 0.1166 | 0.1145 | 0.113 | 0.1118 | 0.1108 | 0.1094 | 0.1079 | 0.1062 | 0.104 | 0.1022 | 0.1002 | 0.098 | 0.0955 | 0.0925 | 0.0897 | 0.0875 | 0.0852 | 0.0829 | 0.0807 | 0.0788 | 0.0773 | 0.0756 | 0.0738 | 0.0718 | 0.07 | 0.0682 | 0.0664 | 0.0647 | 0.0628 | 0.0608 | 0.0589 | 0.0569 |
| 10 | 8561207 | rs72775239 | na | 3.6944 | 0.00437 | 0.1849 | 0.1849 | 0.1845 | 0.1849 | 0.1853 | 0.1851 | 0.1835 | 0.1816 | 0.1789 | 0.1779 | 0.1778 | 0.1773 | 0.1757 | 0.174 | 0.1721 | 0.17 | 0.1682 | 0.1672 | 0.1661 | 0.1647 | 0.1629 | 0.1608 | 0.1581 | 0.1555 | 0.1533 | 0.1512 | 0.1494 | 0.1479 | 0.1463 | 0.145 | 0.1435 | 0.1416 | 0.1403 | 0.1388 | 0.1372 | 0.1354 | 0.1336 | 0.1321 | 0.1302 | 0.1282 | 0.1263 | 0.1243 | 0.1223 | 0.12 | 0.1174 | 0.1145 | 0.1114 | 0.1086 | 0.1056 | 0.1023 |
| 10 | 8561207 | rs72775239 | na | 1.1623 | 0.00249 | 0.177 | 0.177 | 0.1763 | 0.1721 | 0.1678 | 0.1642 | 0.161 | 0.1589 | 0.1575 | 0.1566 | 0.155 | 0.1539 | 0.1534 | 0.1533 | 0.1524 | 0.1515 | 0.1501 | 0.1483 | 0.1465 | 0.145 | 0.143 | 0.1408 | 0.139 | 0.1372 | 0.1359 | 0.1349 | 0.1339 | 0.1326 | 0.1309 | 0.1291 | 0.127 | 0.1252 | 0.1233 | 0.1218 | 0.1205 | 0.1193 | 0.1183 | 0.1171 | 0.1159 | 0.1144 | 0.1127 | 0.1109 | 0.109 | 0.1075 | 0.106 | 0.1045 | 0.1027 | 0.1006 | 0.0985 | 0.0967 |
| 10 | 70023896 | rs61854810 | 70028315 | 0.2475 | 0.00105 | 0.1848 | 0.1848 | 0.1847 | 0.1829 | 0.1794 | 0.1761 | 0.1743 | 0.174 | 0.1742 | 0.1737 | 0.1743 | 0.1764 | 0.1794 | 0.1809 | 0.1822 | 0.183 | 0.1842 | 0.1859 | 0.1882 | 0.1907 | 0.1931 | 0.1947 | 0.1959 | 0.1969 | 0.198 | 0.1988 | 0.1992 | 0.1992 | 0.1991 | 0.1986 | 0.1978 | 0.1971 | 0.1967 | 0.1961 | 0.1956 | 0.1949 | 0.1939 | 0.1928 | 0.1918 | 0.1906 | 0.189 | 0.1871 | 0.1849 | 0.1827 | 0.1805 | 0.1785 | 0.177 | 0.1756 | 0.1741 | 0.1728 |
| 10 | 70023896 | rs61854810 | 70028315 | 0.3993 | 0.00142 | 0.1704 | 0.1704 | 0.1695 | 0.164 | 0.1565 | 0.1538 | 0.1526 | 0.1534 | 0.1541 | 0.1555 | 0.1557 | 0.1567 | 0.1587 | 0.1608 | 0.1626 | 0.1642 | 0.1643 | 0.1636 | 0.1623 | 0.1604 | 0.1588 | 0.1575 | 0.1566 | 0.156 | 0.1558 | 0.1557 | 0.155 | 0.1544 | 0.1541 | 0.1537 | 0.1536 | 0.1532 | 0.1526 | 0.152 | 0.1516 | 0.1509 | 0.1504 | 0.1501 | 0.1499 | 0.1496 | 0.1494 | 0.1491 | 0.149 | 0.1487 | 0.1483 | 0.1478 | 0.1469 | 0.1458 | 0.1443 | 0.1426 |
| 11 | 9432090 | rs12279202 | na | 0.3479 | 0.00183 | 0.072 | 0.072 | 0.0721 | 0.0719 | 0.0722 | 0.0739 | 0.0752 | 0.0761 | 0.0776 | 0.0799 | 0.0817 | 0.0826 | 0.0823 | 0.0803 | 0.0782 | 0.0764 | 0.0751 | 0.0746 | 0.0748 | 0.0754 | 0.0758 | 0.0762 | 0.0768 | 0.0776 | 0.0782 | 0.0788 | 0.0792 | 0.08 | 0.0808 | 0.0817 | 0.0824 | 0.0831 | 0.0836 | 0.084 | 0.0845 | 0.085 | 0.0855 | 0.0859 | 0.0861 | 0.0861 | 0.0859 | 0.0855 | 0.085 | 0.0843 | 0.0834 | 0.0822 | 0.0809 | 0.0796 | 0.0785 | 0.0774 |
| 11 | 9432090 | rs12279202 | na | 0.1895 | 0.00144 | 0.0773 | 0.0773 | 0.0774 | 0.0779 | 0.0789 | 0.0781 | 0.0768 | 0.0778 | 0.0783 | 0.0793 | 0.0803 | 0.0807 | 0.0811 | 0.0812 | 0.0815 | 0.081 | 0.0801 | 0.0787 | 0.0774 | 0.0757 | 0.0734 | 0.0715 | 0.0702 | 0.0697 | 0.0692 | 0.069 | 0.0689 | 0.0688 | 0.0687 | 0.0686 | 0.0687 | 0.0687 | 0.0685 | 0.0681 | 0.0675 | 0.0666 | 0.0656 | 0.0643 | 0.0629 | 0.0617 | 0.0607 | 0.0598 | 0.0588 | 0.0576 | 0.0567 | 0.0557 | 0.0544 | 0.0531 | 0.0521 | 0.0513 |
| 11 | 101133512 | rs11224678 | na | 3.7616 | 0.00444 | 0.1911 | 0.1911 | 0.1903 | 0.1863 | 0.1843 | 0.1824 | 0.1819 | 0.1816 | 0.1822 | 0.1807 | 0.1769 | 0.1734 | 0.1693 | 0.1671 | 0.1658 | 0.1651 | 0.1643 | 0.1629 | 0.161 | 0.1601 | 0.1582 | 0.156 | 0.1542 | 0.1525 | 0.1503 | 0.1476 | 0.1448 | 0.143 | 0.1414 | 0.1401 | 0.1383 | 0.136 | 0.1336 | 0.1315 | 0.1297 | 0.1277 | 0.1262 | 0.1246 | 0.123 | 0.1212 | 0.1194 | 0.1175 | 0.1153 | 0.1128 | 0.1102 | 0.1077 | 0.1049 | 0.1021 | 0.0993 | 0.0963 |
| 11 | 101133512 | rs11224678 | na | 0.9573 | 0.00215 | 0.1861 | 0.1861 | 0.1857 | 0.1848 | 0.1836 | 0.1837 | 0.1842 | 0.1858 | 0.1881 | 0.1905 | 0.1921 | 0.1933 | 0.1934 | 0.192 | 0.1905 | 0.1888 | 0.1864 | 0.1836 | 0.1818 | 0.1811 | 0.1809 | 0.1797 | 0.1779 | 0.1767 | 0.1758 | 0.1746 | 0.1736 | 0.1721 | 0.1705 | 0.1682 | 0.1658 | 0.164 | 0.1628 | 0.1617 | 0.1608 | 0.1604 | 0.1601 | 0.1597 | 0.1595 | 0.159 | 0.1583 | 0.1577 | 0.1572 | 0.1565 | 0.1556 | 0.1545 | 0.1531 | 0.1516 | 0.1499 | 0.1479 |
| 12 | 56657888 | rs75598973 | na | 3.0456 | 0.0064 | 0.0935 | 0.0935 | 0.0926 | 0.0877 | 0.0811 | 0.0751 | 0.0704 | 0.0634 | 0.0597 | 0.0584 | 0.0575 | 0.0564 | 0.0551 | 0.0539 | 0.0515 | 0.0498 | 0.0483 | 0.0471 | 0.0462 | 0.0455 | 0.0451 | 0.0446 | 0.0438 | 0.0428 | 0.0415 | 0.0403 | 0.0392 | 0.0382 | 0.0371 | 0.0357 | 0.0342 | 0.0323 | 0.0305 | 0.0288 | 0.0271 | 0.025 | 0.0231 | 0.0217 | 0.0204 | 0.019 | 0.0178 | 0.0166 | 0.0158 | 0.015 | 0.0141 | 0.0131 | 0.0121 | 0.011 | 0.0097 | 0.0088 |
| 12 | 56657888 | rs75598973 | na | 2.8512 | 0.00684 | 0.0902 | 0.0902 | 0.0893 | 0.0876 | 0.0869 | 0.0865 | 0.0862 | 0.0854 | 0.0831 | 0.0796 | 0.0758 | 0.0735 | 0.0706 | 0.0684 | 0.0662 | 0.0643 | 0.0622 | 0.0599 | 0.058 | 0.0563 | 0.0549 | 0.0543 | 0.054 | 0.0534 | 0.0528 | 0.0522 | 0.0513 | 0.0502 | 0.0489 | 0.0478 | 0.0468 | 0.046 | 0.0449 | 0.0437 | 0.0428 | 0.0417 | 0.0406 | 0.0396 | 0.0386 | 0.0377 | 0.0371 | 0.0365 | 0.0358 | 0.035 | 0.0342 | 0.0332 | 0.0321 | 0.0308 | 0.0295 | 0.028 |
| 12 | 133750212 | rs61958756 | na | 0.0048 | -0.00012 | 0.3067 | 0.3067 | 0.3066 | 0.305 | 0.3018 | 0.2965 | 0.2936 | 0.2924 | 0.2909 | 0.2898 | 0.2892 | 0.2899 | 0.2913 | 0.293 | 0.2945 | 0.2958 | 0.2968 | 0.2987 | 0.3013 | 0.3032 | 0.3039 | 0.3044 | 0.3051 | 0.3055 | 0.3059 | 0.3058 | 0.3059 | 0.3061 | 0.307 | 0.3081 | 0.3089 | 0.3093 | 0.3094 | 0.3102 | 0.3115 | 0.3122 | 0.3126 | 0.3125 | 0.3121 | 0.3115 | 0.3109 | 0.3104 | 0.3098 | 0.3094 | 0.3096 | 0.3098 | 0.3103 | 0.3106 | 0.3108 | 0.3107 |
| 12 | 133750212 | rs61958756 | na | 0.3383 | 0.00103 | 0.3138 | 0.3138 | 0.3129 | 0.31 | 0.307 | 0.3031 | 0.3015 | 0.3003 | 0.3007 | 0.3028 | 0.308 | 0.3133 | 0.3168 | 0.3196 | 0.3204 | 0.3213 | 0.3228 | 0.3238 | 0.3243 | 0.3253 | 0.3264 | 0.3268 | 0.3269 | 0.3267 | 0.3272 | 0.3276 | 0.3276 | 0.3275 | 0.3265 | 0.3256 | 0.3245 | 0.3236 | 0.3226 | 0.3221 | 0.3217 | 0.3211 | 0.3205 | 0.3198 | 0.3192 | 0.3184 | 0.3179 | 0.3171 | 0.3162 | 0.315 | 0.3139 | 0.3126 | 0.3118 | 0.3109 | 0.3092 | 0.3072 |
| 13 | 27959981 | rs77509251 | na | 1.3918 | 0.00259 | 0.2007 | 0.2007 | 0.2005 | 0.199 | 0.1969 | 0.1926 | 0.1861 | 0.1808 | 0.1779 | 0.1752 | 0.1718 | 0.1669 | 0.1633 | 0.1603 | 0.1577 | 0.1555 | 0.1529 | 0.1499 | 0.1474 | 0.1452 | 0.1433 | 0.1415 | 0.1397 | 0.1379 | 0.1363 | 0.1347 | 0.1333 | 0.1319 | 0.1302 | 0.1286 | 0.1273 | 0.1263 | 0.1255 | 0.1248 | 0.124 | 0.1229 | 0.1217 | 0.1206 | 0.1193 | 0.1179 | 0.1164 | 0.1149 | 0.1133 | 0.1115 | 0.1099 | 0.1082 | 0.1063 | 0.1048 | 0.1031 | 0.1013 |
| 13 | 27959981 | rs77509251 | na | 1.7376 | 0.00276 | 0.1981 | 0.1981 | 0.1978 | 0.1981 | 0.1984 | 0.1992 | 0.2029 | 0.206 | 0.2076 | 0.2074 | 0.2067 | 0.2056 | 0.2033 | 0.2004 | 0.1967 | 0.1924 | 0.1888 | 0.1858 | 0.1826 | 0.1789 | 0.1756 | 0.1727 | 0.1694 | 0.1664 | 0.1643 | 0.1627 | 0.1615 | 0.1601 | 0.1588 | 0.1573 | 0.1561 | 0.1551 | 0.1542 | 0.1537 | 0.1531 | 0.1524 | 0.1516 | 0.1507 | 0.1498 | 0.1487 | 0.1472 | 0.1454 | 0.1433 | 0.1414 | 0.1393 | 0.1372 | 0.1349 | 0.1327 | 0.1303 | 0.1277 |
| 14 | 68909571 | rs112506612 | na | 1.8652 | 0.00271 | 0.2412 | 0.2412 | 0.24 | 0.2336 | 0.2278 | 0.2256 | 0.2235 | 0.2207 | 0.2182 | 0.2184 | 0.218 | 0.2168 | 0.2154 | 0.2143 | 0.2121 | 0.2109 | 0.2093 | 0.2073 | 0.2054 | 0.2031 | 0.1995 | 0.195 | 0.1906 | 0.1864 | 0.1826 | 0.1795 | 0.1764 | 0.1731 | 0.1691 | 0.1658 | 0.1632 | 0.1609 | 0.1589 | 0.1568 | 0.1548 | 0.1528 | 0.151 | 0.1487 | 0.1459 | 0.1433 | 0.1411 | 0.1393 | 0.1373 | 0.1351 | 0.1331 | 0.1312 | 0.1286 | 0.126 | 0.1234 | 0.1205 |
| 14 | 68909571 | rs112506612 | na | 2.256 | 0.00291 | 0.2338 | 0.2338 | 0.2321 | 0.2259 | 0.2194 | 0.2164 | 0.2145 | 0.2139 | 0.2128 | 0.2108 | 0.2105 | 0.2092 | 0.2077 | 0.2072 | 0.208 | 0.2084 | 0.2086 | 0.2092 | 0.2101 | 0.2105 | 0.2104 | 0.2099 | 0.2087 | 0.2075 | 0.2067 | 0.206 | 0.2056 | 0.2051 | 0.2047 | 0.2039 | 0.2031 | 0.2019 | 0.2004 | 0.1986 | 0.1971 | 0.1954 | 0.1932 | 0.1907 | 0.1883 | 0.1859 | 0.1833 | 0.1808 | 0.1778 | 0.1746 | 0.1715 | 0.1684 | 0.1651 | 0.1618 | 0.1587 | 0.1555 |
| 14 | 69055743 | rs61985819 | na | 0.8637 | 0.00181 | 0.24 | 0.24 | 0.2399 | 0.2396 | 0.2393 | 0.2367 | 0.2342 | 0.2294 | 0.2265 | 0.2256 | 0.2248 | 0.2253 | 0.2251 | 0.2241 | 0.2219 | 0.2185 | 0.2143 | 0.2096 | 0.2059 | 0.2023 | 0.1992 | 0.1967 | 0.1948 | 0.1937 | 0.193 | 0.1923 | 0.1912 | 0.1898 | 0.1882 | 0.187 | 0.1853 | 0.1834 | 0.1817 | 0.1797 | 0.1774 | 0.1751 | 0.1726 | 0.1703 | 0.1682 | 0.1666 | 0.165 | 0.1634 | 0.1614 | 0.1592 | 0.1567 | 0.1546 | 0.1522 | 0.1498 | 0.1475 | 0.1454 |
| 14 | 69055743 | rs61985819 | na | 1.0717 | 0.00205 | 0.2442 | 0.2442 | 0.2438 | 0.2411 | 0.24 | 0.2395 | 0.2402 | 0.2388 | 0.2368 | 0.2349 | 0.2331 | 0.2319 | 0.2305 | 0.2294 | 0.228 | 0.2259 | 0.2229 | 0.2204 | 0.2181 | 0.2163 | 0.2137 | 0.2115 | 0.2093 | 0.2068 | 0.2055 | 0.2047 | 0.2038 | 0.2025 | 0.2012 | 0.2003 | 0.1996 | 0.1988 | 0.1978 | 0.1967 | 0.1957 | 0.1944 | 0.193 | 0.1915 | 0.1908 | 0.1902 | 0.1893 | 0.1886 | 0.1877 | 0.1865 | 0.1856 | 0.1845 | 0.183 | 0.1815 | 0.1798 | 0.1778 |
| 14 | 74209097 | rs11624425 | 74204686 | 2.2336 | 0.00259 | 0.3739 | 0.3739 | 0.3727 | 0.3703 | 0.3635 | 0.3561 | 0.3516 | 0.3467 | 0.3412 | 0.3359 | 0.3312 | 0.3277 | 0.3256 | 0.323 | 0.3199 | 0.3176 | 0.3151 | 0.3124 | 0.3103 | 0.3076 | 0.3057 | 0.3037 | 0.3016 | 0.2989 | 0.2952 | 0.2907 | 0.2864 | 0.2825 | 0.2784 | 0.2746 | 0.2705 | 0.267 | 0.2632 | 0.2591 | 0.2551 | 0.2519 | 0.2493 | 0.2467 | 0.2441 | 0.2413 | 0.238 | 0.2345 | 0.2312 | 0.228 | 0.2249 | 0.2221 | 0.2188 | 0.2155 | 0.2117 | 0.2086 |
| 14 | 74209097 | rs11624425 | 74204686 | 1.5617 | 0.0021 | 0.3802 | 0.3802 | 0.3795 | 0.3771 | 0.3758 | 0.3735 | 0.3692 | 0.3608 | 0.3508 | 0.3448 | 0.3402 | 0.3368 | 0.3339 | 0.3315 | 0.3298 | 0.3291 | 0.3276 | 0.3249 | 0.3215 | 0.3182 | 0.3159 | 0.3141 | 0.3126 | 0.3105 | 0.3086 | 0.3066 | 0.3049 | 0.3028 | 0.3006 | 0.2981 | 0.2967 | 0.2953 | 0.2932 | 0.2913 | 0.2896 | 0.2884 | 0.2871 | 0.2858 | 0.2847 | 0.2832 | 0.2815 | 0.2796 | 0.278 | 0.2762 | 0.2742 | 0.2718 | 0.2691 | 0.2665 | 0.2637 | 0.2607 |
| 14 | 76780815 | rs11626073 | na | 1.4416 | 0.00244 | 0.2277 | 0.2277 | 0.227 | 0.2241 | 0.2206 | 0.2188 | 0.2169 | 0.2177 | 0.2209 | 0.2243 | 0.2285 | 0.2316 | 0.2341 | 0.237 | 0.2381 | 0.2388 | 0.2395 | 0.2405 | 0.2423 | 0.2438 | 0.2451 | 0.2457 | 0.2467 | 0.2474 | 0.2474 | 0.247 | 0.246 | 0.2448 | 0.2433 | 0.2415 | 0.24 | 0.2384 | 0.2369 | 0.2356 | 0.2346 | 0.2334 | 0.2328 | 0.2322 | 0.2315 | 0.2308 | 0.2295 | 0.2279 | 0.2258 | 0.2236 | 0.2211 | 0.2182 | 0.2145 | 0.2104 | 0.2066 | 0.2034 |
| 14 | 76780815 | rs11626073 | na | 0.6477 | 0.00161 | 0.2255 | 0.2255 | 0.2251 | 0.2245 | 0.224 | 0.2242 | 0.2226 | 0.2182 | 0.2151 | 0.2128 | 0.211 | 0.2085 | 0.2061 | 0.2029 | 0.1992 | 0.1969 | 0.1942 | 0.1919 | 0.1902 | 0.1891 | 0.1876 | 0.1857 | 0.1838 | 0.1814 | 0.1792 | 0.1773 | 0.1754 | 0.1735 | 0.1712 | 0.1688 | 0.1668 | 0.1647 | 0.1625 | 0.1607 | 0.1586 | 0.157 | 0.1558 | 0.1546 | 0.1538 | 0.1531 | 0.1525 | 0.1518 | 0.151 | 0.15 | 0.1491 | 0.1479 | 0.1466 | 0.1453 | 0.1439 | 0.1423 |
| 15 | 85203704 | rs62021209 | na | 0.0031 | 0.0001 | 0.2181 | 0.2181 | 0.2183 | 0.2178 | 0.2171 | 0.2198 | 0.2221 | 0.2253 | 0.2273 | 0.2303 | 0.232 | 0.234 | 0.2356 | 0.2368 | 0.2383 | 0.2405 | 0.2437 | 0.2471 | 0.2509 | 0.2548 | 0.2583 | 0.2611 | 0.2641 | 0.2679 | 0.2713 | 0.2743 | 0.2771 | 0.2795 | 0.2816 | 0.2832 | 0.2847 | 0.2858 | 0.2867 | 0.2875 | 0.288 | 0.2878 | 0.2876 | 0.2872 | 0.2869 | 0.2862 | 0.2854 | 0.2846 | 0.2836 | 0.2822 | 0.281 | 0.28 | 0.2791 | 0.2787 | 0.278 | 0.2773 |
| 15 | 85203704 | rs62021209 | na | 0.09 | 0.00056 | 0.2108 | 0.2108 | 0.2107 | 0.2103 | 0.2108 | 0.2127 | 0.218 | 0.2245 | 0.2302 | 0.236 | 0.2413 | 0.2451 | 0.2476 | 0.2489 | 0.249 | 0.2495 | 0.2504 | 0.2513 | 0.2519 | 0.2526 | 0.2532 | 0.253 | 0.2527 | 0.2525 | 0.2521 | 0.2514 | 0.2498 | 0.249 | 0.2486 | 0.2482 | 0.2483 | 0.2485 | 0.2488 | 0.2489 | 0.249 | 0.2489 | 0.2489 | 0.2488 | 0.2486 | 0.2483 | 0.2482 | 0.2482 | 0.248 | 0.2475 | 0.2468 | 0.246 | 0.2449 | 0.2437 | 0.2427 | 0.2421 |
| 16 | 3517787 | rs1690443 | 3530658 | 2.8467 | 0.00391 | 0.1607 | 0.1607 | 0.1601 | 0.1591 | 0.1573 | 0.1567 | 0.1574 | 0.1575 | 0.1563 | 0.1562 | 0.1556 | 0.1556 | 0.1559 | 0.1557 | 0.155 | 0.1542 | 0.1533 | 0.1526 | 0.1516 | 0.1502 | 0.1485 | 0.1469 | 0.1464 | 0.1458 | 0.1449 | 0.1441 | 0.1433 | 0.1425 | 0.1416 | 0.1405 | 0.1393 | 0.1382 | 0.1374 | 0.137 | 0.1364 | 0.1355 | 0.1342 | 0.1326 | 0.1308 | 0.129 | 0.1269 | 0.1246 | 0.1221 | 0.1193 | 0.1165 | 0.1145 | 0.1127 | 0.1106 | 0.1083 | 0.1054 |
| 16 | 3517787 | rs1690443 | 3530658 | 1.0179 | 0.00225 | 0.165 | 0.165 | 0.1647 | 0.1634 | 0.1632 | 0.1651 | 0.1661 | 0.1653 | 0.1637 | 0.1613 | 0.1583 | 0.1561 | 0.1555 | 0.1554 | 0.1547 | 0.1526 | 0.1502 | 0.1481 | 0.1461 | 0.1441 | 0.1424 | 0.141 | 0.1398 | 0.1381 | 0.1364 | 0.135 | 0.1337 | 0.1321 | 0.1305 | 0.129 | 0.1276 | 0.1259 | 0.1239 | 0.1224 | 0.1215 | 0.1208 | 0.1198 | 0.1188 | 0.1178 | 0.1169 | 0.1166 | 0.1162 | 0.1159 | 0.1155 | 0.115 | 0.1142 | 0.1132 | 0.1121 | 0.1107 | 0.1091 |
| 19 | 49382969 | rs55706579 | na | 0.7811 | 0.00215 | 0.1427 | 0.1427 | 0.1424 | 0.1423 | 0.1451 | 0.148 | 0.148 | 0.1479 | 0.1474 | 0.1472 | 0.147 | 0.1468 | 0.1459 | 0.1445 | 0.142 | 0.1392 | 0.1371 | 0.1354 | 0.1337 | 0.1324 | 0.132 | 0.1316 | 0.1307 | 0.1302 | 0.1297 | 0.1289 | 0.1278 | 0.127 | 0.1262 | 0.1253 | 0.1244 | 0.1234 | 0.1223 | 0.1211 | 0.1205 | 0.1203 | 0.1201 | 0.1198 | 0.1194 | 0.1189 | 0.1181 | 0.1173 | 0.1168 | 0.1162 | 0.1156 | 0.1148 | 0.1137 | 0.1125 | 0.1111 | 0.1096 |
| 19 | 49382969 | rs55706579 | na | 2.0567 | 0.00386 | 0.1446 | 0.1446 | 0.1442 | 0.1428 | 0.1401 | 0.1359 | 0.134 | 0.1325 | 0.1303 | 0.1266 | 0.1227 | 0.1199 | 0.1183 | 0.1162 | 0.1134 | 0.1117 | 0.11 | 0.1078 | 0.1058 | 0.1041 | 0.1023 | 0.1003 | 0.0981 | 0.0965 | 0.0955 | 0.0946 | 0.0937 | 0.0929 | 0.092 | 0.0911 | 0.0901 | 0.089 | 0.0881 | 0.0872 | 0.0865 | 0.0858 | 0.085 | 0.0844 | 0.0837 | 0.0828 | 0.0819 | 0.0809 | 0.0799 | 0.0789 | 0.0777 | 0.0763 | 0.0747 | 0.0729 | 0.0711 | 0.0693 |
| 20 | 756531 | rs6140159 | 757491 | 0.1663 | 0.0009 | 0.1829 | 0.1829 | 0.1824 | 0.1815 | 0.1811 | 0.1816 | 0.1821 | 0.1802 | 0.1774 | 0.175 | 0.1737 | 0.1717 | 0.17 | 0.1686 | 0.1673 | 0.1659 | 0.1647 | 0.1643 | 0.1643 | 0.1638 | 0.1634 | 0.1628 | 0.1628 | 0.163 | 0.1642 | 0.1658 | 0.1673 | 0.1692 | 0.1707 | 0.1721 | 0.1733 | 0.1743 | 0.175 | 0.1754 | 0.1758 | 0.1759 | 0.176 | 0.176 | 0.1758 | 0.1753 | 0.1749 | 0.1746 | 0.1741 | 0.1735 | 0.1726 | 0.1715 | 0.1703 | 0.1689 | 0.168 | 0.167 |
| 20 | 756531 | rs6140159 | 757491 | 0.1819 | 0.00098 | 0.188 | 0.188 | 0.1884 | 0.1901 | 0.1904 | 0.1891 | 0.1869 | 0.1848 | 0.1827 | 0.1821 | 0.1813 | 0.1808 | 0.1801 | 0.179 | 0.1779 | 0.1777 | 0.1777 | 0.1773 | 0.1769 | 0.1765 | 0.1762 | 0.1761 | 0.1757 | 0.1753 | 0.1748 | 0.1747 | 0.1744 | 0.1743 | 0.1743 | 0.1744 | 0.1742 | 0.174 | 0.1737 | 0.1734 | 0.1735 | 0.1739 | 0.1745 | 0.175 | 0.1753 | 0.1755 | 0.1756 | 0.1759 | 0.176 | 0.176 | 0.1758 | 0.1755 | 0.1747 | 0.1739 | 0.1732 | 0.1722 |
| 21 | 43221111 | rs9983923 | na | 0.3756 | 0.00205 | 0.0742 | 0.0742 | 0.0741 | 0.0743 | 0.0751 | 0.0748 | 0.0754 | 0.0759 | 0.077 | 0.0788 | 0.0805 | 0.0822 | 0.0841 | 0.0855 | 0.0868 | 0.0884 | 0.0893 | 0.0896 | 0.0903 | 0.0906 | 0.0902 | 0.0894 | 0.0886 | 0.0873 | 0.0864 | 0.0852 | 0.0839 | 0.0827 | 0.0813 | 0.0798 | 0.078 | 0.0758 | 0.0734 | 0.0716 | 0.0696 | 0.0673 | 0.065 | 0.0628 | 0.0606 | 0.0584 | 0.0562 | 0.0539 | 0.0515 | 0.0493 | 0.0475 | 0.0459 | 0.0439 | 0.042 | 0.0409 | 0.04 |
| 21 | 43221111 | rs9983923 | na | 0.0403 | 0.00068 | 0.0647 | 0.0647 | 0.0642 | 0.062 | 0.0586 | 0.057 | 0.0571 | 0.058 | 0.0597 | 0.0605 | 0.061 | 0.0611 | 0.0605 | 0.0596 | 0.0586 | 0.0574 | 0.0572 | 0.0567 | 0.0561 | 0.0553 | 0.0553 | 0.0552 | 0.0554 | 0.0556 | 0.0556 | 0.0554 | 0.055 | 0.0548 | 0.0545 | 0.0542 | 0.0541 | 0.0541 | 0.0542 | 0.0548 | 0.0553 | 0.0558 | 0.0561 | 0.0565 | 0.0568 | 0.057 | 0.0572 | 0.0573 | 0.0573 | 0.0571 | 0.0569 | 0.0566 | 0.0562 | 0.0557 | 0.0552 | 0.0546 |

Table S11: CLUES derived allele frequency estimates over last 500 generations and respective summary statistics for trans-eQTL aSNPs

| | | | | | | generations | | | | | | | | | | | | | | | | | | | | | | | | | | | | | | | | | | | | | | | | | | | | | | | | | |
|---|---|---|---|---|---|---|---|---|---|---|---|---|---|---|---|---|---|---|---|---|---|---|---|---|---|---|---|---|---|---|---|---|---|---|---|---|---|---|---|---|---|---|---|---|---|---|---|---|---|---|---|---|---|---|---|
| chr | pos | rs | proxy_for | logLR | s | g0 | g10 | g20 | g30 | g40 | g50 | g60 | g70 | g80 | g90 | g100 | g110 | g120 | g130 | g140 | g150 | g160 | g170 | g180 | g190 | g200 | g210 | g220 | g230 | g240 | g250 | g260 | g270 | g280 | g290 | g300 | g310 | g320 | g330 | g340 | g350 | g360 | g370 | g380 | g390 | g400 | g410 | g420 | g430 | g440 | g450 | g460 | g470 | g480 | g490 |
| 1 | 22587728 | rs72647484 | na | 1.0488 | 0.00332 | 0.101 | 0.101 | 0.1007 | 0.0993 | 0.0974 | 0.0957 | 0.0953 | 0.0961 | 0.0968 | 0.0969 | 0.0974 | 0.098 | 0.0981 | 0.0973 | 0.096 | 0.0939 | 0.0922 | 0.0896 | 0.0867 | 0.0847 | 0.0834 | 0.0826 | 0.0818 | 0.081 | 0.0802 | 0.0788 | 0.0775 | 0.0763 | 0.0755 | 0.0747 | 0.0738 | 0.0729 | 0.0721 | 0.0714 | 0.0704 | 0.0693 | 0.068 | 0.0667 | 0.0654 | 0.064 | 0.0627 | 0.0613 | 0.0596 | 0.0584 | 0.0573 | 0.0564 | 0.0554 | 0.0543 | 0.0531 | 0.0517 |
| 1 | 22587728 | rs72647484 | na | 1.7667 | 0.0042 | 0.0987 | 0.0987 | 0.0986 | 0.0987 | 0.0987 | 0.0995 | 0.1 | 0.0986 | 0.0978 | 0.0972 | 0.0969 | 0.0971 | 0.0965 | 0.0953 | 0.0937 | 0.092 | 0.0918 | 0.092 | 0.0926 | 0.0927 | 0.093 | 0.0936 | 0.0938 | 0.0936 | 0.0933 | 0.0928 | 0.0921 | 0.0915 | 0.0909 | 0.0901 | 0.0892 | 0.0881 | 0.0872 | 0.0866 | 0.0863 | 0.0859 | 0.0856 | 0.0853 | 0.085 | 0.0845 | 0.0837 | 0.0828 | 0.0817 | 0.0806 | 0.0792 | 0.0776 | 0.0757 | 0.0734 | 0.0707 | 0.0685 |
| 1 | 169099037 | rs10919070 | na | 0.6163 | 0.00181 | 0.1535 | 0.1535 | 0.1533 | 0.1524 | 0.151 | 0.1488 | 0.1478 | 0.1464 | 0.1461 | 0.1452 | 0.1439 | 0.1419 | 0.1415 | 0.1411 | 0.1398 | 0.1383 | 0.137 | 0.1366 | 0.136 | 0.1349 | 0.1335 | 0.1318 | 0.1306 | 0.1288 | 0.1269 | 0.1255 | 0.1246 | 0.1234 | 0.1222 | 0.1211 | 0.1201 | 0.119 | 0.1188 | 0.1183 | 0.1174 | 0.1163 | 0.1151 | 0.1139 | 0.1129 | 0.112 | 0.1114 | 0.111 | 0.1104 | 0.1098 | 0.1093 | 0.1087 | 0.108 | 0.107 | 0.1058 | 0.1044 |
| 1 | 169099037 | rs10919070 | na | 0.7393 | 0.002 | 0.1514 | 0.1514 | 0.1512 | 0.1512 | 0.1511 | 0.149 | 0.1495 | 0.1501 | 0.1499 | 0.1479 | 0.1458 | 0.144 | 0.1428 | 0.1419 | 0.1414 | 0.1413 | 0.1403 | 0.1396 | 0.1385 | 0.1369 | 0.1354 | 0.1336 | 0.1315 | 0.129 | 0.1268 | 0.1244 | 0.1222 | 0.1201 | 0.1178 | 0.1158 | 0.1139 | 0.1121 | 0.1106 | 0.109 | 0.1077 | 0.1063 | 0.1048 | 0.103 | 0.101 | 0.0989 | 0.0966 | 0.0942 | 0.092 | 0.0897 | 0.0875 | 0.0852 | 0.0831 | 0.0809 | 0.0785 | 0.0764 |
| 1 | 169099483 | rs10919071 | na | 2.9437 | 0.00449 | 0.1474 | 0.1474 | 0.1468 | 0.1454 | 0.1441 | 0.1417 | 0.139 | 0.1376 | 0.1371 | 0.1346 | 0.1306 | 0.1281 | 0.126 | 0.1247 | 0.1235 | 0.1224 | 0.1206 | 0.1191 | 0.1173 | 0.1157 | 0.1146 | 0.1136 | 0.1128 | 0.1119 | 0.1113 | 0.1103 | 0.1091 | 0.1078 | 0.1066 | 0.1052 | 0.104 | 0.1026 | 0.101 | 0.0989 | 0.0966 | 0.0941 | 0.0915 | 0.0891 | 0.0866 | 0.0844 | 0.0821 | 0.0801 | 0.0781 | 0.0761 | 0.0742 | 0.0722 | 0.0699 | 0.0676 | 0.0654 | 0.0634 |
| 1 | 169099483 | rs10919071 | na | 0.9419 | 0.00229 | 0.1482 | 0.1482 | 0.148 | 0.1471 | 0.1465 | 0.1476 | 0.148 | 0.1483 | 0.1489 | 0.1492 | 0.1494 | 0.1485 | 0.1468 | 0.146 | 0.146 | 0.1463 | 0.1468 | 0.1474 | 0.1479 | 0.1481 | 0.1486 | 0.1495 | 0.1502 | 0.1503 | 0.1503 | 0.1502 | 0.1504 | 0.1501 | 0.1495 | 0.1491 | 0.1484 | 0.1473 | 0.1463 | 0.1457 | 0.1453 | 0.1448 | 0.1441 | 0.1437 | 0.1433 | 0.1428 | 0.1421 | 0.1413 | 0.1403 | 0.1392 | 0.1381 | 0.1368 | 0.1351 | 0.1332 | 0.1311 | 0.1291 |
| 3 | 46173072 | rs13063635 | na | 3.6998 | 0.00432 | 0.1792 | 0.1792 | 0.1783 | 0.1754 | 0.1726 | 0.1692 | 0.1655 | 0.1639 | 0.1638 | 0.1631 | 0.1622 | 0.1605 | 0.1582 | 0.1544 | 0.1504 | 0.1484 | 0.1466 | 0.1448 | 0.1432 | 0.1415 | 0.141 | 0.1403 | 0.1396 | 0.1389 | 0.1385 | 0.138 | 0.1373 | 0.1363 | 0.135 | 0.1336 | 0.1329 | 0.1323 | 0.1316 | 0.1307 | 0.1294 | 0.1278 | 0.1259 | 0.1238 | 0.1219 | 0.12 | 0.1181 | 0.1164 | 0.1148 | 0.113 | 0.111 | 0.1087 | 0.1066 | 0.1041 | 0.1017 | 0.0992 |
| 3 | 46173072 | rs13063635 | na | 4.4199 | 0.0051 | 0.1788 | 0.1788 | 0.1778 | 0.1761 | 0.1746 | 0.1707 | 0.1678 | 0.1648 | 0.161 | 0.1579 | 0.1563 | 0.1553 | 0.1547 | 0.1543 | 0.1544 | 0.1534 | 0.1519 | 0.1501 | 0.1484 | 0.1476 | 0.1472 | 0.1469 | 0.1466 | 0.1459 | 0.1448 | 0.1436 | 0.142 | 0.14 | 0.1376 | 0.1348 | 0.1314 | 0.128 | 0.1255 | 0.1236 | 0.1219 | 0.1198 | 0.118 | 0.1162 | 0.1142 | 0.1124 | 0.1103 | 0.1083 | 0.106 | 0.1035 | 0.1006 | 0.0974 | 0.0938 | 0.0905 | 0.0879 | 0.0851 |
| 3 | 46235201 | rs13098911 | na | 1.8712 | 0.00371 | 0.1373 | 0.1373 | 0.1369 | 0.1351 | 0.1318 | 0.1285 | 0.1273 | 0.1257 | 0.1226 | 0.1198 | 0.1166 | 0.1116 | 0.1068 | 0.1042 | 0.1023 | 0.1008 | 0.0987 | 0.0967 | 0.0945 | 0.0914 | 0.0887 | 0.0865 | 0.0849 | 0.0829 | 0.0806 | 0.0785 | 0.077 | 0.0753 | 0.0734 | 0.0716 | 0.0695 | 0.0676 | 0.0658 | 0.064 | 0.0626 | 0.0613 | 0.06 | 0.0588 | 0.0579 | 0.0569 | 0.0558 | 0.0546 | 0.0532 | 0.0517 | 0.05 | 0.0481 | 0.0461 | 0.0446 | 0.043 | 0.0415 |
| 3 | 46235201 | rs13098911 | na | 1.872 | 0.00396 | 0.1278 | 0.1278 | 0.1275 | 0.1251 | 0.1211 | 0.1178 | 0.1148 | 0.1114 | 0.1084 | 0.1042 | 0.1011 | 0.0983 | 0.0942 | 0.0902 | 0.087 | 0.0833 | 0.0797 | 0.077 | 0.0747 | 0.0721 | 0.0697 | 0.0675 | 0.0653 | 0.0633 | 0.0616 | 0.0601 | 0.0585 | 0.0572 | 0.0562 | 0.055 | 0.0541 | 0.0533 | 0.0526 | 0.052 | 0.0513 | 0.0505 | 0.0497 | 0.0491 | 0.0484 | 0.0481 | 0.0477 | 0.0473 | 0.0468 | 0.0463 | 0.0457 | 0.045 | 0.0441 | 0.0431 | 0.0419 | 0.0406 |
| 6 | 52281072 | rs56032325 | na | 0.001 | -0.0001 | 0.0568 | 0.0568 | 0.0571 | 0.0587 | 0.0602 | 0.0606 | 0.061 | 0.0615 | 0.0617 | 0.0616 | 0.0612 | 0.0601 | 0.0581 | 0.0568 | 0.0563 | 0.0555 | 0.055 | 0.0546 | 0.054 | 0.0532 | 0.0526 | 0.0522 | 0.0521 | 0.0518 | 0.0512 | 0.0505 | 0.0495 | 0.0488 | 0.0482 | 0.048 | 0.0479 | 0.0478 | 0.0476 | 0.0473 | 0.0469 | 0.0463 | 0.0457 | 0.0451 | 0.0446 | 0.0446 | 0.0447 | 0.0448 | 0.0449 | 0.0449 | 0.0449 | 0.0448 | 0.0447 | 0.0445 | 0.0443 | 0.044 |
| 6 | 52281072 | rs56032325 | na | 0.3272 | 0.00225 | 0.0556 | 0.0556 | 0.0556 | 0.0563 | 0.0573 | 0.0579 | 0.0578 | 0.0572 | 0.0573 | 0.0577 | 0.0581 | 0.0596 | 0.0611 | 0.0623 | 0.0632 | 0.0639 | 0.0647 | 0.0655 | 0.0664 | 0.0675 | 0.0688 | 0.0703 | 0.0718 | 0.073 | 0.0738 | 0.0744 | 0.0747 | 0.0749 | 0.0753 | 0.0756 | 0.0757 | 0.0757 | 0.0758 | 0.0755 | 0.0755 | 0.0753 | 0.0749 | 0.0743 | 0.0738 | 0.0733 | 0.0727 | 0.072 | 0.0714 | 0.0706 | 0.0697 | 0.0686 | 0.0676 | 0.0666 | 0.0655 | 0.0643 |
| 7 | 46402269 | rs7811653 | na | 0.0847 | -0.00088 | 0.0646 | 0.0646 | 0.0646 | 0.0648 | 0.0658 | 0.0655 | 0.0665 | 0.0682 | 0.0695 | 0.0698 | 0.0702 | 0.0713 | 0.0726 | 0.0741 | 0.0761 | 0.0782 | 0.0802 | 0.0819 | 0.0831 | 0.0842 | 0.0849 | 0.0855 | 0.0855 | 0.0856 | 0.0855 | 0.0851 | 0.0847 | 0.0845 | 0.0846 | 0.085 | 0.0852 | 0.0851 | 0.085 | 0.085 | 0.0854 | 0.0862 | 0.0872 | 0.088 | 0.089 | 0.0901 | 0.0912 | 0.0921 | 0.093 | 0.0939 | 0.0947 | 0.0954 | 0.096 | 0.0965 | 0.0969 | 0.0971 |
| 7 | 46402269 | rs7811653 | na | 0.3802 | -0.00181 | 0.0663 | 0.0663 | 0.0665 | 0.0667 | 0.0678 | 0.069 | 0.0696 | 0.0703 | 0.0731 | 0.0757 | 0.0776 | 0.0781 | 0.0777 | 0.0775 | 0.0774 | 0.0784 | 0.0802 | 0.0815 | 0.0824 | 0.0832 | 0.084 | 0.085 | 0.0864 | 0.0872 | 0.0878 | 0.088 | 0.0879 | 0.0879 | 0.0877 | 0.0877 | 0.088 | 0.0884 | 0.089 | 0.0897 | 0.0905 | 0.0914 | 0.0924 | 0.0932 | 0.0939 | 0.0946 | 0.0952 | 0.0956 | 0.0961 | 0.0964 | 0.0969 | 0.0974 | 0.0981 | 0.099 | 0.0997 | 0.1004 |
| 11 | 100453046 | rs72996113 | na | 0.0365 | -0.00059 | 0.9252 | 0.9252 | 0.9253 | 0.9257 | 0.9274 | 0.928 | 0.9281 | 0.9278 | 0.9268 | 0.9261 | 0.9256 | 0.9264 | 0.9271 | 0.928 | 0.9275 | 0.9271 | 0.9271 | 0.9274 | 0.9278 | 0.9285 | 0.9288 | 0.9286 | 0.9281 | 0.9273 | 0.9263 | 0.9254 | 0.9246 | 0.9239 | 0.9232 | 0.9228 | 0.9224 | 0.9222 | 0.9222 | 0.922 | 0.9215 | 0.921 | 0.9206 | 0.9202 | 0.9195 | 0.919 | 0.9185 | 0.918 | 0.9176 | 0.9174 | 0.9174 | 0.9175 | 0.9176 | 0.9175 | 0.9175 | 0.9176 |
| 11 | 100453046 | rs72996113 | na | 0.2055 | -0.00146 | 0.9172 | 0.9172 | 0.9172 | 0.9174 | 0.9187 | 0.9181 | 0.9151 | 0.9115 | 0.9093 | 0.9076 | 0.9064 | 0.9057 | 0.9055 | 0.9055 | 0.9056 | 0.9058 | 0.9057 | 0.9058 | 0.9059 | 0.9061 | 0.9064 | 0.9065 | 0.9067 | 0.9075 | 0.9084 | 0.9091 | 0.9094 | 0.9098 | 0.9105 | 0.9112 | 0.9116 | 0.9123 | 0.9131 | 0.9137 | 0.9143 | 0.9151 | 0.9159 | 0.9165 | 0.9167 | 0.9169 | 0.9173 | 0.9177 | 0.9182 | 0.9188 | 0.9195 | 0.9203 | 0.9214 | 0.9225 | 0.9233 | 0.9239 |
| 12 | 56726340 | rs74496027 | 56740682 | 2.9347 | -0.00596 | 0.9075 | 0.9075 | 0.9081 | 0.9105 | 0.9156 | 0.9204 | 0.9249 | 0.9292 | 0.9338 | 0.9367 | 0.9383 | 0.9403 | 0.9423 | 0.9434 | 0.9443 | 0.9458 | 0.9472 | 0.9487 | 0.9503 | 0.9519 | 0.9526 | 0.9528 | 0.9528 | 0.953 | 0.9535 | 0.9541 | 0.9546 | 0.9552 | 0.9558 | 0.9564 | 0.9568 | 0.9573 | 0.9579 | 0.9586 | 0.9595 | 0.9605 | 0.9617 | 0.963 | 0.9646 | 0.9664 | 0.9677 | 0.9686 | 0.9696 | 0.9704 | 0.9712 | 0.9721 | 0.973 | 0.974 | 0.9752 | 0.9765 |
| 12 | 56726340 | rs74496027 | 56740682 | 3.4366 | -0.00752 | 0.908 | 0.908 | 0.9087 | 0.9104 | 0.9128 | 0.9169 | 0.9197 | 0.9228 | 0.9249 | 0.9264 | 0.9279 | 0.9295 | 0.9297 | 0.9297 | 0.9298 | 0.9296 | 0.9291 | 0.9287 | 0.9283 | 0.928 | 0.9279 | 0.9281 | 0.9284 | 0.9285 | 0.9285 | 0.9285 | 0.9286 | 0.9289 | 0.9294 | 0.9302 | 0.9313 | 0.9324 | 0.9335 | 0.9347 | 0.9358 | 0.9367 | 0.9379 | 0.9391 | 0.9403 | 0.9414 | 0.9424 | 0.9433 | 0.9442 | 0.9453 | 0.9465 | 0.9479 | 0.9494 | 0.9511 | 0.9531 | 0.9553 |
| 12 | 56750204 | rs2066819 | na | 3.5288 | 0.00723 | 0.0908 | 0.0908 | 0.0902 | 0.0889 | 0.0879 | 0.0867 | 0.0857 | 0.0827 | 0.0799 | 0.078 | 0.0767 | 0.0754 | 0.0737 | 0.0719 | 0.0701 | 0.0691 | 0.0676 | 0.067 | 0.0666 | 0.0662 | 0.0663 | 0.0662 | 0.0659 | 0.0656 | 0.065 | 0.0642 | 0.0633 | 0.0622 | 0.0609 | 0.0597 | 0.0582 | 0.0566 | 0.055 | 0.0531 | 0.051 | 0.0493 | 0.0478 | 0.0464 | 0.0449 | 0.0435 | 0.0424 | 0.0413 | 0.0401 | 0.039 | 0.038 | 0.0368 | 0.0355 | 0.0341 | 0.0326 | 0.0309 |
| 12 | 56750204 | rs2066819 | na | 6.0687 | 0.00986 | 0.094 | 0.094 | 0.0926 | 0.0891 | 0.0865 | 0.082 | 0.0747 | 0.069 | 0.064 | 0.0594 | 0.0534 | 0.0475 | 0.0437 | 0.042 | 0.0405 | 0.0387 | 0.0368 | 0.0348 | 0.0321 | 0.0294 | 0.0273 | 0.0254 | 0.0238 | 0.0225 | 0.0212 | 0.02 | 0.0191 | 0.0183 | 0.0173 | 0.0163 | 0.0152 | 0.0144 | 0.0135 | 0.0125 | 0.0115 | 0.0107 | 0.0099 | 0.009 | 0.0081 | 0.0071 | 0.0059 | 0.0047 | 0.0033 | 0.0022 | 0.0016 | 0.0011 | 0.0008 | 0.0005 | 0.0003 | 0.0001 |
| 15 | 85207825 | rs12908161 | na | 0.6149 | -0.00144 | 0.2324 | 0.2324 | 0.2328 | 0.2363 | 0.2439 | 0.2492 | 0.2541 | 0.2602 | 0.2662 | 0.2704 | 0.2744 | 0.2788 | 0.2827 | 0.2881 | 0.2922 | 0.2958 | 0.2986 | 0.3007 | 0.302 | 0.3032 | 0.3042 | 0.3054 | 0.3072 | 0.3092 | 0.3111 | 0.3129 | 0.3149 | 0.3168 | 0.3189 | 0.3216 | 0.3246 | 0.3273 | 0.3303 | 0.3335 | 0.3367 | 0.3396 | 0.3423 | 0.3445 | 0.3466 | 0.3487 | 0.3512 | 0.3531 | 0.3549 | 0.3568 | 0.3585 | 0.3599 | 0.3616 | 0.3633 | 0.3651 | 0.3673 |
| 15 | 85207825 | rs12908161 | na | 0.069 | 0.00049 | 0.2347 | 0.2347 | 0.2351 | 0.2392 | 0.2438 | 0.2498 | 0.2547 | 0.2567 | 0.2607 | 0.2638 | 0.2663 | 0.2684 | 0.2706 | 0.2721 | 0.2728 | 0.2725 | 0.2703 | 0.267 | 0.2646 | 0.2634 | 0.263 | 0.2623 | 0.2619 | 0.2609 | 0.2595 | 0.2581 | 0.2576 | 0.2577 | 0.2574 | 0.2569 | 0.2563 | 0.2555 | 0.2547 | 0.2541 | 0.2534 | 0.2529 | 0.2526 | 0.2519 | 0.2511 | 0.2508 | 0.2509 | 0.2506 | 0.25 | 0.2493 | 0.2487 | 0.2478 | 0.2467 | 0.2458 | 0.245 | 0.244 |
| 17 | 76259847 | rs11650665 | na | 0.1637 | 0.00078 | 0.776 | 0.776 | 0.7758 | 0.7739 | 0.7693 | 0.7645 | 0.7624 | 0.759 | 0.7567 | 0.7546 | 0.7527 | 0.7505 | 0.7507 | 0.7516 | 0.7515 | 0.7518 | 0.7533 | 0.7548 | 0.7562 | 0.7568 | 0.7565 | 0.7562 | 0.7564 | 0.7562 | 0.7558 | 0.7552 | 0.7546 | 0.7537 | 0.7529 | 0.7527 | 0.7527 | 0.7526 | 0.7528 | 0.7522 | 0.7513 | 0.7503 | 0.7494 | 0.7487 | 0.7483 | 0.7481 | 0.7478 | 0.7473 | 0.7463 | 0.7453 | 0.7441 | 0.7431 | 0.742 | 0.7408 | 0.7395 | 0.7382 |
| 17 | 76259847 | rs11650665 | na | 0.0378 | -0.00039 | 0.7779 | 0.7779 | 0.7778 | 0.7753 | 0.7721 | 0.7698 | 0.7702 | 0.7727 | 0.7739 | 0.7726 | 0.7708 | 0.7694 | 0.7683 | 0.7682 | 0.7688 | 0.7699 | 0.7693 | 0.7682 | 0.7676 | 0.7669 | 0.7663 | 0.7663 | 0.7663 | 0.7662 | 0.7655 | 0.7651 | 0.7648 | 0.7647 | 0.7645 | 0.7642 | 0.764 | 0.7639 | 0.7635 | 0.7628 | 0.7621 | 0.7617 | 0.761 | 0.7607 | 0.7604 | 0.76 | 0.7594 | 0.759 | 0.7588 | 0.7589 | 0.7588 | 0.7587 | 0.7587 | 0.7589 | 0.7591 | 0.7591 |
| 18 | 74072245 | rs72973711 | na | 0.0038 | 0.0002 | 0.0942 | 0.0942 | 0.0943 | 0.0947 | 0.0953 | 0.097 | 0.0995 | 0.1022 | 0.1043 | 0.1067 | 0.108 | 0.1089 | 0.1087 | 0.1079 | 0.1086 | 0.1092 | 0.1099 | 0.1111 | 0.1116 | 0.112 | 0.1125 | 0.1129 | 0.1133 | 0.1135 | 0.1137 | 0.114 | 0.1145 | 0.1149 | 0.1158 | 0.1165 | 0.1173 | 0.1179 | 0.1183 | 0.1186 | 0.1186 | 0.1184 | 0.1182 | 0.1178 | 0.1181 | 0.1186 | 0.119 | 0.1192 | 0.1194 | 0.1195 | 0.1197 | 0.1196 | 0.1194 | 0.119 | 0.1185 | 0.1182 |
| 18 | 74072245 | rs72973711 | na | 0.0133 | -0.00032 | 0.0886 | 0.0886 | 0.0886 | 0.0892 | 0.0897 | 0.0904 | 0.0897 | 0.088 | 0.0885 | 0.0901 | 0.0923 | 0.0943 | 0.0948 | 0.094 | 0.0924 | 0.0912 | 0.09 | 0.0887 | 0.0878 | 0.0871 | 0.0871 | 0.0874 | 0.0874 | 0.0875 | 0.0876 | 0.0875 | 0.0879 | 0.0883 | 0.0886 | 0.0892 | 0.09 | 0.0907 | 0.0912 | 0.0917 | 0.0924 | 0.093 | 0.0936 | 0.094 | 0.0943 | 0.0947 | 0.0949 | 0.095 | 0.0949 | 0.0946 | 0.0945 | 0.0944 | 0.0941 | 0.0938 | 0.0936 | 0.0934 |
| 19 | 33390796 | rs10418340 | na | 0.0884 | 0.00083 | 0.0978 | 0.0978 | 0.098 | 0.099 | 0.1007 | 0.1024 | 0.1021 | 0.102 | 0.1033 | 0.1046 | 0.1044 | 0.1029 | 0.1021 | 0.1032 | 0.1046 | 0.106 | 0.1071 | 0.1087 | 0.111 | 0.113 | 0.1144 | 0.1157 | 0.1166 | 0.1173 | 0.1179 | 0.1184 | 0.119 | 0.1199 | 0.1207 | 0.1216 | 0.1222 | 0.1226 | 0.123 | 0.1231 | 0.1228 | 0.1225 | 0.1225 | 0.1223 | 0.1218 | 0.1211 | 0.1205 | 0.1199 | 0.119 | 0.1181 | 0.1171 | 0.1159 | 0.1142 | 0.1125 | 0.111 | 0.1099 |
| 19 | 33390796 | rs10418340 | na | 0.4418 | 0.00195 | 0.0934 | 0.0934 | 0.0932 | 0.0923 | 0.0902 | 0.0865 | 0.0845 | 0.0819 | 0.0796 | 0.0781 | 0.0779 | 0.0789 | 0.0795 | 0.0804 | 0.0804 | 0.0812 | 0.0826 | 0.0838 | 0.0854 | 0.0875 | 0.0894 | 0.0912 | 0.0927 | 0.0938 | 0.0949 | 0.096 | 0.0967 | 0.0976 | 0.0985 | 0.0993 | 0.0999 | 0.1005 | 0.1008 | 0.101 | 0.1009 | 0.1007 | 0.1004 | 0.0997 | 0.0988 | 0.0976 | 0.0965 | 0.0954 | 0.0941 | 0.093 | 0.0921 | 0.0913 | 0.0906 | 0.0898 | 0.0887 | 0.0875 |
| 20 | 4132364 | rs13043612 | na | 0.4219 | 0.00115 | 0.3921 | 0.3921 | 0.3923 | 0.3911 | 0.3858 | 0.38 | 0.3741 | 0.3713 | 0.3677 | 0.365 | 0.3653 | 0.3646 | 0.3629 | 0.3617 | 0.361 | 0.3604 | 0.3602 | 0.3606 | 0.3602 | 0.3601 | 0.3593 | 0.3577 | 0.3559 | 0.3541 | 0.3525 | 0.3515 | 0.3506 | 0.3496 | 0.3481 | 0.3463 | 0.3446 | 0.3432 | 0.3418 | 0.34 | 0.338 | 0.3362 | 0.3345 | 0.3326 | 0.3307 | 0.3284 | 0.3256 | 0.323 | 0.3209 | 0.3192 | 0.3177 | 0.3163 | 0.3148 | 0.3129 | 0.3112 | 0.3091 |
| 20 | 4132364 | rs13043612 | na | 0.452 | 0.0011 | 0.3907 | 0.3907 | 0.3903 | 0.3897 | 0.3874 | 0.3862 | 0.3859 | 0.3835 | 0.3823 | 0.3814 | 0.3794 | 0.3769 | 0.3752 | 0.3747 | 0.3725 | 0.3696 | 0.3655 | 0.3593 | 0.3531 | 0.3482 | 0.344 | 0.3415 | 0.3393 | 0.3375 | 0.3358 | 0.3347 | 0.3343 | 0.334 | 0.333 | 0.3323 | 0.3314 | 0.3306 | 0.3302 | 0.3299 | 0.3291 | 0.3278 | 0.3263 | 0.3248 | 0.3234 | 0.3218 | 0.3204 | 0.3185 | 0.3164 | 0.3146 | 0.3132 | 0.3114 | 0.3098 | 0.3082 | 0.3067 | 0.3051 |
| 20 | 16054982 | rs16997087 | na | 3.6372 | 0.00757 | 0.0687 | 0.0687 | 0.0682 | 0.0667 | 0.0652 | 0.0645 | 0.0644 | 0.0657 | 0.0673 | 0.0685 | 0.0698 | 0.0713 | 0.073 | 0.0741 | 0.0747 | 0.0757 | 0.0767 | 0.0772 | 0.0775 | 0.0772 | 0.0765 | 0.0753 | 0.0738 | 0.0729 | 0.072 | 0.071 | 0.0697 | 0.0687 | 0.068 | 0.0676 | 0.0672 | 0.0668 | 0.0662 | 0.0656 | 0.0651 | 0.0645 | 0.0639 | 0.0634 | 0.0628 | 0.0621 | 0.0612 | 0.0603 | 0.0592 | 0.058 | 0.0566 | 0.055 | 0.0531 | 0.0509 | 0.0486 | 0.0463 |
| 20 | 16054982 | rs16997087 | na | 0.285 | 0.00176 | 0.0693 | 0.0693 | 0.0692 | 0.0694 | 0.0711 | 0.0749 | 0.0778 | 0.0791 | 0.0795 | 0.0803 | 0.0819 | 0.0831 | 0.084 | 0.0846 | 0.0847 | 0.085 | 0.0861 | 0.0871 | 0.0877 | 0.0884 | 0.0887 | 0.0884 | 0.0875 | 0.0869 | 0.086 | 0.0856 | 0.0855 | 0.0851 | 0.0848 | 0.0842 | 0.0834 | 0.0824 | 0.0817 | 0.0811 | 0.0803 | 0.0796 | 0.0791 | 0.0786 | 0.0778 | 0.0772 | 0.0768 | 0.0763 | 0.0762 | 0.0761 | 0.0759 | 0.0755 | 0.075 | 0.0744 | 0.0736 | 0.0727 |

**Table S12: MPRA information for aSNPs with cis regulatory activity in STAT2 region**

| chr | position | rs | REF | ALT | CRE | emVar | Non-Introg LFCb | Introg LFCb | Expression Modulation LFCb | a-cTF \| trans-eQTL |
|---|---|---|---|---|---|---|---|---|---|---|
| 12 | 56638077 | rs2066807 | T | C | X | X | -0,28 | 0,53 | 0,81 | STAT2 \| rs2066807/rs2066819 |
| 12 | 56660905 | rs2066807 | G | T | X | X | 3,21 | 2,67 | -0,54 | STAT2 \| rs2066807/rs2066819 |
| 12 | 56727705 | rs2066807 | G | A | X | X | 1,36 | 0,71 | -0,65 | STAT2 \| rs2066807/rs2066819 |
| 3 | 46184620 | rs13083881 | C | G | X | X | x | -0,33 | -0,53 | - \| rs13063635/rs13098911 |
| 3 | 46250008 | rs34919616 | G | A | X | X | 1,61 | 1,20 | -0,36 | - \| rs13063635/rs13098911 |
| 3 | 46272440 | rs1542755 | G | T | X | X | 1,90 | 0,32 | -1,53 | - \| rs13063635/rs13098911 |
